# The European Reference Genome Atlas: piloting a decentralised approach to equitable biodiversity genomics

**DOI:** 10.1101/2023.09.25.559365

**Authors:** Ann M Mc Cartney, Giulio Formenti, Alice Mouton, Diego De Panis, Luisa S Marins, Henrique G Leitao, Genevieve Diedericks, Joseph Kirangwa, Marco Morselli, Judit Salces, Nuria Escudero, Alessio Iannucci, Chiara Natali, Hannes Svardal, Rosa Fernandez, Tim De Pooter, Geert Joris, Mojca Strazisar, Jo Wood, Katie E Herron, Ole Seehausen, Phillip C Watts, Felix Shaw, Robert P Davey, Alice Minotto, Jose Maria Fernandez Gonzalez, Astrid Bohne, Carla Alegria, Tyler Alioto, Paulo C Alves, Isabel R Amorim, Jean-Marc Aury, Niclas Backstrom, Petr Baldrian, Loriano Ballarin, Laima Baltrunaite, Endre Barta, Bertrand BedHom, Caroline Belser, Johannes Bergsten, Laurie Bertrand, Helena Bilandija, Mahesh Binzer-Panchal, Iliana Bista, Mark Blaxter, Paulo AV Borges, Guilherme Borges Dias, Mirte Bosse, Tom Brown, Remy Bruggmann, Elena Buena-Atienza, Josephine Burgin, Elena Buzan, Alessia Cariani, Nicolas Casadei, Matteo Chiara, Sergio Chozas, Fedor Ciampor, Angelica Crottini, Corinne Cruaud, Fernando Cruz, Love Dalen, Alessio De Biase, Javier del Campo, Teo Delic, Alice B Dennis, Martijn FL Derks, Maria Angela Diroma, Mihajla Djan, Simone Duprat, Klara Eleftheriadi, Philine GD Feulner, Jean-Francois Flot, Giobbe Forni, Bruno Fosso, Pascal Fournier, Christine Fournier-Chambrillon, Toni Gabaldon, Shilpa Garg, Carmela Gissi, Luca Giupponi, Jessica Gomez-Garrido, Josefa Gonzalez, Miguel L Grilo, Bjoern Gruening, Thomas Guerin, Nadege Guiglielmoni, Marta Gut, Marcel P Haesler, Christoph Hahn, Balint Halpern, Peter Harrison, Julia Heintz, Maris Hindrikson, Jacob Hoglund, Kerstin Howe, Graham Hughes, Benjamin Istace, Mark J. Cock, Franc Jancekovic, Zophonias O Jonsson, Sagane Joye-Dind, Janne J. Koskimaki, Boris Krystufek, Justyna Kubacka, Heiner Kuhl, Szilvia Kusza, Karine Labadie, Meri Lahteenaro, Henrik Lantz, Anton Lavrinienko, Lucas Leclere, Ricardo Jorge Lopes, Ole Madsen, Ghislaine Magdelenat, Giulia Magoga, Tereza Manousaki, Tapio Mappes, Joao Pedro Marques, Gemma I Martinez Redondo, Florian Maumus, Shane A. McCarthy, Hendrik-Jan Megens, Jose Melo-Ferreira, Sofia L Mendes, Matteo Montagna, Joao Moreno, Mai-Britt Mosbech, Monica Moura, Zuzana Musilova, Eugene Myers, Will J. Nash, Alexander Nater, Pamela Nicholson, Manuel Niell, Reindert Nijland, Benjamin Noel, Karin Noren, Pedro H Oliveira, Remi-Andre Olsen, Lino Ometto, Rebekah A Oomen, Stephan Ossowski, Vaidas Palinauskas, Snaebjorn Palsson, Jerome P Panibe, Joana Pauperio, Martina Pavlek, Emilie Payen, Julia Pawlowska, Jaume Pellicer, Graziano Pesole, Joao Pimenta, Martin Pippel, Anna Maria Pirttila, Nikos Poulakakis, Jeena Rajan, Ruben MC Rego, Roberto Resendes, Philipp Resl, Ana Riesgo, Patrik Rodin-Morch, Andre ER Soares, Carlos Rodriguez Fernandes, Maria M. Romeiras, Guilherme Roxo, Lukas Ruber, Maria Jose Ruiz-Lopez, Urmas Saarma, Luis P Silva, Manuela Sim-Sim, Lucile Soler, Vitor C Sousa, Carla Sousa Santos, Alberto Spada, Milomir Stefanovic, Viktor Steger, Josefin Stiller, Matthias Stock, Torsten Hugo H Struck, Hiranya Sudasinghe, Riikka Tapanainen, Christian Tellgren-Roth, Helena Trindade, Yevhen Tukalenko, Ilenia Urso, Benoit Vacherie, Steven M Van Belleghem, Kees van Oers, Carlos Vargas-Chavez, Nevena Velickovic, Noel Vella, Adriana Vella, Cristiano Vernesi, Sara Vicente, Sara Villa, Olga Vinnere Pettersson, Filip AM Volckaert, Judit Voros, Patrick Wincker, Sylke Winkler, Claudio Ciofi, Robert M Waterhouse, Camila J Mazzoni

**Affiliations:** Genomics Institute, University of California, Santa Cruz, CA 95060 USA; The Vertebrate Genome Laboratory, The Rockefeller University, 1240 York Ave 10065 New York; Department of Biology, University of Florence, Sesto Fiorentino, Italy; InBios-Conservation Genetics Laboratory, University of Liege, Chemin de la Vallee 4, 4000 Liege, Belgium;Department of Biology, University of Florence, 50019 Sesto Fiorentino (FI), Italy;SEED-Departement des sciences et gestion de l'environnement, Univers; Leibniz Institut fur Zoo und Wildtierforschung, Berlin, Germany;Berlin Center for Genomics in Biodiversity Research, Berlin, Germany; Leibniz Institut fur Zoo und Wildtierforschung, Berlin, Germany; Department of Biology, University of Antwerp, 2610, Antwerp, Belgium; Institute of Zoology, University of Cologne, Zulpicher Strasse 47b, 50674 Cologne, Germany; Department of Chemistry, Life Sciences and Environmental Sustainability, University of Parma, Parco Area delle Scienze 23A, 43124 Parma, Italy; Metazoa Phylogenomics Lab, Institute of Evolutionary Biology, Spain; Metazoa Phylogenomics Lab, Institute of Evolutionary Biology, Barcelona, Spain; Department of Biology, University of Florence; University of Florence; Department of Biology, University of Antwerp, 2610 Antwerp, Belgium;Naturalis Biodiversity Center, Leiden, The Netherlands; Metazoa Phylogenomics Lab, Institute of Evolutionary Biology (CSIC-Universitat Pompeu Fabra), Spain; Neuromics Support Facility, VIB Center for Molecular Neurology, VIB, Antwerp, Belgium;Neuromics Support Facility, Department of Biomedical Sciences, University of Antwerp, Antwerp, Belgium; Tree of Life, Wellcome Sanger Institute, Hinxton, Cambridge, UK, CB10 1SA; School of Biology and Environmental Science, University College Dublin, Belfield, D04 V1W8 Dublin, Ireland; Institute of Ecology & Evolution, University of Bern, Baltzerstrasse 6, 3012 Bern, Switzerland;Department Fish Ecology & Evolution, Eawag, Center for Ecology, Evolution and Biogeochemistry, Seestrasse 79, 6047 Kastanienbaum, Switzerland; Department of Biological and Environmental Science, University of Jyvaskyla, Jyvaskyla, Finland, 40014; Earlham Institute, Norwich Research Park, Colney Lane, Norwich, NR4 7UZ; Digital Science, London; Barcelona Supercomputing Center;Spanish National Bioinformatics Institute, ELIXIR Spain; Leibniz Institute for the Analysis of Biodiversity Change, Museum Koenig Bonn; cE3c Centre for Ecology, Evolution and Environmental Changes, CHANGE Global Change and Sustainability Institute, Faculdade de Ciencias, Universidade de Lisboa, 1749 016 Lisbon, Portugal; Centro Nacional de Analisis Genomico, C Baldiri Reixac 4, 08028 Barcelona, Spain; CIBIO, Centro de Investigacao em Biodiversidade e Recursos Geneticos, InBIO Laboratorio Associado, Universidade do Porto, Vairao, Portugal;Departamento de Biologia, Faculdade de Ciencias, Universidade do Porto, 4099 002 Porto, Portugal; BIOPOLIS Program i; Centre for Ecology, Evolution and Environmental Changes, Azorean Biodiversity Group, CHANGE Global Change and Sustainability Institute, and University of the Azores, Rua Capitao Joao dAvila, Pico da Urze, 9700 042, Angra do Heroismo, Portugal; Genomique Metabolique, Genoscope, Institut Francois Jacob, CEA, CNRS, Univ Evry, Universite Paris Saclay, Evry, 91057, France; Evolutionary Biology Program, Department of Ecology and Genetics, Uppsala University, Sweden; Institute of Microbiology of the Czech Academy of Sciences, Videnska 1083, 14200, Praha 4, Czech Republic; University of Padova, Department of Biology, Italy; Nature Research Centre; Institute of Biochemistry and Molecular Biology, Faculty of Medicine, University of Debrecen, Debrecen, Hungary; Institut de Systematique, Evolution, Biodiversite, UMR7205, Museum National d Histoire Naturelle, CNRS, Sorbonne Universite, EPHE, Universite des Antilles, Paris, France; Department of Zoology, Swedish Museum of Natural History, Box 50007, SE10405 Stockholm, Sweden;Department of Zoology, Faculty of Science, Stockholm University, SE106 91 Stockholm, Sweden; Genoscope, Institut Francois Jacob, Commissariat a lEnergie Atomique, Universite Paris Saclay, 2 Rue Gaston Cremieux, 91057 Evry, France.; Ruder Boskovic Institute; National Bioinformatics Infrastructure Sweden, Science for Life Laboratory, Sweden.;Uppsala University; Senckenberg Research Institute, 60325 Frankfurt, Germany;LOEWE Centre for Translational Biodiversity Genomics, 60325 Frankfurt, Germany;Wellcome CRUK Gurdon Institute, University of Cambridge, Tennis Court Rd, Cambridge, CB2 1QN, UK; Tree of Life, Wellcome Sanger Institute, Cambridge, UK; cE3c Centre for Ecology, Evolution and Environmental Changes Azorean Biodiversity Group, CHANGE Global Change and Sustainability Institute, School of Agricultural and Environmental Sciences, University of the Azores, Rua Capitao Joao d Avila, Pico da Urze; National Bioinformatics Infrastructure Sweden, Science for Life Laboratory, Sweden.;Department of Cell and Molecular Biology, Uppsala University, Uppsala, Sweden.; VU University Amsterdam, Amsterdam, The Netherlands;Wageningen University & Research, Wageningen, The Netherlands; Max Planck Institute of Molecular Cell Biology and Genetics;Leibniz Institute for Zoo and Wildlife Research; Interfaculty Bioinformatics Unit and Swiss Institute of Bioinformatics, University of Bern, Bern, Switzerland; Institute of Medical Genetics and Applied Genomics;NGS Competence Center Tubingen; European Molecular Biology Laboratory, European Bioinformatics Institute, Wellcome Genome Campus, Hinxton, Cambridge CB10 1SD, UK; University of Primorska, Faculty of Mathematics, Natural Sciences, and Information Technologies, Glagoljaska 8, 6000 Koper, Slovenia;Faculty of Environmental Protection, Trg mladosti 7, 3320 Velenje, Slovenia; Department of Biological, Geological and Environmental Sciences, Alma Mater Studiorum Universita di Bologna; Institute of Medical Genetics and Applied Genomics, University of Tubingen;NGS Competence Center Tubingen, Tubingen, Germany; Dipartimento di Bioscienze, Universita degli Studi di Milano;Institute of Biomembranes, Bioenergetics and Molecular Biotechnologies of the National Research Council in Bari.; cE3c Centre for Ecology, Evolution and Environmental Changes, CHANGE Global Change and Sustainability Institute, Sciences Faculty of the University of Lisbon;Sociedade Portuguesa de Botanica, Lisbon, Portugal; Plant Scicence and Biodiversity Centre SAS, Dubravska cesta 9, 84523, Bratislava, SLOVAKIA; CIBIO, Centro de Investigacao em Biodiversidade e Recursos Geneticos, InBIO Laboratorio Associado, Campus de Vairao, Universidade do Porto, 4485 661 Vairao, Portugal;Departamento de Biologia, Faculdade de Ciencias, Universidade do Porto, rua do Campo Aleg; Genoscope, Institut Francois Jacob, Commissariat a l'Energie Atomique, Universite Paris Saclay, 2 Rue Gaston Cremieux, 91057 Evry, France; Centro Nacional de AnAlisis Genomico, C Baldiri Reixac 4, 08028 Barcelona, Spain; Department of Zoology, Stockholm University;Department of Bioinformatics and Genetics, Swedish Museum of Natural History;Centre for Palaeogenetics, Stockholm; Department of Biology and Biotechnologies;Sapienza University of Rome; Institut de Biologia Evolutiva; University of Ljubljana, Biotechnical Faculty, Department of Biology, SubBio Lab; University of Namur, Belgium; Wageningen University & Research, Netherland; Department of Biology, University of Florence,Italy; Department of Biology and Ecology, University of Novi Sad, Serbia; Metazoa Phylogenomics Lab, Biodiversity Program, Institute of Evolutionary Biology, Passeig maritim de la Barceloneta 37-49, 08003 Barcelona, Spain; Eawag Swiss Federal Institute of Aquatic Science and Technology Center for Ecology, Evolution and Biogeochemistry, Department of Fish Ecology & Evolution, Seestrasse 79, 6047 Kastanienbaum, Switzerland;University of Bern Institute of Ecology & Evolution; Department of Organismal Biology, Universite libre de Bruxelles, Brussels, Belgium; University of Bologna, Italy; Universita degli Studi di Bari A. Moro Dipartimento di Bioscienze, Biotecnologie e Ambiente; Groupe de Recherche et d Etude pour la Gestion de l Environnement, 1 La Peyrere, 33730 VILLANDRAUT; Barcelona Supercomputing Centre; Institut for Research in Biomedicine;CIBERINFEC; NNF Center for Biosustainability, Technical University of Denmark, Denmark; Department of Biosciences, Biotechnology and Environment, University of Bari Aldo Moro, Bari, Italy;Institute of Biomembranes, Bioenergetics and Molecular Biotechnologies, Consiglio Nazionale delle Ricerche, Bari, Italy;CoNISMa, Consorzio Nazionale Interu; Centre of Applied Studies for the Sustainable Management and Protection of Mountain Areas CRC Ge.S.Di.Mont., University of Milan, 25048 Edolo, Italy;Department of Agricultural and Environmental Sciences-Production, Landscape and Agroenergy DiSAA, Universi; Institute of Evolutionary Biology, CSIC, UPF; Marine and Environmental Sciences Centre, Aquatic Research Network, Instituto UniversitArio de Ciencias Psicologicas, Sociais e da Vida, Lisboa, Portugal;Egas Moniz Center for Interdisciplinary Research (CiiEM), Egas Moniz School of Health & Science, Capa; University of Freiburg; Genoscope, Institut Francois Jacob, Commissariat a l'Energie Atomique, Universite Paris Saclay, 2 Rue Gaston Cremieux, 91057 Evry, France.; Universitat zu Koln, Zulpicher str. 47b, 50674 Cologne, Germany; Centro Nacional de Analisis Genomico, C Baldiri Reixac 4, 08028 Barcelona, Spain.; Aquatic Ecology & Evolution, Institute of Ecology & Evolution, University of Bern, Switzerland;Departement of Fish Ecology & Evolution, Eawag, Kastanienbaum, Switzerland; Institute of Biology, University of Graz, Austria; MME BirdLife Hungary, Budapest, Hungary;Integrative Ecology ResearDoctoral School of Biology, Department of Systematic Zoology and Ecology, Institute of Biology, ELTE Eotvos LorAnd University, Budapest, Hungary; Integrative Ecology Research Group, Budapes; European Molecular Biology Laboratory - European Bioinformatics Institute; SciLifeLab Genomics NGI, Uppsala Genome Center;Uppsala University; University of Tartu, Department of Zoology Chair of Mammalogy, estonia; Dept of Ecology and Genetics,Uppsala University, Sweden; School of Biology and Environmental Science, University College Dublin, Belfield, Dublin 4, Ireland;UCD Conway Institute, University College Dublin, Belfield, Dublin 4, Ireland; Algal Genetics Group, UMR 8227, CNRS, Sorbonne Universite, UPMC University Paris 06, Paris, France Integrative Biology of Marine Models, Station Biologique de Roscoff, CS 90074, F29688 Roscoff, France; University of Maribor, Faculty of Natural Sciences and Mathematics, Koroska cesta 160, 2000, Maribor, Slovenia; Institute of Life and Environmental Sciences, University of Iceland, Reykjavik, Iceland; University of Lausanne;Swiss Institute of Bioinformatics; Ecology and Genetics Research Unit, University of Oulu, FI 90014, P.O. Box 3000, Oulu, Finland; Slovenian Museum of Natural History, Ljubljana, Slovenia;Science and Research Centre Koper, Koper, Slovenia; Museum and Institute of Zoology, Polish Academy of Sciences; Leibniz Institute of freshwater ecology and inland fisheries, Berlin, DE; University of Debrecen; Department of Zoology, Swedish Museum of Natural History, Box 50007, SE 10405 Stockholm, Sweden;Department of Zoology, Faculty of Science, Stockholm University, SE 106 91 Stockholm, Sweden; Uppsala University;SciLifeLab;National Bioinformatics Infrastructure Sweden; Laboratory of Food Systems Biotechnology, Institute of Food, Nutrition, and Health, ETH Zurich, Zurich 8092, Switzerland; Sorbonne Universite, CNRS, Biologie Integrative des Organismes Marins, BIOM, Banyuls-sur-Mer, France;Sorbonne Universite, CNRS, Laboratoire de Biologie du Developpement de Villefranche-sur-Mer (LBDV), Villefranche-sur-Mer, France; CIBIO, Centro de Investigacao em Biodiversidade e Recursos Geneticos, InBIO Laboratorio Associado, Campus de Vairao, Universidade do Porto, Vairao, Portugal;BIOPOLIS Program in Genomics, Biodiversity and Land Planning, CIBIO, Vairao, Portugal;MHNC-UP, Nat; Animal Breeding & Genomics, Wageningen University & Research, PO Box 338, 6700 AH Wageningen, The Netherlands; Genoscope, Institut Francois Jacob, Commissariat a l'Energie Atomique (CEA), Universite Paris Saclay, 2 Rue Gaston Cremieux, 91057 Evry, France.; Department of Agricultural Sciences, University of Naples Federico II, Via Universita 100, 80055 Portici, Italy; Hellenic Centre for Marine Research (HCMR), Institute of Marine Biology, Biotechnology and Aquaculture (IMBBC), Heraklion, Crete, Greece; Department of Biological and Environmental Science. University of Jyvaskyla; CIBIO, Centro de Investigacao em Biodiversidade e Recursos Geneticos, InBIO Laboratorio Associado, Campus de Vairao, Universidade do Porto, 4485-661 Vairao, Portugal;BIOPOLIS Program in Genomics, Biodiversity and Land Planning, CIBIO, Campus de Vairao, 44; Universite Paris Saclay, INRAE, URGI, 78026, Versailles, France; Department of Genetics, University of Cambridge, Cambridge, UK; Wellcome Sanger Institute, Cambridge, UK; Animal Breeding and Genomics, Wageningen University and Research; CIBIO, Centro de Investigacao em Biodiversidade e Recursos Geneticos, InBIO Laboratorio Associado, Universidade do Porto, Vairao, Portugal;Departamento de Biologia, Faculdade de Ciencias da Universidade do Porto, Porto, Portugal;BIOPOLIS Program in Genomi; cE3c Centre for Ecology, Evolution and Environmental Changes & CHANGE Global Change and Sustainability Institute, Faculdade de Ciencias, Universidade de Lisboa, Campo Grande, 1749-016 Lisboa, Portugal; Department of Agricultural Sciences, University of Naples Federico II, Portici, Italy.;Interuniversity Center for Studies on Bioinspired Agro Environmental Technology, University of Naples Federico II, Portici, Italy.; cE3c Centre for Ecology, Evolution and Environmental Changes & CHANGE Global Change and Sustainability Institute, Faculdade de Ciencias da Universidade de Lisboa, Lisboa, Portugal;MARE/ULisboa Centro de Ciencias do Mar e do Ambiente & ARNET Aquatic Resear; SciLifeLab Genomics NGI, Uppsala Genome Center; Faculty of Sciences and Technology, University of the Azores, Campus of Ponta Delgada, Rua Mae de Deus 13A, 9500 321 Ponta Delgada, Azores, Portugal;Research Centre in Biodiversity and Genetic Resources, InBIO Associate Laboratory; BIOPOLIS Program in Gen; Department of Zoology, Faculty of Science, Charles University, Prague, Czech Republic; Max Planck Institute of Molecular Cell Biology and Genetics; The Earlham Institute, Norwich Research Park, Norwich, NR4 7UZ, UK; Next Generation Sequencing Platform, University of Bern; Andorra Research and Innovation;Catalan initiative for the Earth Biogenome Project; Marine Animal Ecology group, Wageningen University and Research, P.O. Box 338, 6700 AH, Wageningen, The Netherlands; Department of Zoology, Stockholm University; Genomique Metabolique, Genoscope, Institut Francois Jacob, CEA, CNRS, Universite Evry, Universite ParisSaclay, 91057 Evry, France; Science for Life Laboratory, Department of Biochemistry and Biophysics, Stockholm University, 17165 Solna, Sweden; Department of Biology and Biotechnology, University of Pavia, Pavia, Italy;National Biodiversity Future Center, Italy; Centre for Ecological and Evolutionary Synthesis, University of Oslo, Oslo, Norway; University of New Brunswick Saint John, Saint John, New Brunswick, Canada; Institute for Medical Genetics and Applied Genomics, University of Tubingen, Tubingen, Germany;NGS Competence Center Tubingen (NCCT), University of Tubingen, Tubingen, Germany; Institute for Bioinformatics and Medical Informatics (IBMI), University of Tubi; Nature Research Centre, Akademijos 2, LT-09412 Vilnius, Lithuania; Faculty of Life and Environmental Sciences, University of Iceland; Biodiversity Research Center, Academia Sinica; European Molecular Biology Laboratory, European Bioinformatics Institute, Wellcome Genome Campus, Hinxton, CB10 1SD, UK; Ruder Boskovic Institute, Bijenicka cesta 54, 10000 Zagreb; Faculty of Biology, University of Warsaw, Warsaw, Poland; Institut Botanic de Barcelona, Passeig del Migdia s.n., Parc de Montjuic, 08038 Barcelona, Spain; University of Bari Aldo Moro, Department of Biosciences, Biotechnology and Environment; Consiglio Nazionale delle Ricerche, Institute of Biomembranes. Bioenergetics and Molecular Biotechnologies; CIBIO, Centro de Investigacao em Biodiversidade e Recursos Geneticos, InBIO Laboratorio Associado, Universidade do Porto; Vairao, Portugal;BIOPOLIS Program in Genomics, Biodiversity and Land Planning, CIBIO, Vairao, Portugal; National Bioinformatics Infrastructure Sweden;Department of Cell and Molecular Biology, Uppsala university; Ecology and Genetics Research Unit, University of Oulu, PO Box 3000, Linnanmaa, FIN 90014 Oulu, Finland; Department of Biology, School of Sciences and Engineering, University of Crete, Voutes University Campus, 70013 Irakleio, Greece;Natural History Museum of Crete, School of Sciences and Engineering, University of Crete, Knosos Avenue, 71409, Irakleio, Gree; European Molecular Biology Laboratory, European Bioinformatics Institute, Wellcome Genome Campus, Hinxton, Cambridge, United Kingdom; CIBIO, Centro de Investigacao em Biodiversidade e Recursos Geneticos, InBIO Laboratorio Associado, Polo dos Acores Faculdade de Ciencias e Tecnologia da Universidade dos Acores, Ponta Delgada, Portugal;BIOPOLIS Program in Genomics, Biodiversity and Land P; Universidade dos Acores, Departamento de Biologia, Ponta Delgada; University of Graz, Institute of Biology, Universitatsplatz 2, 8010 Graz, Austria; Department of Biodiversity and Evolutionary Biology, Museo Nacional de Ciencias Naturales; Department of Ecology and Genetics, Uppsala University; Uppsala University, Department of Medical Biochemistry and Microbiology, Uppsala, Sweden;National Bioinformatics Infrastructure Sweden, Science for Life Laboratory, Sweden; cE3c Centre for Ecology, Evolution and Environmental Changes, CHANGE Global Change and Sustainability Institute, Faculdade de Ciencias, Universidade de Lisboa, Campo Grande, 1749 016 Lisboa, Portugal;Faculdade de Psicologia, Universidade de Lisboa, Alame; Linking Landscape, Environment, Agriculture and Food, Associated Laboratory TERRA, Instituto Superior de Agronomia, Universidade de Lisboa, Tapada da Ajuda, 1349 017 Lisboa, Portugal;Centre for Ecology, Evolution and Environmental Changes, Global Change a; University of the Azores; Naturhistorisches Museum Bern, Bern, Switzerland;Division of Aquatic Ecology and Evolution, Institute of Ecology and Evolution, University of Bern, Bern, Switzerland; Department of Wetland Ecology EBD CSIC, Estacion Biologica de Donana, Avda. Americo Vespucio 26, E41092 Sevilla, Spain;CIBER of Epidemiology and Public Health, Spain; Department of Zoology, Institute of Ecology and Earth Sciences, University of Tartu, J. Liivi 2, 50409 Tartu, Estonia; CIBIO, Centro de Investigacao em Biodiversidade e Recursos Geneticos, InBIO Laboratorio Associado, Campus de Vairao, Universidade do Porto, 4485 661 Vairao, Portugal;BIOPOLIS Program in Genomics, Biodiversity and Land Planning, CIBIO, Campus de Vairao, 44; cE3c Centre for Ecology, Evolution and Environmental Changes, CHANGE Global Change and Sustainability Institute, Departamento de Biologia Vegetal, Faculdade de Ciencias, Universidade de Lisboa, Campo Grande, 1749 016 Lisboa, Portugal;Museu Nacional de His; Department of Medical Biochemistry and Microbiology, Uppsala University;Science for Life Laboratory, SciLifeLab;National Bioinformatics Infrastructure Sweden, Uppsala, Sweden; Centre for Ecology, Evolution and Environmental Changes, CHANGE Global Change and Sustainability Institute, Departamento de Biologia Animal, Faculdade de Ciencias da Universidade de Lisboa, Lisboa, Portugal; MARE Marine and Environmental Sciences Centre, ARNET Aquatic Research Network; Department of Agricultural and Environmental Sciences Production, Landscape, Agroenergy, University of Milan, Via Celoria 2, 20133 Milan, Italy; Department of Genetics and Genomics, Institute of Genetics and Biotechnology, Hungarian University of Agriculture and Life Sciences, Godollo, Hungary; Section for Ecology and Evolution, Department of Biology, University of Copenhagen, Denmark; Leibniz-Institute of Freshwater Ecology and Inland Fisheries, Muggelseedamm 301, D-12587 Berlin, Germany; Natural History Museum, University of Oslo, P.O. Box 1172, Blindern, 0318 Oslo, Norway; Naturhistorisches Museum Bern, Bernastrasse 15, 3005 Bern, Switzerland;Evolutionary Ecology, Institute of Ecology and Evolution, University of Bern, 3012 Bern, Switzerland; University of Eastern Finland; Centre for Ecology, Evolution and Environmental Change, Faculty of Sciences, University of Lisbon, Global Change and Sustainability Institute, 1749-016 Lisboa, Portugal; Institute for Nuclear Research of the NAS of Ukraine; CNR IBIOM, Italy; Ecology, Evolution and Conservation Biology, Department of Biology, KU Leuven, Leuven, Belgium; Department of Animal Ecology, Netherlands Institute of Ecology, Wageningen, the Netherlands; Metazoa Phylogenomics Lab, Biodiversity Program, Institute of Evolutionary Biology, Passeig maritim de la Barceloneta 37 49, 08003 Barcelona, Spain; Conservation Biology Research Group, Department of Biology, University of Malta, Msida, Malta; Foresto Ecology Unit, Research and Innovation Centre-Fondazione Edmund Mach; cE3c Centre for Ecology, Evolution and Environmental Changes & CHANGE Global Change and Sustainability Institute, Departamento de Biologia Vegetal, Faculdade de Ciencias, Universidade de Lisboa, Campo Grande, 1749 016 Lisboa, Portugal;ERISA Escola Superio; Institute for Sustainable Plant Protection, National Research Council, via Madonna del Piano 10, 50019, Sesto Fiorentino, Italy;Department of Agricultural and Environmental Sciences, University of Milan, via Giovanni Celoria 2, 20133 Milan, Italy; SciLifeLab Genomics NGI, Uppsala Genome Center,Uppsala University; Laboratory of Biodiversity and Evolutionary Genomics, KU Leuven, B3000 Leuven, Belgium; Department of Zoology, Hungarian Natural History Museum; MPI of Molecular Cell Biology and Genetics, Dresden, Germany;DRESDEN concept Genome Center, Dresden, Germany; Department of Biology, University of Florence, 50019 Sesto Fiorentino, Italy; Department of Ecology and Evolution, University of Lausanne, Lausanne, Switzerland;Swiss Institute of Bioinformatics, Lausanne, Switzerland

## Abstract

A global genome database of all of Earth’s species diversity could be a treasure trove of scientific discoveries. However, regardless of the major advances in genome sequencing technologies, only a tiny fraction of species have genomic information available. To contribute to a more complete planetary genomic database, scientists and institutions across the world have united under the Earth BioGenome Project (EBP), which plans to sequence and assemble high-quality reference genomes for all ∼1.5 million recognized eukaryotic species through a stepwise phased approach. As the initiative transitions into Phase II, where 150,000 species are to be sequenced in just four years, worldwide participation in the project will be fundamental to success. As the European node of the EBP, the European Reference Genome Atlas (ERGA) seeks to implement a new decentralised, accessible, equitable and inclusive model for producing high-quality reference genomes, which will inform EBP as it scales. To embark on this mission, ERGA launched a Pilot Project to establish a network across Europe to develop and test the first infrastructure of its kind for the coordinated and distributed reference genome production on 98 European eukaryotic species from sample providers across 33 European countries. Here we outline the process and challenges faced during the development of a pilot infrastructure for the production of reference genome resources, and explore the effectiveness of this approach in terms of high-quality reference genome production, considering also equity and inclusion. The outcomes and lessons learned during this pilot provide a solid foundation for ERGA while offering key learnings to other transnational and national genomic resource projects.

## Background

### Reference genomes as a key biodiversity genomics tool

In the midst of the Earth’s sixth mass extinction, species worldwide are declining at an unprecedented rate^1^ directly impacting ecosystem functioning and services^2^, human health^3^ and our resilience to climate disturbances^4^. Biodiversity and ecosystem decline^5,6^, loss and degradation raise the prospect that much, if not most, of the Earth’s biodiversity will be lost forever before they can be genomically explored - analogous to the ‘dark extinctions’ in the pre-taxonomic period^7^. Our ability to genomically characterise and investigate the species that span the tree of life, and their ecosystems, can help not only scientifically inform decision making processes to flatten the biodiversity extinction curve^8^, but also can unlock diverse genetic-, species- and ecosystem-level^9^ discoveries that can be used for human health, bioeconomy stimulation, food sovereignty, biosecurity amongst many more.

As genomic sequencing has become increasingly cost effective and the platforms and computational algorithms become more technically efficient, many biodiversity genomics tools have become available to expedite the investigation of both known and unknown species e.g., DNA barcoding, genome skimming, reduced representation sequencing, transcriptome sequencing, and whole genome sequencing for reference genome production^10^. Reference genomes (**Glossary)** are one such tool that offer an unparalleled, scalable, and increasingly cost-effective high resolution insight into species, and their accessibility has made the construction of a planetary-wide genomic database of all eukaryotic life a more realistic endeavor^11^.

To date, reference genomes do not exist for most of eukaryotic life. For instance, the largest genomics data repository, the International Nucleotide Sequence Database Collaboration (INSDC), has genome-wide DNA sequence information for just 6,480 eukaryotic species (about 0.43% of described species) of which over 63% (4,082) are short-read based (draft quality)^11^ and most are variable in terms of sequence quality, data type, data volume, associated voucher samples, completeness of metadata and protocol reproducibility^12–14^. Building from this, the biodiversity research community is pushing to expand beyond reference genome production alone and toward the production of a complete reference resource for each species. A complete reference resource includes a reference genome, an annotation, all metadata, and associated *ex-situ* samples (voucher(s) and cryopreserved specimen(s)). Complete reference resources are necessary to unlock the plurality of possible scientific enquiries beyond the scope of any singular research project^9^. However, the scientific enquiries that can be realised from reference resources^15^ are limited in scope due in large to a current lack of standardisation across the multitude of actors involved throughout the production of complete reference resources.

### The state of reference genome production today

After two decades of uncoordinated and unstandardised biodiversity genomics sequencing data production (e.g. with little coordination among individual research laboratories or projects), the Earth BioGenome Project (EBP)^11^ was established. The goal of the EBP is to create a global network of biodiversity genomics researchers that share a mission to produce a database of openly accessible, standardised, and complete reference resources that span the whole eukaryotic phylogenetic tree. The project has a three-phase approach and to date (Phase I) has produced ∼1,213 reference genomes for species across ∼1,010 genera^16^. However the rate of production is fast increasing, for instance in 2022 over 316 reference genomes were produced and in the coming years the rate is estimated to increase by at least 10 fold. It is important to acknowledge that during this initial phase, 910 reference genomes were produced by a single affiliated project, the Darwin Tree of Life^17^, and a further 120 by the Wellcome Sanger Tree of Life Programme (https://www.sanger.ac.uk/programme/tree-of-life/). As the EBP approaches Phase II where 150,000 reference resources for species are planned, the status quo centralised approach poses significant challenges for scaling up reference genome production. Additionally, it raises important concerns regarding inclusion, accessibility, equity, and fairness.

### The goal of building a decentralised model embracing all of Europe and beyond

Given these limitations, the European node of the EBP, the European Reference Genome Atlas (ERGA) **(Box 1)** set out to develop and implement a pilot decentralised infrastructure that would act to test the effectiveness of the approach in creating and scaling reference genomic resources for Europe’s eukaryotes.

##### Box 1: The European Reference Genome Atlas

As the European node of the Earth BioGenome Project (EBP; https://www.earthbiogenome.org/)^11^, the mission of the European Reference Genome Atlas is to coordinate the generation of high-quality reference genomes for all eukaryotic life across Europe^18^. At the core of this mission is ensuring the implementation of an inclusive, accessible, and distributed genomic infrastructure that supports the inclusion of all who wish to participate, advances scientific excellence and data sharing best practices, and increases taxonomic, geographic, and habitat representation of sequenced species in a balanced manner. Embracing diversity in this way brings opportunities for ERGA to build a genomic infrastructure that can be used by the large network of biodiversity researchers and also foster new international and transdiscipline collaborations.

The organisational structure of ERGA currently comprises the governing body of the Council of Country/Regional Representatives, with actions developed and implemented by the Executive Board and nine expert Committees, with participation from the large network of members (https://www.erga-biodiversity.eu/). With over 750 members spanning 38 countries, one regional ERGA affiliated project, and 234 institutions, ERGA is currently the largest initiative of its kind in the world. ERGA membership is open to all who wish to engage in the sequencing of European eukaryotes, foster new collaborations in and beyond Europe, and learn about the most up to date technologies for generating reference genomes for species (individuals interested in becoming a member can register through the ERGA website).

A decentralised approach for the production of genomic reference resources for ERGA supports: 1) an expansion in the diversity of expertise, processes and innovative ideas that can act synergistically to accelerate scientific outcomes, 2) a platform for accessible, equitable, and standard production of, high quality, ethically and legally compliant reference genomic resources, 3) streamlined communication and opportunities for new collaborations to be fostered, 4) an expansion of funding opportunities, 5) mitigation of hierarchical power imbalances, 6) increased access to up-to-date and reproducible tools and workflows, and 7) increased downstream analyses applications.

The ambition of the pilot test was to identify the challenges in constructing and implementing a decentralised infrastructure, but also to understand and find solutions on how best to support the inclusion of ERGA members who face a multitude of different realities whilst participating e.g., resource availability, geographic, and political positioning. The lessons learned from this initial pilot can certainly be used by ERGA to inform future developments, but can also be used to inform the broader EBP strategy as to whether decentralised approaches are effective in the production of reference genomic resources that meet with EBP minimum standards.

The first step towards decentralisation was to create a pan-European network of existing sequencing centres, biobanks, and museum collections that were willing to participate and provide diverse support options for sample storage, wet-lab preparation, sequencing, and data handling and storage. The second step was to obtain adequate funding to support the development and implementation of the infrastructure. Here, no central source of funding was available and so the majority of funds were acquired through the grassroots efforts of individual ERGA members contributing to the pilot test as well as a plethora of partnering institutions **(Supplementary Table 1)**. In many cases, researchers completely, or partially, financed their participation in the pilot test. In other cases, sequencing partners contributed their own grant funds to completely cover or offer heavy discounts for the cost of library preparation, sequence data production and/or assembly services whilst also covering the costs of the scientific personnel within their facilities to participate in the pilot test. In addition, collaborations were fostered with commercial sequencing companies to obtain in-kind contributions that could be used to support those researchers who wished to participate but deserved financial support. All in-kind contributions were shipped to three established ERGA Hubs, two ERGA Library Preparation Hubs (University of Antwerp, Belgium and the Metazoa Phylogenomics Lab at the Institute of Evolutionary Biology (CSIC-UPF) in Barcelona, Spain) and one ERGA Sequencing Hub (University of Florence, Italy).

### Development of a Decentralised Infrastructure

Overall, from the 33 countries (17 Widening countries **(Glossary)**) and regions, 98 species were included in the pilot test **(Figure 2a)**. However despite efforts made during the prioritisation process, the dispersion of species selected was not equal across countries predominantly due to the acceptance of additional species after nomination closure **(Figure 2b)**. Nine iterative steps were developed to support the production of a complete reference genomics resource for each of the species included into the pilot project **(Figure 1).**

**Figure 1:**
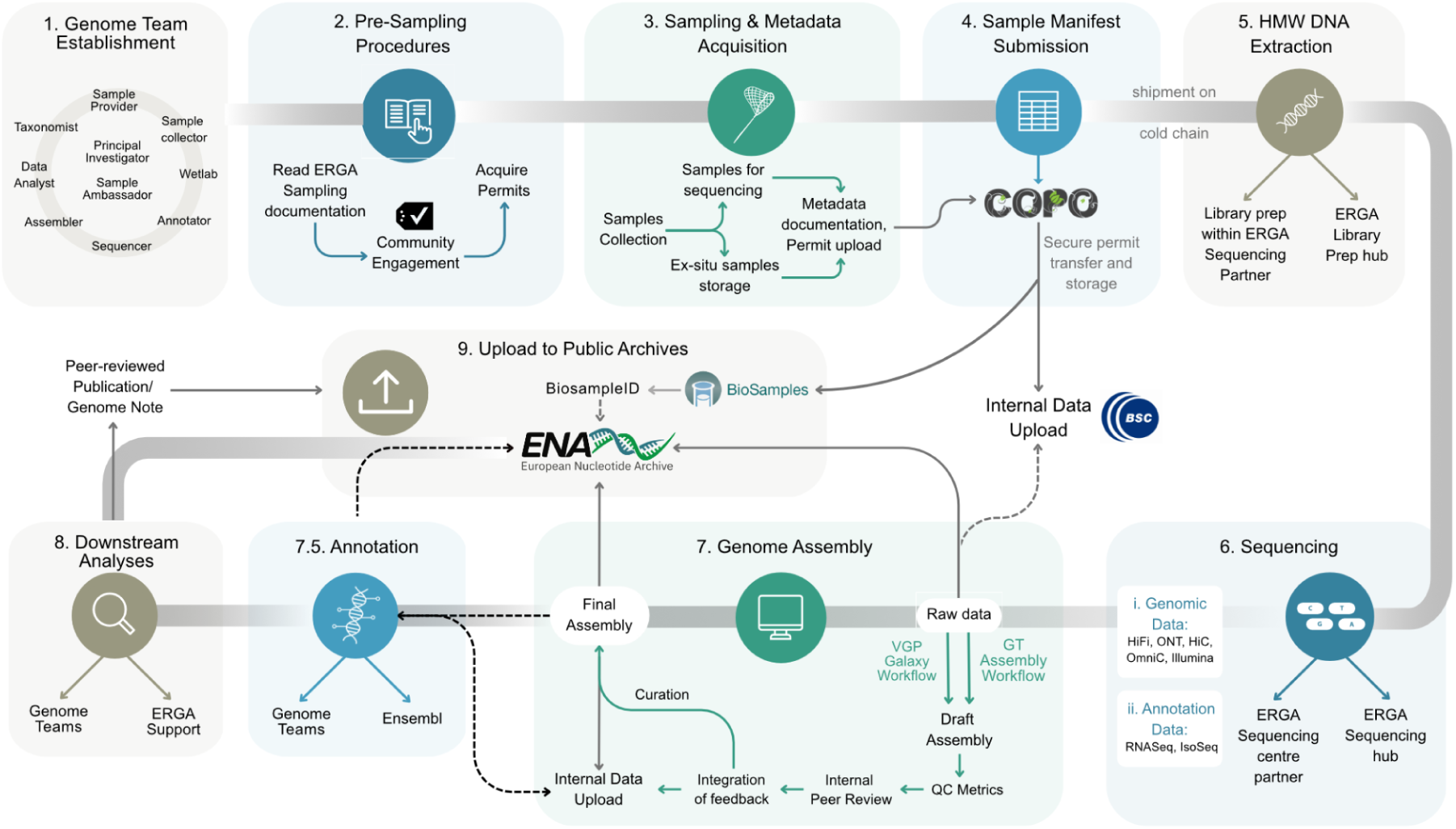
Establishing an inclusive, accessible, distributed and pan-European genomic infrastructure that could support the streamlined and scalable production of genomic resources for all European species.

**Figure 2:**
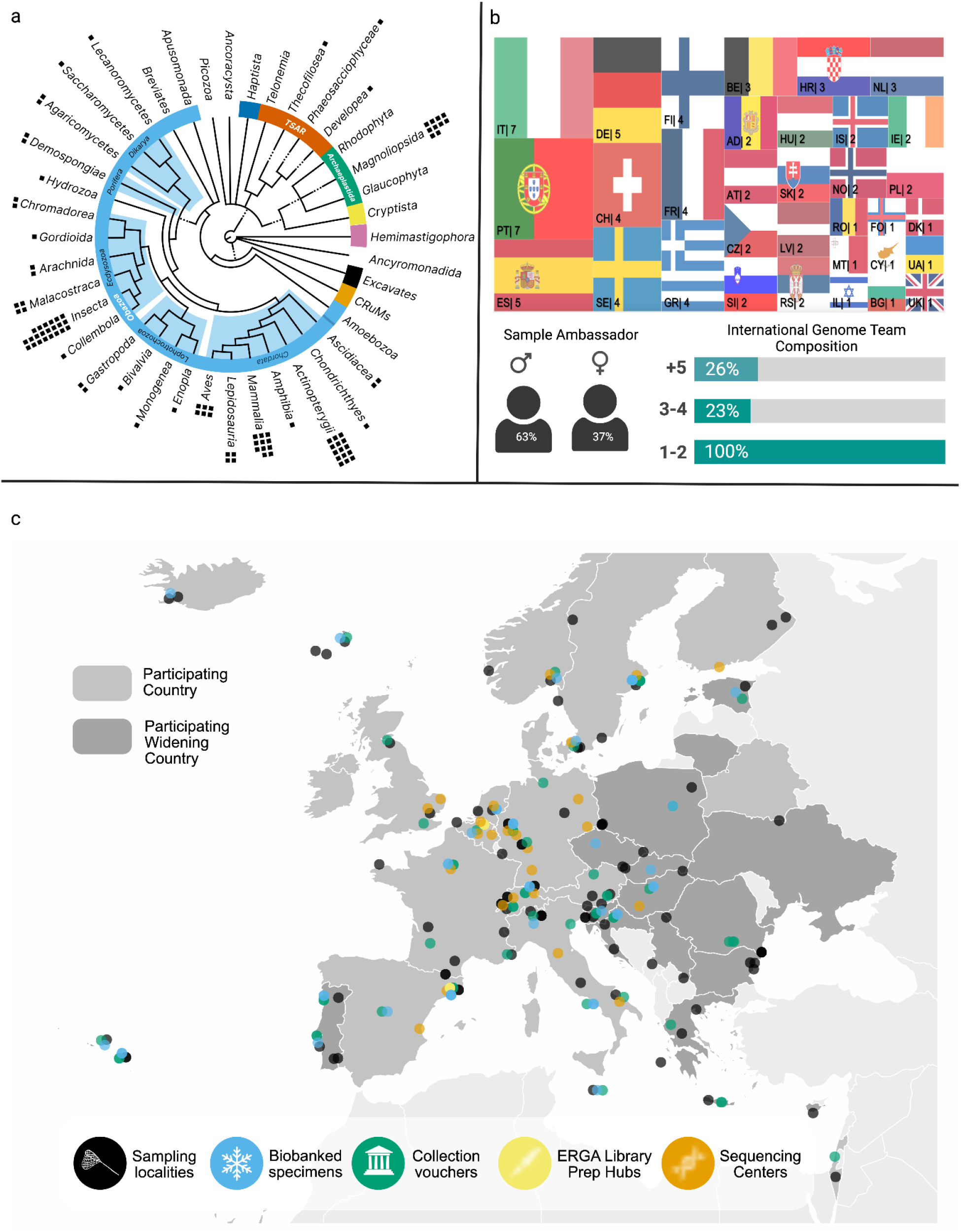
Sample, country and partnering institution distribution across Europe**. a)** Taxonomic distribution of the species included into infrastructure testing. b) Top: Distribution of sample ambassadors per participating country. Bottom-left: self identified sex distribution across sample ambassadors, Bottom-right: frequency of genome teams that have international collaborators i.e., collaborators that are outside of the country of origin that the sample was obtained from. c) Map illustrating the distribution of sampling localities, cryopreserved specimens, collections holding vouchered specimens, sequencing library preparation hubs and sequencing facilities across Europe^32^.

### 1. Genome team establishment

After a successful nomination, including a species into the ERGA infrastructure was reliant on the creation of a ‘genome team’. A genome team is a transdisciplinary group of researchers that have a shared interest in a particular species and assume the shared responsibility of shepherding this species through each of the infrastructure’s steps. Each team member has an assigned role **(Figure 1, Supplementary Table 2)**. Further, all teams were strongly encouraged to include both national and international members and all teams were overseen by the “Principle Investigator” and a “Sample Ambassador” who was ideally from the country of origin of the focal species. The role of the sample ambassador was to coordinate the species project, and to ensure the continuous communication across the team members. In total, 98 genome teams were established and each had at least one international team member, 23% having three members, and 26% having >five members (*n=93*) **(Figure 2b)**. A total of 76 genome team sample ambassadors were comfortable sharing their self-declared sex, (only “male” and “female” were proposed as choices) from this subset, 63 (16%) self-identified as male and 36 (84%) as female **(Figure 2b).** To ensure compliance with GDPR regulations, no other data was collected to assess representation by other critically important dimensions of diversity e.g., race, ethnicity, religion, sexual orientation or their intersections. Hence, ERGA does not currently have any means to evaluate its inclusiveness beyond sex and it is likely that it suffers the same lack of racial representation and inclusion that characterises European science at large^21^.

### 2. Building a representative species list

Prior to developing and testing the decentralised infrastructure **(Figure 1)**, we first needed to consider the species that would test it. For this, a nomination form was issued for completion by all ERGA members that were willing to contribute samples for a species. The form collected information on genome properties, vouchering, habitat and sampling, conservation status, permit prerequisites, sample properties, species identification, and sex (https://treeofsex.sanger.ac.uk/)^19^ for each suggested species^20^. To prioritise nominations, a scoring system was applied based on several feasibility criteria: small genome size (<1Gb), an ease of availability, possibility for being freshly collected and flash frozen, >1g of tissue, a well-established nucleic acid extraction protocol, a specimen voucher present, no species identification ambiguity, all necessary permits existing, and no restrictions on export^20^. ERGA council representatives were given the prioritised species list and asked to select three species per predefined ERGA target category (pollinators, freshwater species and endangered/iconic) from the nominations from members within their country. After nomination form closure, many additional species were nominated by ERGA members. However, only nominations that fulfilled all of the selection criteria, had funding available, and/or were from a country not yet represented were accepted for inclusion into the test.

### 3. Developing a communication and coordination strategy

The nature of the infrastructure constructed required streamlined communication between ERGA genome teams and partnering sequencing facilities spanning large geographic distances. To facilitate this, we created avenues to maximise continuous communication both in and outside of the ERGA community. In partnership with Ensembl at EMBL’s European Bioinformatics Institute (EMBL-EBI)^40^, we built an ERGA Data Portal (https://portal.erga-biodiversity.eu/) to provide a comprehensive overview of all ERGA data. The portal provides a powerful and intuitive ability to search over each ERGA metadata, genomic dataset, assembly and annotation, with filters for component project, sequencing status and taxonomy. Additionally, an interactive phylogeny provides another route to exploring available species, and can display ERGA species sequenced at any taxonomic level. We developed the current portal rapidly to support the goals of the pilot test, but it will be continually and iteratively improved to enhance usability, for example by potentially adding species imagery and distribution ranges, Ensembl^38^ and community annotations, interactive geographic map searches, and cross referencing to key resources such as the Global Biodiversity Information Facility (https://www.gbif.org/) and climate data. Progress data is continuously shared through the portal’s public tracking pages (https://portal.erga-biodiversity.eu/status_tracking) and the GoAT database^16^ https://goat.genomehubs.org/projects/ERGA-PIL).

### 4. Developing a training and knowledge transfer strategy

Investing in building competency is important if ERGA is to provide scientists across disciplines, experience levels, demographic sectors of society, and geographies with equitable opportunities to leverage and benefit from the use of the enormous volume of data expected to be generated through ERGA, but also other large biodiversity genomics initiatives including and especially those in parts of the world where economic opportunities are much more limited. However, a significant gap remains in expertise between countries due to the diverse nature of resource availability, genomic research capacity and capability, and access to state-of-the-art training **(Box 2)**. To increase the accessibility and stimulate the use of existing infrastructure within ERGA across all the infrastructure steps, efforts were made to share expertise through conference participation, webinar organisation and through organising hands-on training workshop opportunities. For instance, many ERGA members participated in a [BioHackathon](https://biohackathon-europe.org/) to integrate new genome assembly methods into an openly accessible Galaxy pipeline and worked on the development of robust user guidance^41^. In addition, we organised a virtual workshop entitled “Building high-quality reference genome assemblies of eukaryotes” as part of the European Conference in Computational Biology 2022^42^ and now freely available online to further educate researchers in best practices for genome assembly. We also organised a webinar on ‘Access and Benefit-Sharing’ with the National Focal Points across Europe to help genome team sample ambassadors to understand their Nagoya permitting obligations during the sample collection stage of the project and organised an online workshop on structural genome annotation with [BRAKER & TSEBRA](https://github.com/KatharinaHoff/BRAKER-TSEBRA-Workshop).

##### Box 2: Opportunities for Training & Knowledge Transfer

During an EMBO Practical Course ‘Hands-on course in genome sequencing, assembly and downstream analyses’ held at the Université libre de Bruxelles (ULB), Belgium (https://meetings.embo.org/event/22-gen-seq-analysis), the organisers chose to use the endophytic yeast *Debaromyces* sp. RF-E1 (13 Mb) for sequencing during the course. Microorganisms are excellent objects for genome sequencing and bioinformatics teaching due to their small genome size (making it possible to try many workflows and sets of parameters). The genus *Debaryomyces* comprises species of extremophilic yeasts, some of which support plant health by modulating pathogen invasion^43,44^. A high-quality reference genome will help study the impacts of radiation on this genome and elucidate the adaptive potential of host-microbe interactions. The yeast was isolated from a silver birch tree in the Red Forest, one of the most radioactive areas in the Chernobyl Exclusion Zone (CEZ) in Ukraine^45^. Anthropogenic stresses caused by radionuclide contamination can adversely affect organism health through genotoxicity^46,47^. Although symbiotic interactions with endophytic microorganisms can facilitate a host’s capacity to adapt and persist under such environmental stress^48^, little is known about radiation exposure’s impact on these endophytic interactions. ONT genomic and cDNA sequencing was performed during the course, then the data were assembled with Flye^49^ and annotated with BRAKER^50^ by the course participants. The pedagogy of the EMBO course effectively combined hands-on research training with the necessary theoretical framing to support active learning of participants. Feedback by course participants was extremely positive, and as a result a second EMBO-funded Practical Course will be organised by the same team in 2024 (this time in Valencia, Spain). In addition to providing participants with a realistic insight into the research process, the training also created a suite of high-quality publicly available genomic resources for the yeast species sequence that will be directly useful to the sample provider’s ongoing research, but also to potentially many more researchers. This successful teaching-through-research model will inform future ERGA training and capacity-building activities at locations across Europe and beyond.

An online workshop was also organised to train pilot genome teams to identify the external actors (international, national, and local levels) involved in their reference genome project. During this training, we conducted a stakeholder and rightsholder, herein interested parties, mapping exercise, and examined sample ambassador perceptions of how to interact with interested parties across high and low GBARD (government budget allocations for R&D) countries. The results indicated that researchers did not categorise their project’s interested parties differently (*X^2^*(3, (153-130)) = 5.66, p = 0.12) **(Extended Data Figure 6)** depending on whether they were situated in a low or high GBARD country. However, there does appear to be a tendency in the ‘Consult’ category (df = 1, p = 0.08), suggesting that researchers located in low GBARD countries may place a higher value on the involvement and collaboration of interested parties as opposed to those located in high GBARD countries **(Extended Data Figure 6)**.

### 5. Technical workflows

#### Pre-sampling requirements

Supporting genome team compliance with all relevant ethical and legal customary, local, regional, national, and international obligations was a priority during the infrastructure development process. Through ERGA expert committees, namely the Ethics, Legal and Social Issues (ELSI) Committee and the Sampling and Sample Processing (SSP) Committee, comprehensive documentation was developed including a “Sampling Code of Best Practice” and “Guidelines on implementing the Traditional Knowledge and Biocultural Labels and Notices when partnering with Indigenous Peoples and Local Communities (IPLC)”^20,22^. The Traditional Knowledge (TK) and Biocultural BC) Label and Notice implementation and guideline documentation was developed through a funded partnership ([European Open Science Cloud Grant](https://eoscfuture-grants.eu/node/257)) with representatives of the Global Indigenous Data Alliance (https://www.gida-global.org/), Local Context Hub (https://localcontexts.org/) and the Research Data Alliance (https://www.rd-alliance.org/node/77186). Complying with this documentation was mandatory as it codifies the official ERGA standards for how to ethically and legally collect samples, as well as how to responsibly engage all interested parties **(Glossary)**. In addition, educational webinars were used as a researcher capacity-building tool, providing more general information on pertinent topics such as the Nagoya Protocol on Access and Benefit Sharing, and Digital Sequence Information (https://www.youtube.com/@erga-consortium1001).

#### Sampling and metadata acquisition

During sample collection important metadata concerning the species collection event were expected to be documented by the sample collector. To standardise this process a robust metadata schema was developed, using the DToL metadata schema as a foundation^23^. The tailored ERGA schema, including unique ERGA specimen identifiers as well as ToLID (https://id.tol.sanger.ac.uk/), was codified into a .csv formatted ‘manifest’ and made publicly available (https://github.com/ERGA-consortium/ERGA-sample-manifest). In tandem, a standard operating procedure document^24^ was developed to provide details on how to complete all of the 81 validatable manifest fields. Inspired by the Genomic Observatories Metadatabase ^25^, ERGA also developed fields to mandate important information disclosure e.g., permanent unique identifiers (PUID) associated with *ex-situ* specimens, permits, and Indigenous rights and interests (TK and BC Labels and Notices)^22,26–28^. Overall, samples were collected for 98 species spanning 92 genera, 81 families, 61 orders, 26 classes, and 13 phyla (https://goat.genomehubs.org/projects/ERGA-PIL, **Figure 2a**). The geographic distribution of samples collected was relatively even, although some countries contributed more species than others **(Figure 2b**). Altogether 89% of genome teams (*n=93*) reported >90% confidence level in that they had obtained all permits required with ten Nagoya permits and three CITES (**Supplementary Case-study 1**) permits being obtained.

#### Sample manifest submission, validation, ex-situ storage

An accessible and streamlined metadata manifest submission system was implemented to ensure that all ERGA’s sample metadata was accurately validated and promptly submitted into the public archive. To achieve this, a user-friendly and highly customised data and metadata brokering system called Collaborative Open Omics (COPO) (https://wellcomeopenresearch.org/articles/7-279/v1) was used^29^. The COPO submission system validated each manifest submitted against an ERGA provided checklist to standardise and automate entry into the BioSamples public archive. By automating this process it ensured that all species samples collected had a permanent unique identifier (PUID) from BioSamples that can be automatically linked to the associated genomic sequencing data submitted to the European Nucleotide Archive (ENA; https://www.ebi.ac.uk/ena). Additionally, the submission system had the capability to upload permit documentation and supported its immediate transfer to a private and secure location on an internal ERGA data repository (that was built for the purposes of the pilot test) to avoid privacy concerns and data leakages. All documents were subsequently deleted from COPO’s internal servers. The internal data repository itself was constructed in partnership with the Barcelona Supercomputing Centre (BSC; https://www.bsc.es/), and was a Nextcloud instance containing a group folder with a tiered storage system, or HSM **(Glossary)**. All ERGA members could request access to the ERGA data repository and upon approval, members were assigned appropriate access privileges depending on their needs (read-, write-, or full file control access). To support repository utilisation, guidelines were developed detailing protocols for data upload/download as well as directory structure, to ensure standardisation, reusability, and interoperability^30^.

We highly recommended that both voucher specimen(s) and cryopreserved specimen(s) be associated with all genomic resources produced during the pilot test. To support this we issued supporting guidance for biobanking and vouchering^31^. The vouchering best practices developed recommended the deposition of both a physical and digital e-voucher(s) (high-quality, informative photographs). Through ERGA’s SSP Committee, we also supported genome teams in seek of a permanent collection for voucher deposition and a partnership with the LIB Biobank at Museum Koenig (Bonn) (https://bonn.leibniz-lib.de/en/biobank) was established to support the deposition of cryopreserved samples for those without access to a local biobank. Samples biobanked in LIB were made publicly visible via the international biodiversity biobanking portal (GGBN.org). Although not a mandatory requirement, voucher specimens were provided for 67% of the species (19% digital, 40% physical, and 40% had both physical and digital) and deposited in museum collections across 23 countries (**Figure 2c**). Of the specimens, 45% had an associated cryopreserved sample that were stored in 34 biobanks in 22 countries **(Figure 2c)**. All 98 genome teams successfully completed, validated and uploaded their metadata publicly to BioSamples through the COPO system and manifest submissions are publicly available through the ERGA Data Portal (https://portal.erga-biodiversity.eu/) that provides intuitive search and direct links to all of the data held in the public archives (**Communication and Coordination Section)**.

#### Sample Preparation

Sample quality and shipment requirements were formalised for each data-type across ERGA sequencing facility partners, including sample requirements for long reads (Oxford Nanopore Technologies (ONT)/Pacific BioSciences (PacBio)), scaffolding (Omni-C/Hi-C), and annotation (RNA-Seq/IsoSeq) of data^31^. Sample collectors were expected to adhere to the requirements of the ERGA sequencing facility specified and ensure that samples shipped are: 1) of a quality suitable for HMW DNA extraction, and 2) of an appropriate quantity for long-read, proximity ligation and annotation sequence data production. Two ERGA Library Preparation Hubs were established to support genome teams that required resource support for the library preparation of samples prior to sequencing. To increase the likelihood that the HMW DNA of sufficient quantity was obtained for effective sequencing, most library preparation was conducted by partnering sequencing facilities. However, the ERGA Library Preparation Hubs facilitated the production of 99 libraries: 15 libraries for proximity ligation data (Hi-C/Omni-C® kit) that were provided by 27 countries; Eight libraries for PacBio data provided by eight countries; and the remainder were for RNAseq data **(Supplementary Table 3, 4, Extended Data Figure 1,2).**

#### Sequencing Strategy

A key component and strength of the decentralised infrastructure was the intentional distribution of sequence data production across partnering European sequencing facilities. To initialise these partnerships, a sequencing platform landscape assessment was conducted across all of the countries that had ERGA council representation. This effort assessed the quantity, distribution, and diversity of the sequencing platforms available across Europe and specifically examined their capability to produce long read (PacBio HiFi reads/ONT reads /IsoSeq reads), and short read (Hi-C/Omni-C/RNA-Seq/PCR-free Illumina) sequencing data. This mapping indicated an uneven distribution of sequencing platforms across Europe, and so we decided that any sequencing facility with a platform to produce long read sequencing data could be an ERGA partner. We took this long read data-type agnostic approach to maximise geographic breadth and increase accessibility but also to reduce shipping costs and the likelihood of customs issues. An additional strength was that it could facilitate the development of more standardised and automated approaches for long read technologies that are currently underrepresented in generating genomic references for biodiversity genomics. Supporting a variety of technologies is important as it takes advantage of their individual characteristics (e.g., portability or lower priced solutions) to increase sequencing capability and accessibility in under-resourced countries, regions and institutions in ERGA. In the end, we partnered with a total of 26 sequencing facilities, 17 with PacBio and 9 with ONT sequencing platforms available **(Figure 2c)**, and documented the minimum sample collection and quality requirements for each partner^31^. Here, we recommended the following data-type volumes for assembly generation: 30X HiFi or 60X ONT, 25X Hi-C (per haplotype) and 25X (per haplotype) Illumina (in cases where ONT data was used), and the following data-type volumes for annotation: total of 100 million reads if >five tissue types are available, or 30 million reads if tissue samples are pooled^32^. IsoSeq production was not a mandatory requirement but was promoted, where feasible. The pilot test’s 98 species were sequenced across 25 main partnering sequencing facilities (**Supplementary Table 1**), and additional data was generated by Novogene for four species from the Netherlands and Hungary. 27 species were sequenced using a ONT platform, 75 using the PacBio Sequel II platform, and four by both platforms. For scaffolding and curation purposes, proximity ligation sequencing was highly recommended. A total of 76 species had some form of proximity ligation sequencing conducted, 47 species with Arima-Hi-C (Arima Genomics), 24 species with Dovetail Omni-C® (Dovetail Genomics), and five with Proximo (Phase Genomics) **(Figure 3a).** Regardless of the partnering sequencing facility utilised or species being sequenced, the facilities were expected to produce sufficient data to reach at minimum EBP recommendations^33^. An ERGA Sequencing Hub was also established at the University of Florence (Italy) Genomics Core to support the sequencing of the 99 libraries prepared by the ERGA Library Preparation Hubs **(Supplementary Table 3,4**). Upon sequencing data generation, both genomic and transcriptomic data were shared with the genome teams through the internal ERGA data repository.

**Figure 3:**
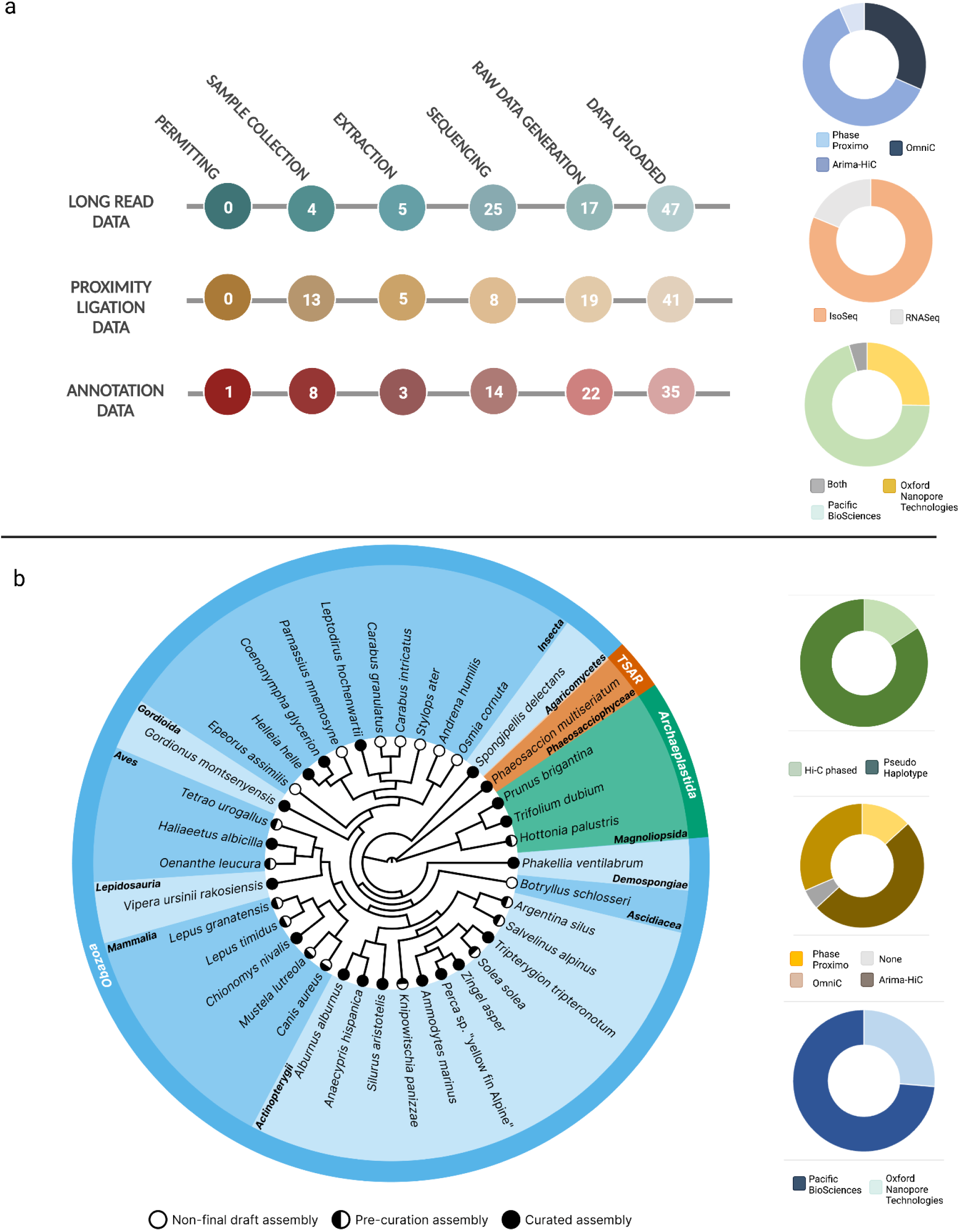
Pilot test data production per species progression. a) total data production progress across all 98 species included, noting that data not planned/required for 12 species for proximity ligation, and 15 species for annotation data. b) species distribution of species with genome assemblies available, both draft and curated assemblies are shown here. The data-type distribution for these species is also supplied. See **Extended Data Figure 3** for complete species tree.

#### Genome Assembly and Annotation

A requirement for becoming an ERGA reference genome was that the genome assembly reached, at minimum, the EBP standard for assembly quality^33^. To ensure the infrastructure supported the production of genomic references to this standard, we developed assembly guidelines with workflows tailored for both ONT and HiFi based genome assemblies^34^. The use of these workflows was not mandatory, and any assembly workflow would be accepted if the resulting assembly met the appropriate assembly quality^33^. To streamline the assessment and validate all ERGA genomic references, we established a stepwise procedure of 1) QC metrics assessment, 2) internal peer-review, and 3) manual curation. On completion of a draft assembly, each genome team reported a set of standard QC metrics^35^ that include a contaminant assessment, K-mer metrics, Hi-C map and graph production, gene prediction analyses, and a set of summary statistics. After this, the assembly and the associated metrics underwent an internal round of peer-review from assembly experts (ERGA Sequencing and Assembly Committee). After feedback integration, each genome team uploaded the pre-curation assembly to the internal ERGA data repository along with details of the assembly construction (https://gitlab.com/wtsi-grit/documentation/-/blob/main/yaml_format.md) and each team was provided with the opportunity to submit their reference genome to an internal panel of expert curators who conducted a final manual curation^36^.

Due to the decentralised nature of the infrastructure, all 98 species progressed through the steps at different rates, depending on the number and complexity of permits **(Supplementary Case-study 1)**, difficulty of sample collection **(Supplementary Case-study 2)**, need for sample specific protocol development **(Supplementary Case-study 3)**, partnering sequencing facility capacity, and assembly complexity. **Figure 3a** highlights the current status of each species that has an assembly generated and shows that 13 complete and curated reference genomes have been generated (11 of which can be found in the INSDC), a further 17 are complete but require curation, and 8 are in non-final draft stage.

From the 30 reference genome assemblies with a ‘Curated’ or ‘Pre-curation’ status, we found 14 cases where the assemblies do not meet the quality standard 6.C.Q40 EBP standard criteria **(See glossary and Supplementary Table 6, Figure 4a)**. For instance, *Argentina silus* (fArgSil1) and *Knipowitschia panizzae* (fKniPan1) have scaffold N50 values that meet the minimum requirement, indicating successful Hi-C scaffolding, however both fall short in terms of contig contiguity (N50 < 1 Mbp). In addition, those two pre-curation assemblies contained many small scaffolds, which increased the total number and translated to higher values of Scaffold L95. Notably, *Phaeosaccion multiseriatum* (uoPhaMult1) meets the contig N50 but does not meet the scaffold N50 metric (N50 > 10 Mbp). In the cases of *Spongipellis delectans* (gfSpoDele1) and *Phakellia ventilabrum* (odPhaVent1), they reached a chromosomal scale N50 scaffolding (6.C.Q40), but not the N50 threshold used as a proxy in **Figure 4a** (6.7.Q40), a minimum criteria set for vertebrates but that cannot be applied to taxa with chromosome length N50 less than 10 Mbp.

**Figure 4:**
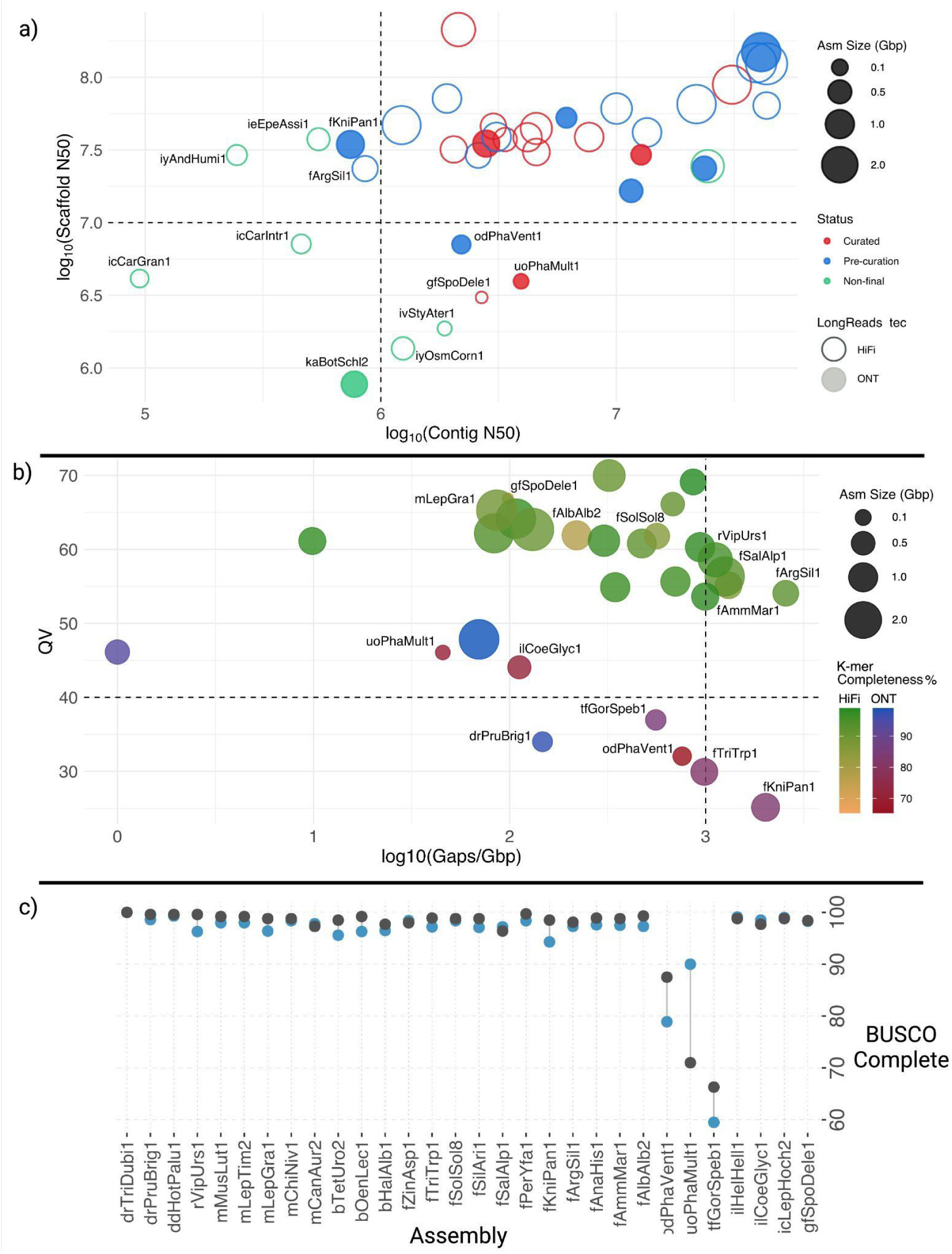
Quality control and status of the 38 genome assemblies evaluated. a) Genome assemblies are represented according to their Scaffold N50 (y-axis, log_10_) and number of the longest scaffolds that comprise at least 95% of the assembly (x-axis, log_2_). Bubble size is proportional to assembly span. Empty bubbles depict HiFi-based genomes, while full bubbles are ONT-based. Colours are according to assembly status (Curated, Pre-curation, Non-final draft). Lower values for both axes indicate better assembly contiguity. Assemblies not reaching the EBP-recommended One Megabase Contig N50 (log_10_1,000,000=6) or 10 Megabase Scaffold N50 (log_10_10,000,000=7) here a proxy for chromosome-level scaffolds, are labelled with their ToLIDs* (https://id.tol.sanger.ac.uk/). b) Completed HiFi- and ONT-based genomes assemblies are represented according to their Quality value (QV, y-axis) and number of gaps per Gbp (log_10_, x-axis). The bubble size is proportional to assembly size. Colour grade of the bubbles is according to the K-mer completeness score. ToLIDs are reported for the assemblies that are below the recommended EBP metric for QV (40), Gaps/Gbp (log_10_1,000=3) or K-mer completeness (90%). Quality values are calculated differently for HiFi-based assemblies than for ONT-based assemblies and should not be compared directly. c) BUSCO completeness scores for genome assemblies with ‘Curated’ and ‘Pre-curation’ status. Using two orthologs databases, one for a more recent last common ancestor encompassing related species (blue), and one for all eukaryotes (grey), we seek a more comprehensive estimation of the assembly completeness. Number of single-copy orthologs present on each database is reported. *Briefly, a ToLID is a unique identifier for an individual organism within a species sampled for genome sequencing, consisting of one or two lowercase letters for high-level taxonomic rank and clade, respectively, followed by three letters for genus and species each. Thus, within insects (i), the Hemiptera (i) includes Andrena humilis (iyAndHumi1) and Osmia cornuta (iyOsmCorn1). The Coleoptera (c) contains Carabus granulatus (icCarGran1), C. intricatus (icCarIntr1), and Leptodirus hochenwarti (icLepHoch2). Ephemeroptera (e) features Epeorus assimilis (ieEpeAssi1), and among Strepsiptera (v) it is found Stylops ater (ivStyAter1). Lepidoptera (l) includes Coenonympha glycerion (ilCoeGlyc1), Helleia helle (ilHelHell1), and Parnassius mnemosyne (ilParMnem1). Within the fungi (g), Agaricomycetes (f) are represented by Spongipellis delectans (gfSpoDele1). For sponges (o), Demospongiae (d) includes Phakellia ventilabrum (odPhaVent1), and among algae (u), Heterokontophyta (o) are represented by Phaeosaccion multiseriatum (uoPhaMult1). The fishes (f) include Alburnus alburnus (fAlbAlb2), Ammodytes marinus (fAmmMar1), Anaecypris hispanica (fAnaHis1), Argentina silus (fArgSil1), Knipowitschia panizzae (fKniPan1), Perca sp. ‘yellow fin Alpine’ (fPerYfa1), Salvelinus alpinus (fSalAlp1), Silurus aristotelis (fSilAri1), Solea solea (fSolSol8), Tripterygion tripteronotum (fTriTrp1), and Zingel asper (fZinAsp1). Birds (b) are represented by Haliaeetus albicilla (bHalAlb1), Oenanthe leucura (bOenLec1), and Tetrao urogallus (bTetUro2). Mammals (m) include Canis aureus (mCanAur2), Chionomys nivalis (mChiNiv1), Lepus granatensis (mLepGra1), Lepus timidus (mLepTim2), and Mustela lutreola (mMusLut1). Among reptiles (r) is Vipera ursinii (rVipUrs1). Within dicotyledons (d), the Ericales (d) include Hottonia palustris (ddHotPalu1), and Rosales and Fabales (r) features Prunus brigantina (drPruBrig1) and Trifolium dubium (drTriDubi1), respectively. Finally, among ‘other chordates’ (k), Ascidiacea (a) includes Botryllus schlosseri (kaBotSchl2), while in the category ‘other animal phyla’ (t), Nematomorpha (f) is exemplified by Gordionus montsenyensis (tfGorSpeb1).

We found differences between HiFi- and ONT-based assemblies in the K-mer-based analyses, for example the average quality value (QV) for HiFi-based assemblies was 61, while for ONT-based it was 38. From these ONT assemblies, five species showed values below the recommended 40, which corresponds to an error rate > 0.01% (**Figure 4b**). It should be noted that in the case of ONT-based assemblies K-mers were derived from orthogonal Illumina reads from the same individual, whereas in the case of Hifi assemblies the K-mers were derived from the same data used to generate the genome assembly, likely inflating QV estimation due to data-interdependence. Further research is warranted on how to mitigate this issue. Recent unpublished results from within ERGA suggest that assembly of newer ONT data (Kit14, Q20+) consistently generates assemblies with QV>40, perhaps side-stepping this issue. Eleven species showed K-mer completeness below 90%, with four being below 80% and one also lower than 70%. Out of these, six belonged to ONT-based assemblies while eight had curated status (**Figure 4b**). A caveat to K-mer completeness is that pseudohaploid assemblies (the typical output of ONT-based assemblies) of heterozygous genomes tend to have lower K-mer completeness. This highlights the need for continued development of diploid assembly strategies to ensure high K-mer completeness.

Five genomes exceeded the recommended metric Gaps/Gbp^23^ as they all had >1,000 remaining (*Argentina silus* (fArgSil1), *Knipowitschia panizzae* (fKniPan1), *Ammodytes marinus* (fAmmMar1), *Salvelinus alpinus* (fSalAlp1) and *Vipera ursinii rakosiensis* (rVipUrs1)). Despite this, for all the completed assemblies, Ns accounted for less than 0.05% of the genome, with the exception of *Mustela lutreola* (mMusLut1). For this genome assembly, which has yet to undergo final curation (the only large ONT-based assembly evaluated >2 Gbp), 0.55% of its sequence was composed of Ns **(Figure 4b)**.

Besides EBP metrics, when estimating completeness using single-copy orthologs, *Phakellia ventilabrum* (odPhaVent1) and *Gordionus montsenyensis* (tfGorSpeb1) assemblies had lower values than recommended. tfGorSpeb1 is one of the first of its phylum to be sequenced^37^, and so is therefore underrepresented in the BUSCO database (**Figure 4c**)^38^. Two species, *Trifolium dubium* (drTriDubi1) and the *Salvelinus alpinus* (fSalAlp1), both have higher ploidy levels (tetraploid and partial tetraploid, respectively) and had much higher BUSCO duplicate values than the recommended 5% (**Supplementary Table 1**).

For the pilot test, the sample collection process for the included species was ideally conducted to facilitate simultaneous genomic and transcriptomic data production. After data deposition to the ERGA data repository, we designed the infrastructure to have the flexibility necessary for each genome team to decide whether the annotation will be conducted i) by the genome team or sequencing facility, ii) with supporting expertise from the internal ERGA community, iii) or wait until the assembly and annotation data is uploaded to ENA where a gold standard annotation will be generated by Ensembl^39^. Although annotation was not mandatory, we produced sequencing data to support annotation data for 81 species (66 with RNA-Seq data, and 15 IsoSeq data). For those species with IsoSeq data generated, 13 also obtained RNA-Seq data. For the 30 genome teams spanning 16 countries that lacked the resources necessary to generate annotation data, we ensured that samples were shipped to a dedicated ERGA Library Preparation Hub. Here, 76 libraries were prepared and shipped to the ERGA Sequencing Hub for data production. In some groups annotation is still underway, but seven genome teams reported that they have a finalised annotation.

#### Data Analysis

Reference genomes can support many downstream analyses, including population genomics, phylogenomics, functional genomics and comparative genomics^9^. Following the assembly and annotation of the newly-built reference genomes, we offered assistance through the ERGA Data Analysis Committee to genome teams by suggesting and supporting avenues of downstream data analyses that could be followed to answer their biological questions of interest. In addition, we connected genome teams with relevant ERGA members that may be able to assist or mentor downstream biological exploration, sparking new collaboration and working groups. As many of the 98 species participating had not yet reached the point of data analysis, we conducted a brief survey to better understand what downstream analysis was planned across the genome teams participating **(Extended Data Figure 5)**. For 59.8% of genome teams, the downstream analyses planned would not have been possible without the reference genome, and 70.7% reported that their planned analyses will be significantly improved by the availability of the reference genome, reinforcing that the biodiversity genomics community is in great need of genomic resources of this kind and quality. Results across the genome teams indicate that the most common type of downstream genomic analyses planned was population genomic based analyses (37.7%) for assessments of population history, structure and status of endangered and endemic species (e.g. demography, inbreeding, hybridization, and association with morphological or environmental factors). Comparative genomics was also a common analysis type across genome teams (27%) who seek to examine relevant evolutionary processes across species (e.g. trait-associated gene family evolution analysis, repeat content evolution, synteny, inversions, tRNA evolution).

Overall, the results of this survey show that the availability of reference genomes are considered a key tool for downstream applications.

#### Upload to Public Archives

To follow the principles of Open Access to Scientific Publications and Research Data [Guidelines](https://ec.europa.eu/research/participants/docs/h2020-funding-guide/cross-cutting-issues/open-access-data-management/open-access_en.htm) of the European Research Council under Horizon 2020, ERGA adopted the data policy of “as open as possible but as closed as necessary”. To support this policy, we developed an ERGA Pilot Project Data Sharing and Management Policy^40^ specifically seeking to balance data openness with respecting the needs of diverse ERGA genome teams. The policy itself codified that all reference genome, annotation and raw sequence data was expected to be uploaded upon generation to the internal ERGA data repository, ensuring its immediate accessibility to the ERGA community. The policy also grants each genome team the ability to place an embargo on public upload of ERGA data into the public archives until the first publication but no longer than two years after data release. Laid clear in the policy is the provisions for fair and rightful attribution in all associated publications.

### Decentralisation Challenges

From the outset of the pilot test, we realised that the decentralised infrastructure built would have huge implications on who was included, had access to, and benefited from the production of genomic resources into the future. Collecting, identifying, storing, and cold-chain shipping of specimens as well as producing, analysing, and storing sequencing data is expensive, requiring ex-situ long term storage facilities, sequencing equipment, laboratory access, a skilled workforce, and significant computational resources. The resources to create genomic resources are neither evenly distributed across the globe, nor across Europe. A key goal of the pilot test was to identify how the existing inequitable structures and systems would manifest whilst building a distributed genomic infrastructure. Intertwining and embedding justice, equity, diversity and inclusion into the scientific mission was considered essential if a decentralised, accessible, and scalable infrastructure was to be achieved that truly supported the production of complete reference genomics resources for all species, and was accessible to all researchers. Overall, the main objectives we set out for the decentralised infrastructure were achieved as it: i) supported the ethical and legal production of high quality genomic resources; ii) created a network of the researchers and institutions engaged in the field of biodiversity genomics; iii) leveraged the network’s existing institutional capacities and capabilities; and iv) harnessed the diverse expertise of the ERGA memberbase and streamlined, as much as possible, equitable participation. However, the decentralised approach also revealed a number of challenges that need to be addressed by ERGA moving forward.

### 1) Technical

#### Phylogenetic representativeness and sampling bias

Bias was found in the representation of countries (**Figure 2**), distribution of species sampled per country (even when population size is considered **(Extended Data Figure 7)** and species distribution across the phylogenetic tree. Generally, non-Widening countries were more strongly represented than Widening countries and certain branches of the tree of life were overrepresented (Mammalia, Aves, Actinopterygii and Magnoliopsida), whilst others (Insecta, Amphibia Mollusca, Annelida, Fungi and most protist groups) were underrepresented. Feasibility was another obstacle. First, the production of long-read and -range sequencing on a species sample requires a significant amount of HMW DNA per 1 Gb of genome size and so small-sized species or species with very large genomes remain an unsolved challenge **(Supplementary Case-study 2).** Second, for some taxa and species, co-purification of secondary compounds resulted in sequencing chemistry interferences. Finally, ideal tissue preservation was not always possible due to sampling at remote destinations or from scientific collections where samples were preserved a long time ago **(Supplementary Case-study 6)**.

Moving forward, a more robust species prioritisation process could ensure that all species are assessed using clearly specified criteria with a scoring system that is responsive to the needs of both equity deserving countries **(see glossary)** and underrepresented taxa. For example, species from higher taxonomic groups without reference genomes could be prioritised over those more resource abundant groups or Widening countries could be prioritised over non-Widening countries. A more robust species prioritisation process could also facilitate knowledge transfer and serve as a seed for national investments in biodiversity genomics. Tackling these challenges will require a greater investment in research and development as well as highly-skilled personnel, additionally researchers may need incentives to prioritise the interest of species or taxa that remain underrepresented in public databases.

#### Enhancing end-use through genome annotation

The first hurdle in annotation is the availability of sufficient evidence (transcriptomic and protein sequence data) from focal species, databases and predictive models of repeats. Secondly, even with appropriate data, the most accurate genome annotation pipelines require advanced skills to both install and run which reduces their accessibility and ultimately their utility. Finally, robust annotation quality assessment tools are lacking particularly for species with underrepresented genomic resources, for instance gene content assessment tools such as BUSCO^52^ remain unable to account for species within taxonomic groups that have incomplete gene sets available leading to unreliable quality assessments.

Obtaining, and equitably distributing, financial resources will be required to equip researchers, labs, and regions for annotation in a manner that responds to their varying resource realities. Additionally, the development of more easily installable and reproducible pipelines are needed, and thankfully some new tools are now emerging with this in mind^53^. Standardised and streamlined annotation pipelines are needed for consistency which is crucial for many analyses such as comparative genomics as it can facilitate more confident comparisons. Finally, sequencing more underrepresented genomes will help improve quality assessment tools. Filling in phylogenetic gaps will provide more opportunities for comparisons among taxa but also to develop better models for gene predictions. Despite these challenges, it is important that genomes are annotated. Many downstream analyses are based solely on the predicted genes from the annotation, and incomplete or incorrect results will negatively impact studies of both short-term and broad evolutionary processes.

#### Decentralising reference production and reproducibility

During the pilot test the reference genomics resources were produced across diverse and transdisciplinary research groups, institutes and countries. This diversity resulted in variances in accessibility, capacity and capability in sequencing technologies, computation, and software but also across different taxa. The overrepresentation of pilot sequencing facility partners located in Western Europe compared to Eastern Europe demonstrates such disparity. Furthermore, the data agnostic approach taken led to challenges in standardising assembly, annotation and curation protocols, workflows and procedures across the project. For instance, a blanket adoption of the VGP pipeline for diploid genomes based on PacBio HiFi and Hi-C sequencing (https://gxy.io/GTN:T00039; https://workflowhub.eu/workflows/325?version=1) was not appropriate as this approach would not cater for polyploid genomes nor those assemblies produced that were ONT-based. A further challenge was the provision of a centralised system for the storage and transfer of raw and final genomic and transcriptomic data. This was particularly challenging in cases where data production spanned two or more locations (e.g., PacBio sequenced at one site, Hi-C at a second, and RNA at a third) and was subsequently assembled at another site. While the Nextcloud instance created by BSC was an elegant solution for transferring vast quantities of data between parties, it required a vast amount of personnel hours to manage, in addition to its baseline system-wide maintenance requirements.

Moving forward, a key goal for ERGA is the production of standardised and reusable pipelines that are: responsive to all sequencing “recipes” (PacBio, ONT, or other future technologies); written for Galaxy, Snakemake, and/or Nextflow workflow managers; made publicly available (https://github.com/ERGA-consortium/pipelines); and are actively maintained by the ERGA community with regular scheduled and versioned updates. It would also be beneficial to diversify the availability of sequencing instruments to allow for more instances where sequencing and assembly can be produced concurrently at the same location, reducing the need for transferring files that can reach up to 1Tb in size.

### 2) Ethical and legal

ERGA is an international initiative and so safeguarding production of only ethical and legal reference genomes was a complex endeavour. Decentralisation of the infrastructure resulted in many species samples being transported across national and regional jurisdictions as well as in and out of the European Union, creating an ethical and legal compliance tribulation. Additionally, depending on the species in question, the legal landscape may differ drastically e.g., CBD^54^, CITES^55^, ITPGRFA^56^, UNCLOS^57^ etc. Understanding legislation can be complex and difficult especially for researchers who do not have formal legal training, usually lack legal support within their institution, and often do not have the time or resources to acquire either. This created uncertainty amongst many researchers, especially those navigating this for the first time **(Supplementary Case-Study 1)**. To add to this uncertainty, the pilot test coincided with international discussions on the fair and equitable sharing of benefits from the access and use of digital sequence information (i.e., genomic sequences) under the Nagoya Protocol adding increased uncertainty surrounding the legal compliance landscape^58^. Additionally, although researchers were supplied with documentation and infrastructural support to aid ethical and legal compliance, the pilot test had no means to monitor compliance.

Moving forward addressing the ethical, legal and social implications of ERGA will require professionalisation through a dedicated funding stream. Funded positions will attract trained personnel with the necessary experience needed to navigate complex permitting issues and compliance monitoring. Additionally, a greater effort needs to be made on training ERGA members on the importance of ethical and legal compliance in biodiversity genomics research.

### 3) Social justice

#### Building a more socially just infrastructure

Building a truly inclusive, diverse and equitable infrastructure for biodiversity genomics faces structural constraints. They are mainly twofold: first, lack of equity for and inclusion of minorities in science within the countries of Europe^21,59^; second, extreme economic and political inequity between Europe and countries in the Global South^60^. For the pilot test, no data was collected on race, ethnicity, religion, sexual orientation, disability, career-level or the intersections of these with gender and with each other. This data deficiency made it impossible to critically evaluate the consortium in terms of inclusiveness. Some preliminary data generated in regards to sex suggested that by allowing genome teams to organically form it resulted in sex imbalances. Hence, there is a high likelihood that this also resulted in an underrepresentation of many other minoritised groups^21^, a known trend across European science^21^. The second constraint arises from a pressure to confine a biodiversity genomics consortium to the political boundary of Europe and the nation states within. Europe, and the nations within it, are not naturally occuring units of biodiversity. In fact Europe is part of a much wider biogeographical realm (the Palearctic) that includes large parts of Africa and Asia^61^ (https://www.britannica.com/science/biogeographic-region).

As ERGA progresses, the consortium should prioritise the collection of applicable demographic data. Moreover, outreach activities should be conducted to explicitly recruit researchers from sectors of the population that are underrepresented in science. To really address the biodiversity crisis in a meaningful way, it will be important for ERGA to expand its reach globally. Afterall, most biodiversity by far resides not in Europe but in the Global South. Much commitment and ingenuity will be required to overcome the effects on biodiversity genomics of the equity gap that separates Europe as a block from many countries in the Global South. It will be a challenge to overcome the boundaries and constraints often dictated by scientific funding, but it is a challenge that must be overcome on the road towards a sustainable future. Through harnessing the power of its positioning in the EBP, ERGA should make efforts to become more integrated with other ongoing and related initiatives in neighbouring regions, e.g.,Africa BioGenome Project^62^.

### Prioritising engagement and outreach

Effective engagement is commonly seen as a constraint rather than an opportunity due to resource and time limitations, and a lack of training and awareness. Although a virtual workshop was provided during the pilot test to train researchers on 1) the significance of interested party engagement and 2) the skills to identify, map, and comprehend the needs of potential interested parties, it remained a challenge to transition researcher focus from reference genomes to the practical applications of genomics more broadly. Additionally, although the infrastructure was designed to recognise and include the rights and interests of Indigenous Peoples and Local Communities (TK and BC Labels and Notices, supporting guidelines for researcher implementation, and an “Open to Collaborate’’ Notice on the ERGA website), researchers require more training on why and how to proactively engage and establish sustainable partnerships with Indigenous Peoples and Local Communities.

Overall, more training is needed for interested party identification, mapping, tailored engagement (varying interests and cultural perspectives), and communication. To address this, a comprehensive framework that encompasses targeted communication strategies, tailored dissemination channels, and proactive exploitation of research findings would be useful. This plan, if developed, could ensure that all interested parties receive timely and relevant information, fostering broader awareness, understanding, and utilisation of the results generated by biodiversity genomics research. Supporting ERGA members in this way could empower researchers to get more involved at the interface between biodiversity genomics research and biodiversity policy **(Supplementary Case-study 4)**.

### Scaling training and knowledge transfer

Financial resources are not equally distributed among countries, institutions or researchers, leading to limited access to crucial state-of-the-art training, resulting in significant disparities in terms of the expertise required to access and utilise these resources. Given the economic privilege that even the least wealthy EU countries have when compared to countries in the Global South, it is clear that access to funding for mobility is a huge barrier globally. Throughout the pilot test, several trainings were held and guidelines developed to enhance the user-friendliness of the infrastructure as well as to streamline its use; however there was no clear long-term strategy for training and knowledge transfer.

To develop a genomics curriculum that is responsive to the needs of researchers and trainees, and promote the long-term building of capacity within these countries, an investment into a long-term strategy will be required. For instance, a publicly available knowledge transfer platform could be created to provide ERGA members with resources and trainings relating to each step of reference genome production, but could also provide links to complementary initiative resources e.g., EBP, Elixir (https://elixir-europe.org/), Galaxy (https://usegalaxy.org/), DSI Network (https://www.dsiscientificnetwork.org/), gBIKE (https://g-bikegenetics.eu/en), CETAF (https://cetaf.org/) etc. Such a platform could also provide a space for the sharing of relevant biodiversity genomics educational materials that could further aid collaborations between researchers who are shaping the future of biodiversity genomics curricula development globally.

### Future Directions

The decentralised approach taken by ERGA through the pilot test illustrates the huge potential of the consortium to become a model for equitable and inclusive biodiversity genomics in the future. The power of such an approach was evident through the momentum it built across its participants. Not only did the pilot test successfully unite an international community of biodiversity researchers, but it also stimulated communities of researchers within the same country to combine and consolidate efforts under the ERGA umbrella e.g., DeERGA and Portugal BioGenome^63^. Additionally, it allowed participating researchers to apply the lessons learned from the test to build localised infrastructures that would remain interoperable with partners across Europe, e.g., ATLASea.

A key aim for testing the approach was making visible the challenges and issues that would manifest whilst working at an international level, and at scale and working to improve and build upon these learnings as the consortium moves forward. Some key challenges highlighted by the pilot test concerned: species selection processes (criteria, prioritisation) and sampling procedures (permitting, collection, preservation, metadata); modes of engagement across interested parties (citizen scientists, policymakers, Indigenous Peoples, Local Communities etc); the diversity and inclusion of the researchers participating; defining the scope of ERGA and how that aligns with global efforts particularly those containing the majority of the planets remaining biodiversity; disparities in resources and capacity (personnel, financial, and infrastructural); balancing decentralisation and innovation with standardisation, reproducibility and consistency; a need for more long-term and consistent training opportunities and disproportionate interest; and protocols, research and investment in species that are underrepresented in public data repositories.

As ERGA progresses, now with a dedicated funding stream through Biodiversity Genomics Europe, it can now build upon, learn and make the intentional investments needed to address at least some of these challenges. Although a centralised source of funding to support these endeavours is overall a positive it will also provide many challenges concerning diversity and equity, however efforts are underway to safeguard at least some level of the decentralised process e.g., community sampling and hotspot sequencing.

## Supporting information

ExtendedData_Figures

Abstract_Translations

## Acknowledgements

**<u>ERGA Infrastructure:</u>** We acknowledge access to the storage resources at Barcelona Supercomputing Center. We would like to thank Alisha Ahamed, Josephine Burgin, Joana Paupério, Jeena Rajan and Guy Cochrane from the European Nucleotide Archive (ENA) for their support regarding data coordination and submission. **<u>ERGA Hubs:</u>** We thank the Antwerp University Hospital Center of Medical Genetics and Jarl Bastianen for access to sequencing library quality control equipment. **<u>Commercial Partners:</u>** We would like to acknowledge and thank all supplier partners that have kindly donated kits, reagents to the ERGA pilot Library Preparation Hubs to support species without funding to produce the generation of high-quality genomes and annotations. This support has been key to embedding a culture of diversity, equity, inclusion, and justice in the Pilot Project. Specifically we want to thank Dovetail Genomics, Part of Cantata Bio LLC, especially Mark Daly, Thomas Swale and Lily Shuie; Arima Genomics; PacBio; Integrated DNA Technologies (IDT); MagBio Genomics Europe GmbH; Zymo Research; Agilent Technologies; Fisher Scientific Spain; Illumina Inc. <u>**ERGA sequencing partners:**</u> We would like to thank all the sequencing facilities involved in the project: i) the Integrative Omics platform of the Italian Node of ELIXIR, the European Research Infrastructure for Life Science data, Giovanna Longo for support in library preparation and Apollonia Tullo for the management of the Integrative Omics platform (Italy); ii) the France Génomique network; iii) the DFG-funded NGS Competence Center Tübingen;; iv) SciLifeLab Genomics / National Genomics Infrastructure/Uppsala Genome Center and UPPMAX for aiding in High Molecular Weight DNA/RNA extraction, massive parallel sequencing and computational infrastructure; v) we would like to thank all the teams at Wellcome Sanger institute Tree of life programme, namely the ToL Samples management team, the ToL Core Laboratory, the support of Sanger Scientific Operations, especially the Long Read teams, the Tree of Life Assembly team and the Genome Reference Informatics Team, and the Tree of Life ERGA-Pilot project management team, especially Theodora Anderson.; vi) The Earlham Institute by members of the Genomics Pipelines and Core Bioinformatics Groups. This work was also supported by the Scientific Computing group, as well as support for the physical HPC infrastructure and data centre delivered via the NBI Research Computing group; vii) the Functional Genomics Center Zurich (FGCZ) for library preparation and sequencing; viii) the NGSP team at the University of Bern, from DNA and RNA extraction to data generation on both long read and short-read platforms; ix) The Lausanne University Genomic Technologies Facility (GTF, Switzerland, https://wp.unil.ch/gtf/) for library preparation and sequencing; x) the DRESDEN concept genome Center, part of the MPI-CBG and the technology platform of the CMCB at the TU Dresden; xi) the DFG Research Infrastructure West German Genome Center. NGS analyses were carried out at the production sites Cologne, Bonn and Düsseldorf; xii) We would like to acknowledge the material support through the Oxford Nanopore Technologies ORG.one project in the execution of sequencing the *Acipenser sturio* genome at the University of Wageningen (Netherland) xiii) the Centro Nacional de Análisis Genómico CNAG; the Catalan Initiative of the Catalan Biogenome Project; xiv) Computational resources were provided by the HPC core facility CalcUA of the University of Antwerp and VSC (Flemish Supercomputer Center), and the Biomina network. **<u>ERGA Training and Knowledge Transfer partners:</u>** The 2022 EMBO Practical Course “Hands-on course in genome sequencing, assembly and downstream analyses” at the Université libre de Bruxelles. **<u>Co-authors</u>**: L.R would like to thank the Naturhistorisches Museum Bern; D.D.P is funded by Biodiversity Genomics Europe (BGE); The tissue sample for the European mink has been taken from a male mink in Dordogne (France) in 2006 by Pascal Fournier and Christine Fournier-Chambrillon (GREGE, sample collectors), in the framework of the first National Action Plan for the conservation of the European mink, and sent to a collection of cryopreserved cells (managed by Vitaly Volobouev) at MNHN (National Museum of Natural History). This cryopreserved sample was used by Bertrand Bed’Hom (sample ambassador) to generate fresh cell cultures for the production of high-quality DNA by Genoscope, and the construction of a reference genome. The French institutional partners for the European mink conservation program in France are DREAL (Direction régionale de l’environnement, de l’aménagement et du logement Nouvelle-Aquitaine) and OFB (Office Français de la Biodiversité). **<u>ERGA Community</u>:** We would like to thank: Svein-Ole Mikalsen and Sunnvør í Kongsstovu (University of Faroe Islands) for the providing *Argentina silus* and *Ammodytes marinus* to the pilot project as a of the project Genome Atlas of Faroese Ecology; Michel Sartori (Musée cantonal des sciences naturelles, Lausanne); Karim Gharbi, Suzanne Henderson, Kendall Baker, Tom Barker, Naomi Irish, Jamie McGowan, Will Nash, James Lipscombe, Angela Man, Alex Durrant, Mariano Olivera, Chris Watkins, Jonathan Wright, David Swarbreck, Neil Shearer, Sacha Lucchini, Thomas Brabbs, Vanda Knitlhoffer, Leah Catchpole, Fiona Fraser, Seanna McTaggart (Earlham Institute, Norwich, UK), W.N extracted high molecular weight DNA from *Xylocopa violacea* individuals, conducted assembly of the resulting PacBio HiFi data, coordinated generation of Hi-C data for this species, and manually curated the scaffolded assembly. LIMS support using SapioLIMS; Lada Jovović (Ruđer Bošković Institute, Croatia); Federica Montesanto and Francesco Mastrototaro (Università degli Studi di Bari “A. Moro” Dipartimento di Bioscienze, Biotecnologie e Ambiente), for contributing to the design of the Pilot project on two *Botryllus* species; Jana Bedek (Ruđer Bošković Institute, Croatia); João Jacinto, Helena Trindade, Manuela Sim-Sim (cE3c - Centre for Ecology, Evolution and Environmental Changes, Faculdade de Ciências da Universidade de Lisboa & CHANGE - Global Change and Sustainability Institute, Portugal), M.S.S is Sample Ambassador for *Corema album*; Christian de Guttry, Julien Marquis (University of Lausanne, Switzerland); Native Flora Centre of Lombardy (Centro Flora Autoctona della Lombardia, CFA), c/o Parco Monte Barro, Galbiate (LC) Italy); Reichlin Pascal; Yannick Chittaro (info fauna); Ferran Palero ([Catalan Biogenome Project](https://www.biogenoma.cat/en/who-we-are/)); Sara Vicente and Cristina Máguas (Centre for Ecology, Evolution and Environmental Changes (cE3c) & CHANGE - Global Change and Sustainability Institute, Portugal; Escola Superior de Saúde Ribeiro Sanches (ERISA), IPLUSO - Instituto Politécnico da Lusofonia, Portugal); Maria Judite Alves (Museu Nacional de História Natural e da Ciência, Universidade de Lisboa); Christian Harrison, Dana R. MacGregor (Rothamsted Research) are working with the Earlham Institute to sequence, curate, and annotate an *Alopecurus aequalis* genome; Patrik Rödin Mörch (Department of Ecology and Genetics - Animal Ecology, Uppsala University); Björn Marcus von Reumont (Goethe University Frankfurt, Institute of Cell Biology and Neuroscience, Applied Bioinformatics Group); Veronique Decroocq (University of Bordeaux, INRAE), Provider of data and samples of *Prunus brigantine*, Managing sequencing effort on European *Prunus* species; Lucia Manni, Università di Padova, Dipartimento di Biologia, for contributing to the design of the Pilot project on two *Botryllus* species; Evan G. Williams (Luxembourg Centre for Systems Biomedicine, University of Luxembourg); Julia Pawłowska (Institute of Evolutionary Biology, Faculty of Biology, University of Warsaw, Poland); Craig R Primmer (University of Helsinki); Alicja Okrasińska (Institute of Evolutionary Biology, Faculty of Biology, University of Warsaw); Annamaria Giorgi (Centre of Applied Studies for the Sustainable Management and Protection of Mountain Areas-CRC Ge.S.Di.Mont, University of Milan, 25048 Edolo, Italy; Department of Agricultural and Environmental Sciences-Production, Landscape and Agroenergy-DiSAA, University of Milan, 20133 Milan, Italy); Simon Pierce (Department of Agricultural and Environmental Sciences, University of Milan); Shai Meiri (The Steinhardt Museum of Natural History, School of Zoology, Tel Aviv University); Virginia Vanni (Department of Biological and Medical Sciences, Oxford Brookes University, Oxford, OX3 0BP, United Kingdom); M.M.R. was funded by FCT and AKDN through the project CVAgrobiodiversity/333111699 and (LEAF/ISA) UID/AGR/04129/2020; Akira Peters (Bezhin Rosko, Santec, France) isolated from nature and donated the strain of *Phaeosaccion multiseriatum;* We would finally like to thank Loriano Ballarin and Fabio Gasparini, Università di Padova, Dipartimento di Biologia, for contributing to the design of the Pilot project of two Botryllus species. We would also like to acknowledge and thank Shane Whelan for his efforts in abstract translation.

## Glossary

**Biodiversity genomics** - The application of genomic methods to research biodiversity.

**BUSCO** - A bioinformatic method (Benchmarking Universal Single-Copy Orthologues) used to estimate the completeness of the coding fraction of an organism’s genome based on the proportion of (lineage specific) single copy orthologous genes that are found in a genome assembly ^52^.

**INSDC** - International Nucleotide Sequence Database Collaboration (https://www.insdc.org/) is an initiative between the DDBJ, EMBL-EBI and NCBI that together act as a global repository of sequence data and associated metadata, and provide tools and services that allow access to genomic resources.

**Reference genome** - An accepted standard representation of an organism’s DNA sequence. High-quality reference genomes typically have high completeness (chromosome-level with few gaps in sequence), few errors, and are annotated and accessible. A reference genome serves as a tool for alignment-based analyses, such as variant calling or RNAseq, and has many other applications, for example, phylogenetics and evolutionary relationships, identification of genes and variants, functional analysis and comparative genomics. Reference genomes referred to as “drafts” are those that are under active construction and refinement, and not yet finalised through manual curation.

**Genomic resource -** A genomic resource, for the purpose of this manuscript, refers to a reference genome, genome annotation, voucher specimen, cryopreserved sample and comprehensive metadata.

**FAIR Principles -** A set of principles to guide appropriate management and curation of scientific data (https://www.go-fair.org/fair-principles/) that emphasise data accessibility and use by ensuring that data are Findable, Accessible, Interoperable, and Reusable. Due to the increasing amount of scientific data being reposited, FAIR guidelines promote a data format that is amenable to automated computational access of data by stakeholders^64^.

**CARE Principles -** The CARE principles for Indigenous data governance (https://www.gida-global.org/care) provide a governance framework that supports the recognition of rights and interests Indigenous Peoples’ to their physical and digital data as well as their Indigenous Knowledges^65^.

**Metadata -** A collection of data that provides contextual information about multiple characteristics of other, corresponding original data.

**Voucher -** A voucher specimen is a permanently preserved object (either whole or in part, and/or physical or digital) of an identified organism (verified by a recognised expert) and which is deposited in an accessible facility or database. A voucher provides physical evidence about any specimen’s taxonomic identity^14^. Voucher deposition is a best practice for conducting biodiversity genomics research.

(Genome) **annotation -** The process of identifying the functions of different pieces of a genome. This includes genes that code for proteins and non coding features (e.g. intron-exon structure of protein coding genes, promotors, transposable elements). Typically performed using computational methods, followed by manual curation.

(Genome) **completeness -** An estimate of how well a reference genome represents the complete sequence of the target organism. A complete genome should equal the haploid genome size of the target, but may be defined when ‘*all chromosomes are gapless and have no runs of 10 or more ambiguous bases, there are no unplaced or unlocalized scaffolds, and all expected chromosomes are present.*’ (https://www.ncbi.nlm.nih.gov/assembly/). There are different approaches to estimate the completeness, like BUSCO, analysing K-mers, etc.

**Library -** DNA, cDNA, or RNA that has been prepared for NGS within (usually) a specific size range and containing adapters, which are designed to be appropriate for (a) specific sequencing platform(s).

(Genome) **assembly -** A genome assembly is a representation of an organism’s genome that is made using computer programs to turn (assemble) raw sequence data into longer, continuous sequences.

**PUID -** A permanent unique identifier is a unique label for an object that does not change, such as the Digital Object Identifier (DOI) attached with a scientific publication.

**ENA -** The European Nucleotide Archive (https://www.ebi.ac.uk/ena) is a global repository for sequence data and provides resources that support management and access to sequence data.

**Equity Deserving** - According to the Canadian Council (https://canadacouncil.ca/glossary/equity-seeking-groups) equity deserving groups are those individual researchers, communities, Peoples, regions or countries that have identified barriers to equal access, opportunities, and resources due to disadvantage and/or discrimination and that are actively seeking, and deserving of social justice and reparation. The discrimination experienced could be caused by attitudinal, historic, social, and environmental barriers that could be based on a plethora of characteristics that are including (but not limited to) sex, age, ethnicity, disability, economic status, gender, gender expression, nationality, race, sexual orientation, and creed.

**COPO -** The Collaborative OPen Omics (COPO) platform is for researchers to publish their research assets, providing metadata annotation and deposition capability. It allows researchers to describe their datasets according to community standards and broker the submission of such data to appropriate repositories whilst tracking the resulting accessions/identifiers^29^.

**Open data -** Open data are freely accessible and unrestricted data that can be accessed, used, reused and shared with third parties for any purpose.

**HSM -** Hierarchical Storage Management is both a data management and data storage technique which transparently manages the movement of data between the different layers of a tiered storage based on file size thresholds, usage and I/O pressure. Usually, a tiered storage is composed of one or more layers of disk arrays, ordered by capacity, latency, redundancy and storage cost. A slow but economically effective archival layer is at the bottom, composed of magnetic tape libraries and automated tape robots, with the highest capacity and latency. The movement between layers is automatically triggered.

**ONT -** Oxford Nanopore Technologies (ONT; https://nanoporetech.com/) is a next generation sequencing technology whereby sequence data are generated from the changes in current that occur as single-stranded DNA or RNA molecules pass through nanoscale protein pores (nanopores). ONT provides long read data (up to several megabases) that facilitate genome assembly^66,67^.

**PacBio -** Pacific Biosciences (PacBio; https://www.pacb.com/) is a single-molecule, real time (SMRT) next generation sequencing technology in which sequence data are generated by fluorescent light emission that occurs when a DNA polymerase adds nucleotides. PacBio produces long read data (tens of kilobases) that facilitate genome assembly.

**HiFi reads -** HiFi (High Fidelity) PacBio reads are produced by taking multiple sequences of the same molecule to provide a consensus sequence that is usually 12-20kbp long and has a low error rate (>99.9 % consensus accuracy)^68^.

**Hi-C -** Sequencing-based method used to study three-dimensional interactions among chromatin regions by measuring the frequency of contact between pairs of loci. Since contact frequency is related to the distance between a pair of loci, Hi-C linking information is used to help with scaffolding stages during a genome assembly process.

**Hi-C map / graph production -** The occurrence and frequency of Hi-C contacts are analysed and used in assembly scaffolding. They are typically visualised in Hi-C 2D heatmaps with the full genome sequence on the X and Y axis and a markup for each observed contact.

**Omni-C -** Modified version of Hi-C that uses a sequence-independent endonuclease during its protocol to produce more even sequence coverage increasing overall resolution.

**RNA-Seq -** RNA-Seq is a technique that determines the complete or partial RNA sequence using NGS. The RNA expression profiles vary in different tissues of the same organism and can be influenced by physiopathological circumstances. RNA-Seq data facilitate genome assembly by providing empirical evidence for annotation of transcribed regions^69^.

**IsoSeq -** This is a sequencing protocol developed by PacBio that aims to sequence full-length transcripts using the accurate, long read capabilities of PacBio HiFi technology. IsoSeq data facilitate analysis of transcriptomes and genome annotation by identifying full-length isoforms of transcripts.

**Haplotype -** A haplotype refers to the collection of genetic material within an organism that is inherited together. Haplotype may be used to describe a few loci or any number of chromosomes (a chromosome-scale haplotype).

**K-mer -** A K-mer is a DNA sequence of length k; for example, the sequence AGCT contains the 3-mers (K-mers of length 3) AGC and GCT.

**Transcriptome -** A transcriptome is a set of aligned RNAseq reads representing RNA collected from a sample or collection of samples. This includes both protein-coding and non-coding transcripts. For the ERGA Pilot Project, poly-A+ transcripts were profiled.

**Interested Parties -** This term, for the purposes of this manuscript refers to the range of external stakeholders (e.g., commercial companies, policymakers etc) and rights holders (e.g., Indigenous Peoples) that have an interest in biodiversity genomics research.

**EBP Genome assembly quality standard 6..Q40 -** Minimum reference standard of 6.C.Q40, i.e. megabase N50 contig continuity and chromosomal scale N50 scaffolding, with less than 1/10,000 error rate. For species with chromosome N50 smaller than a megabase this will be C.C.Q40. Additional recommendations include K-mer completeness >90%, BUSCO complete single-copy single >90%, BUSCO complete single duplicate < 5%, and Gaps/Gbp <1000.

**Widening Country -** Widening countries are countries with low participation rates in FP7 and H2020 projects (low level of investment into research and innovation (R&I)). According to the Horizon Europe [regulation](https://rea.ec.europa.eu/news/eu-committed-research-and-innovation-through-horizon-europe-widening-programme-2022-08-29_en) the Widening countries are: Bulgaria, Croatia, Cyprus, Czech republic, Estonia, Greece, Hungary, Latvia, Lithuania, Malta, Poland, Portugal, Romania, Slovakia, Slovenia and all associated countries with equivalent characteristics in terms of R&I performance and the Outermost Regions.

