## ExtendedData_Figures for "The European Reference Genome Atlas: piloting a decentralised approach to equitable biodiversity genomics"

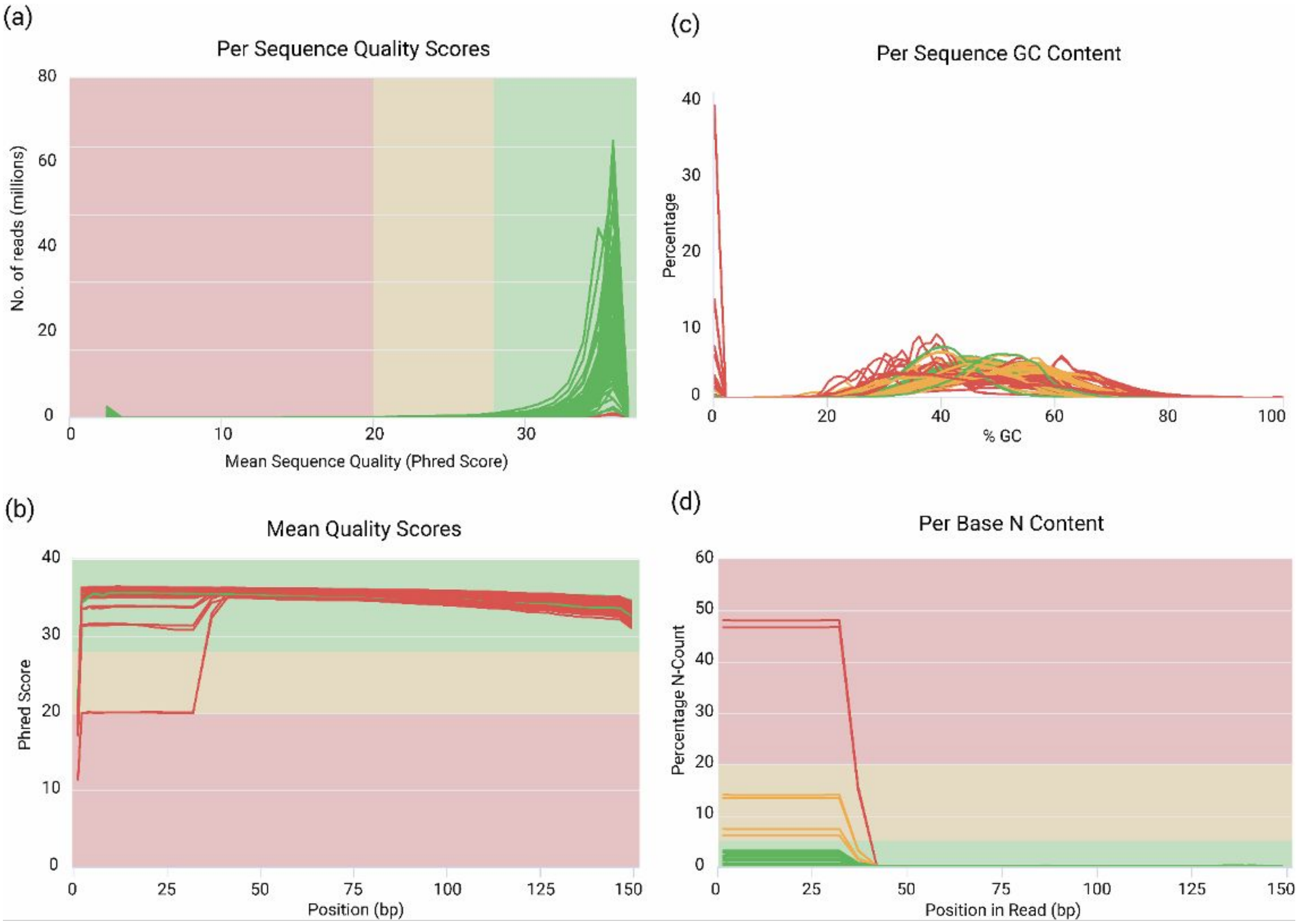


**Extended Data Figure 1:** Quality Assessment of RNA-seq Data . (*a*) Per Sequence Quality Scores: Distribution of quality scores across all sequences, with the y-axis representing the number of reads in millions. (*b*) Mean Quality Scores: The average quality score at each position in a read. (*c*) Per Sequence GC Content: The GC content distribution across all sequences. (*d*) Per Base N Content: The proportion of N content at each base position in the reads, reflecting ambiguous base calls.

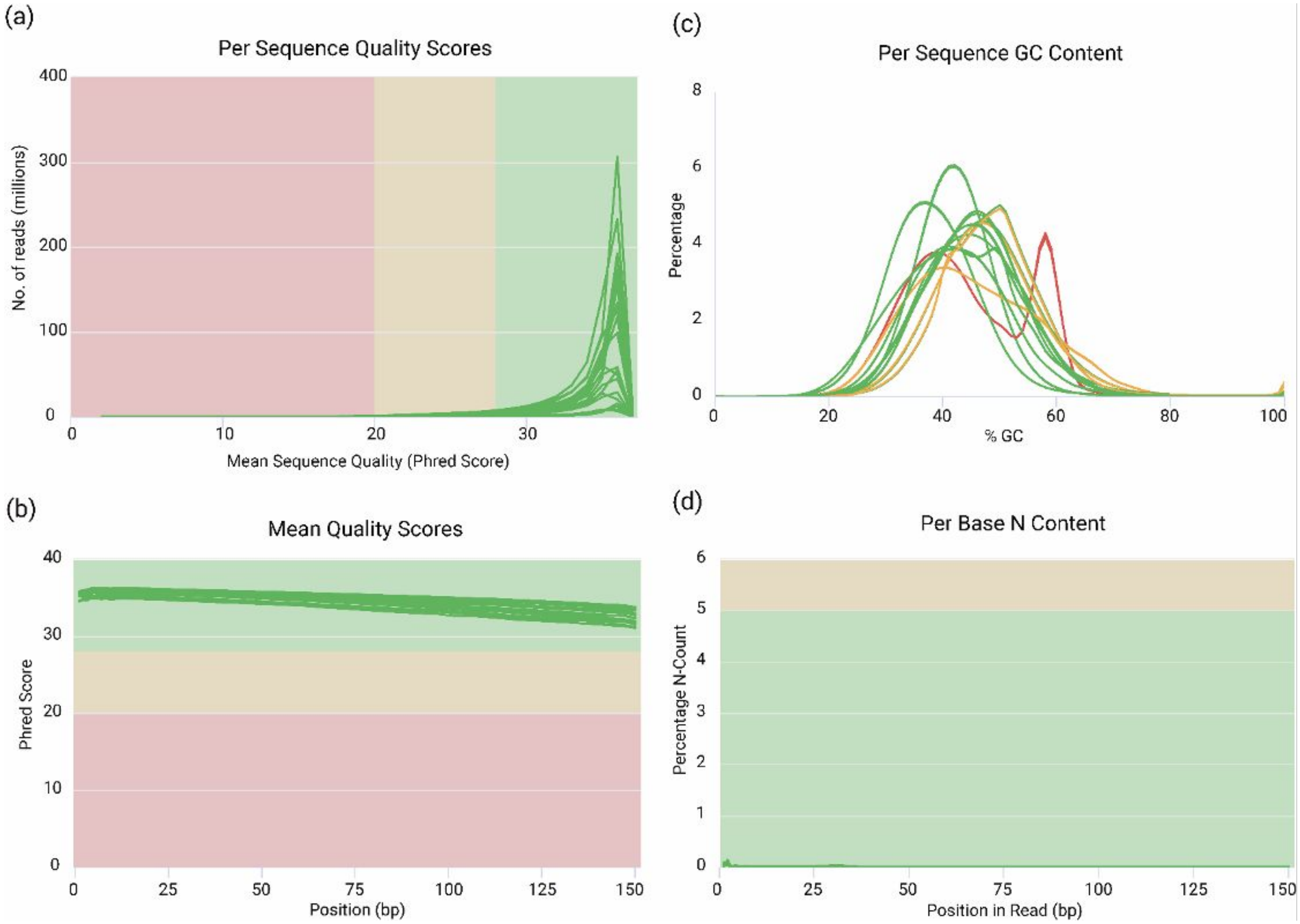


**Extended Data Figure 2:** Quality Assessment of Hi-C Sequencing Data. (*a*) Per Sequence Quality Scores: Distribution of quality scores across all sequences, with the y-axis representing the number of reads in millions. (*b*) Mean Quality Scores: The average quality score at each position in a read. (*c*) Per Sequence GC Content[HL2] : The GC content distribution across all sequences. (*d*) Per Base N Content: The proportion of N content at each base position in the reads, reflecting ambiguous base calls.

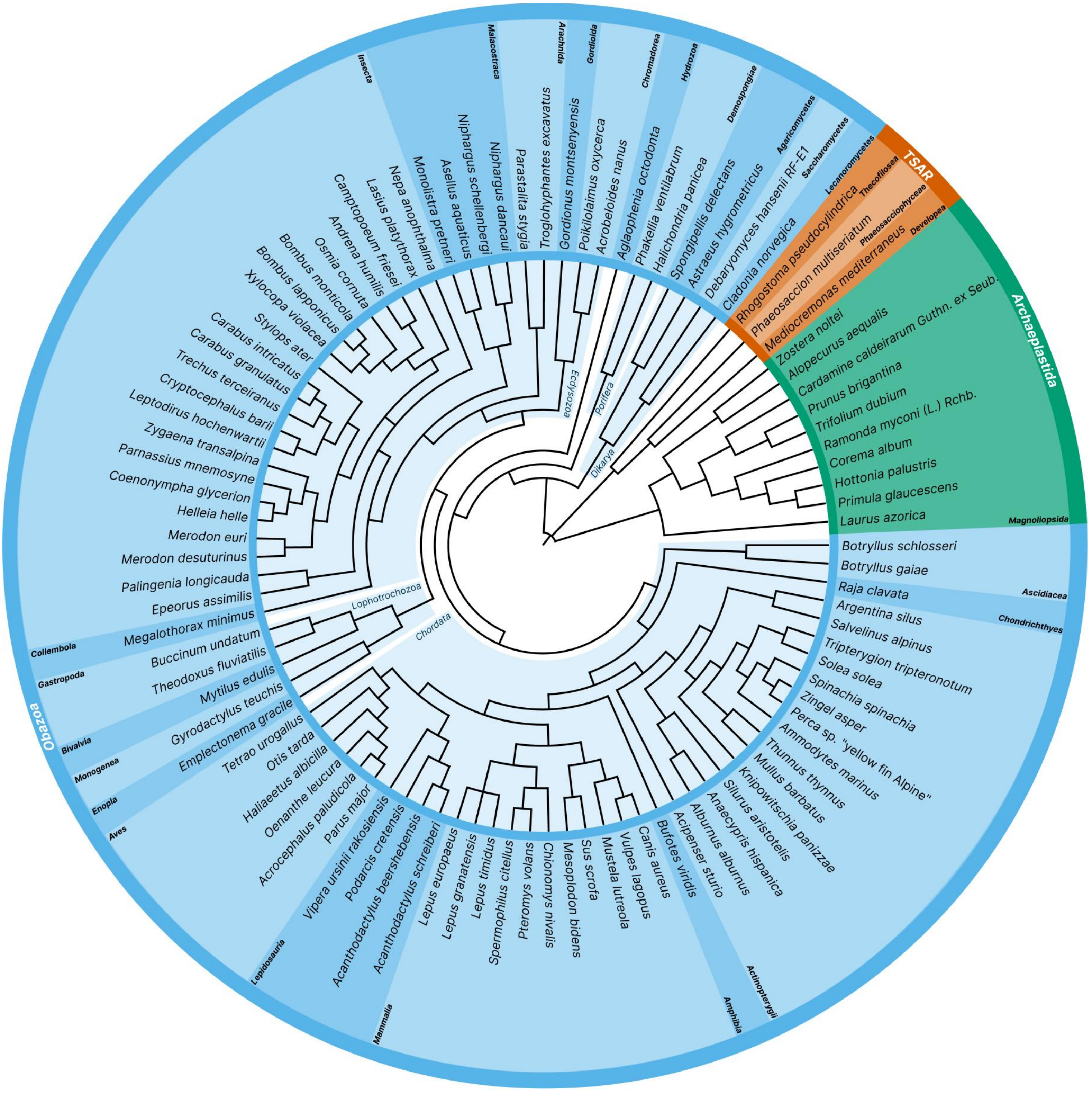


**Extended Data Figure 3:** Phylogenetic tree of the species included within the implementation test for the distributed genomic infrastructure**.**

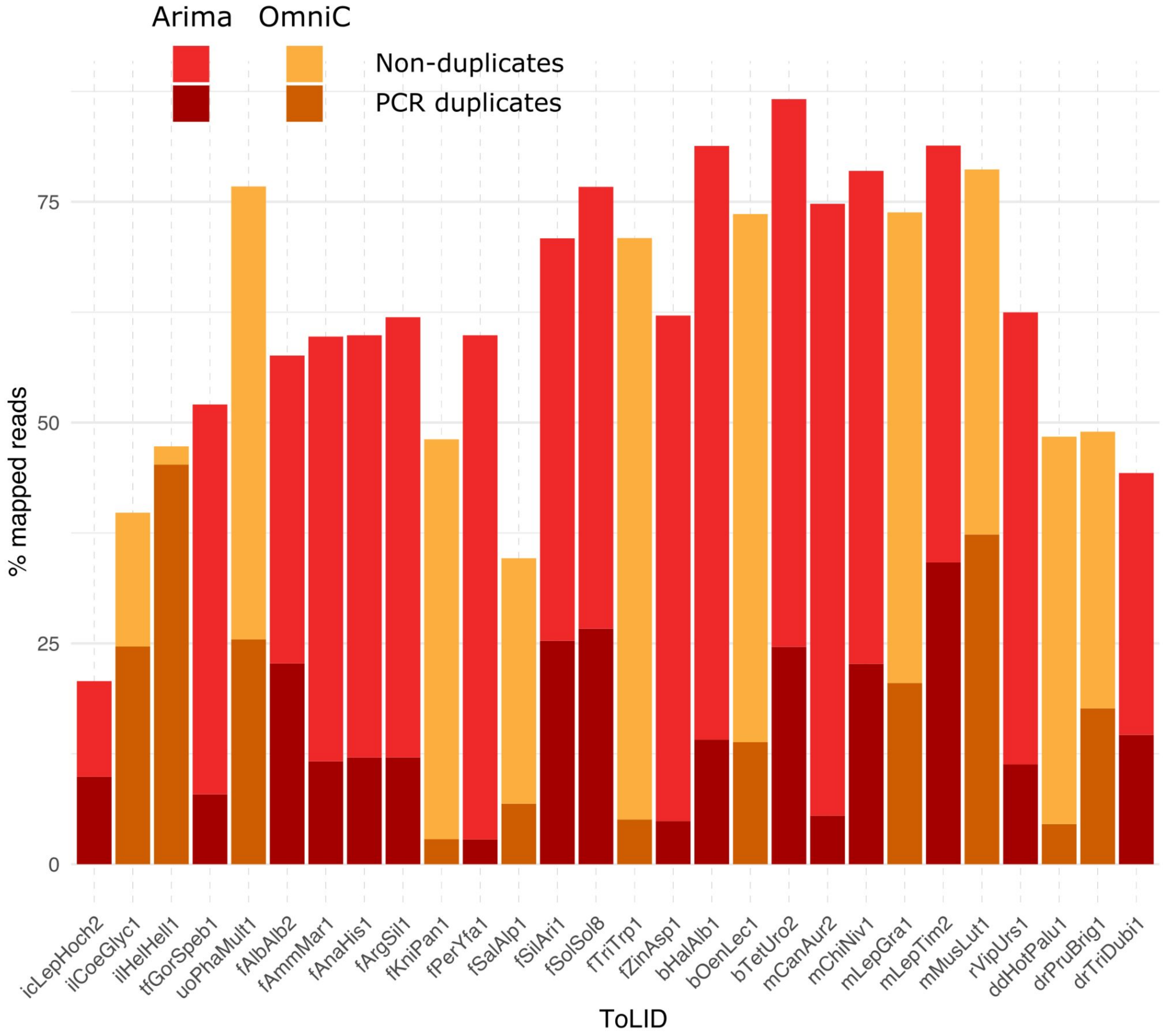


**Extended Data Figure 4:** Hi-C reads mapping metrics for completed (pre-curation and curated) assemblies according to kits used (Arima or OmniC).

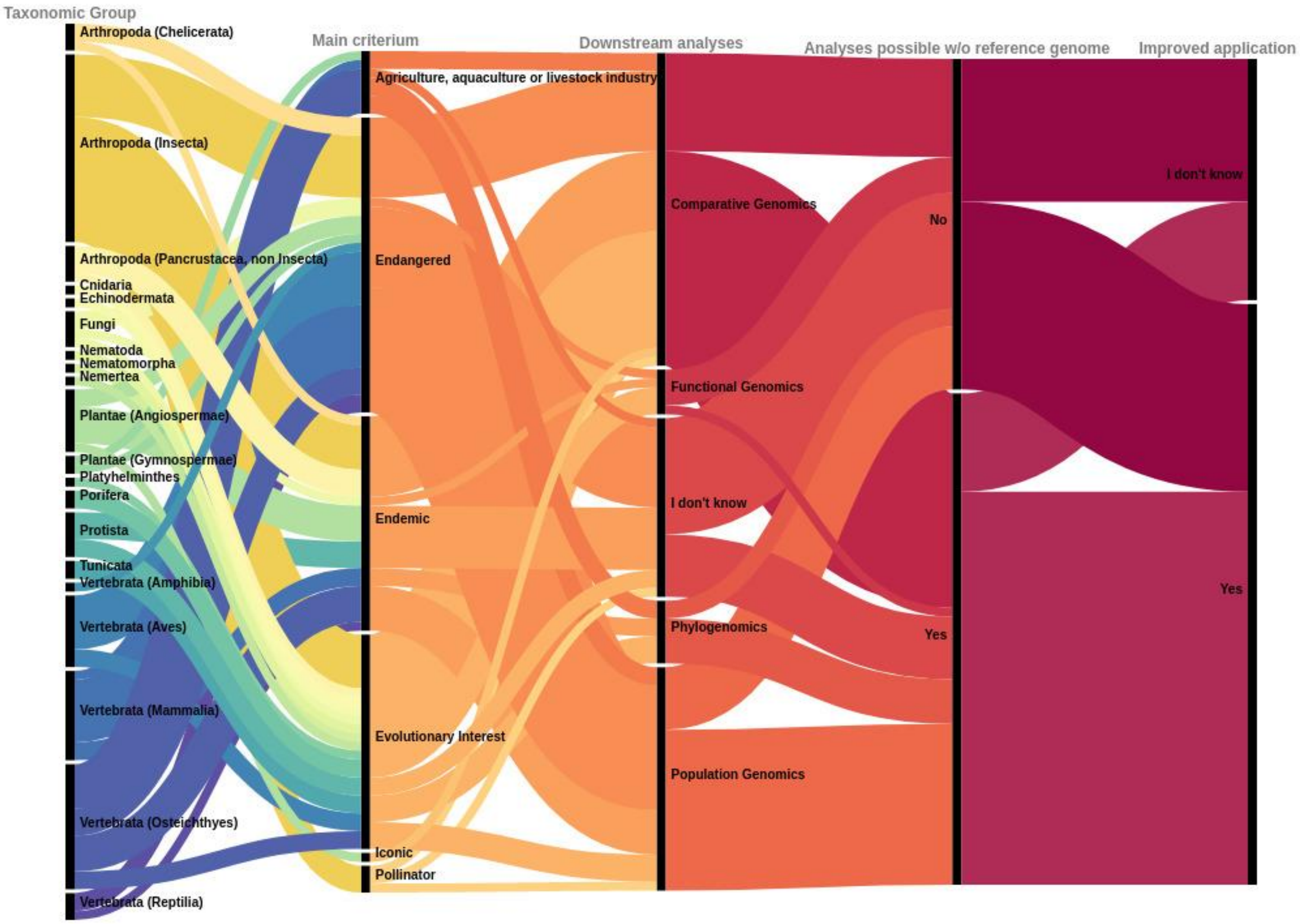


**Extended Data Figure 5: Summary of genome team data analysis survey responses.**

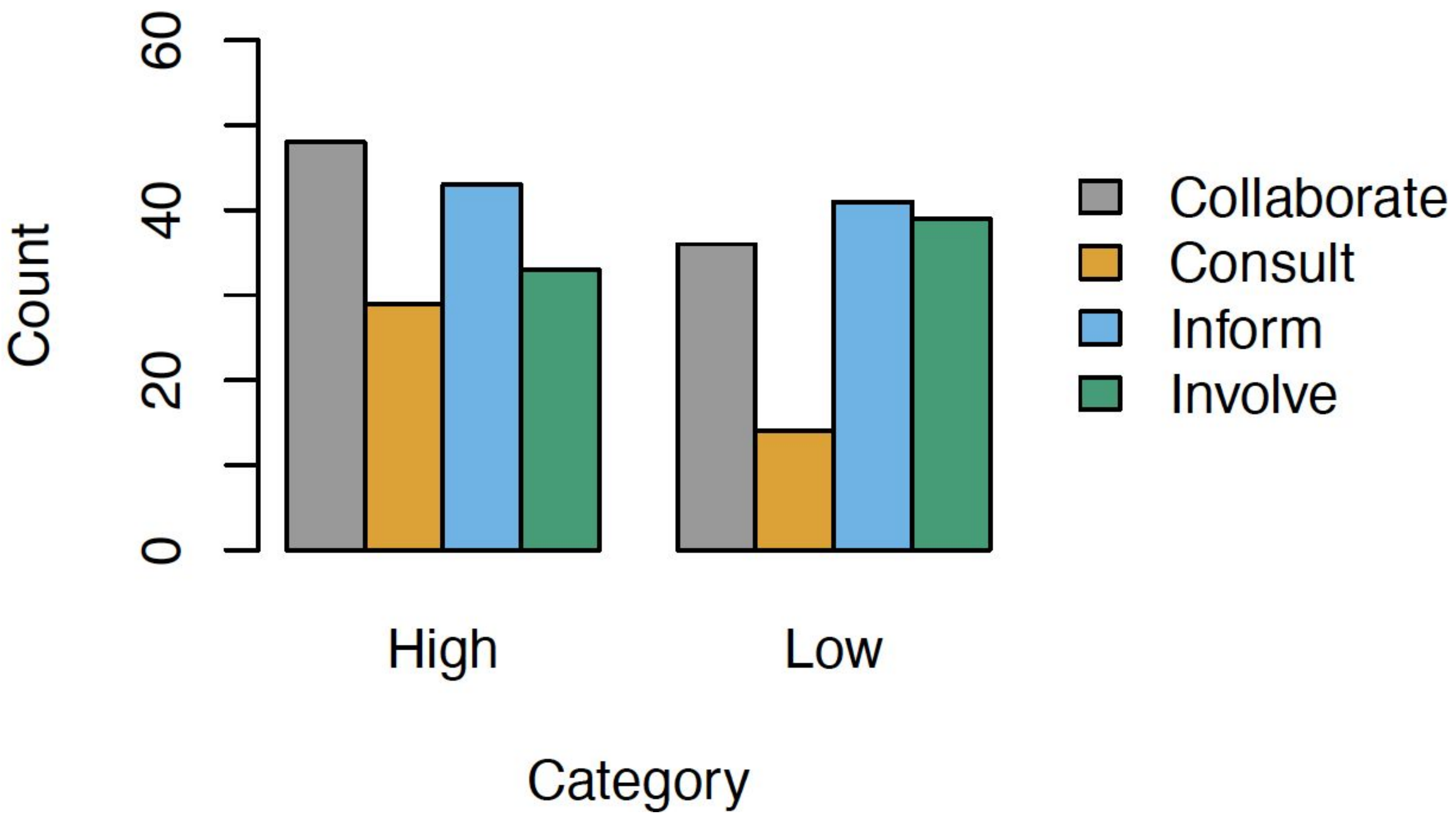


**Extended Data Figure 6:** The differences in how participants from countries with a GBARD higher than 1000 MM (High) and those with a GBARD lower than 1000 MM (Low) divided interested parties into the four categories of engagement.

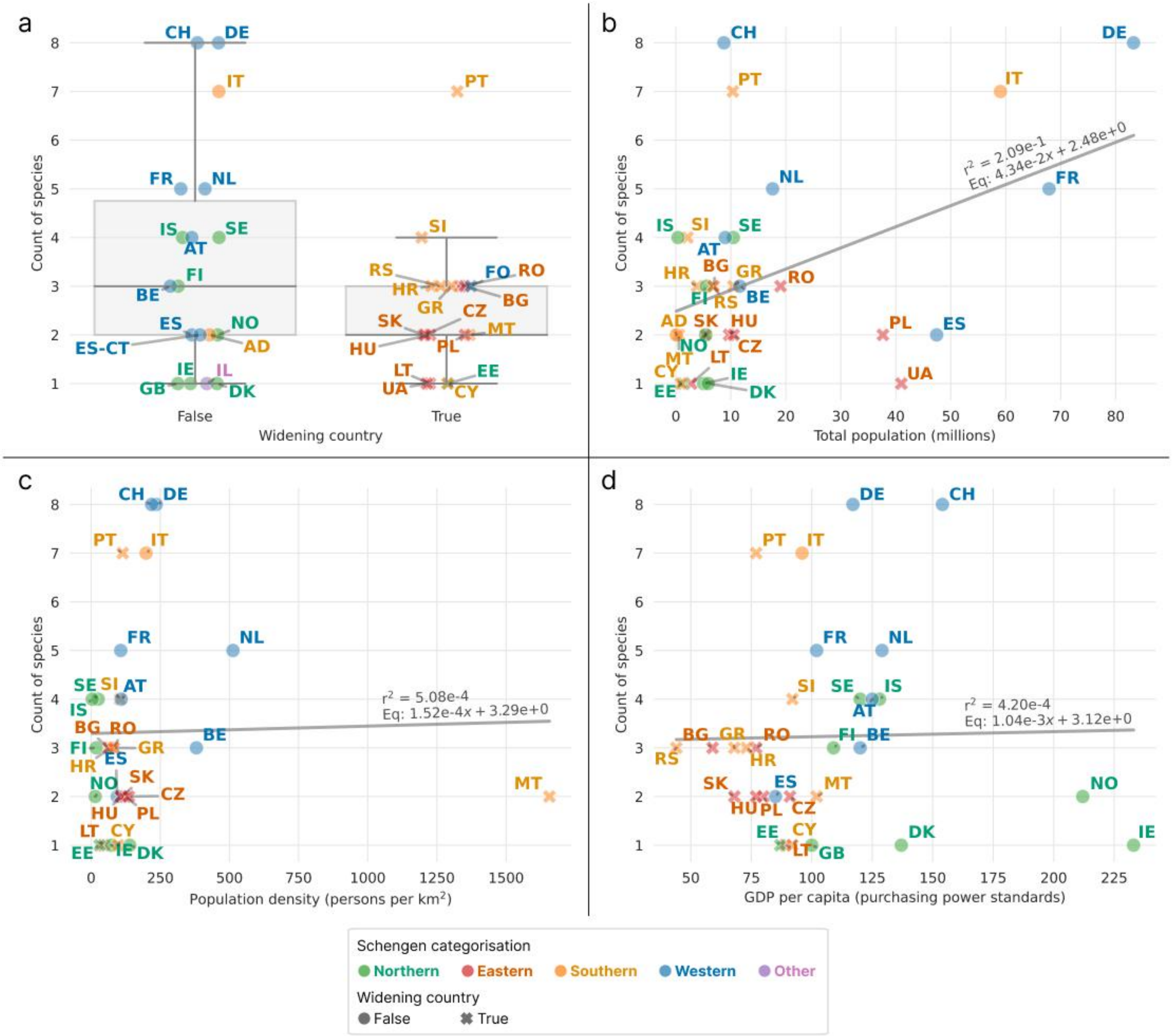


**Extended Data Figure 7.** Relationship between number of species attributed per country and country characteristics. Distribution of the number of species attributed per country: (a) Categorized as non-Widening or Widening; (b) Total population; (c) Population density; (d) Gross Domestic Product (GDP) per capita in purchasing power standards. Data points labelled with ISO country codes. Country-level data for population, population density and GDP per capita in PPS, as sourced from Eurostat (2022).

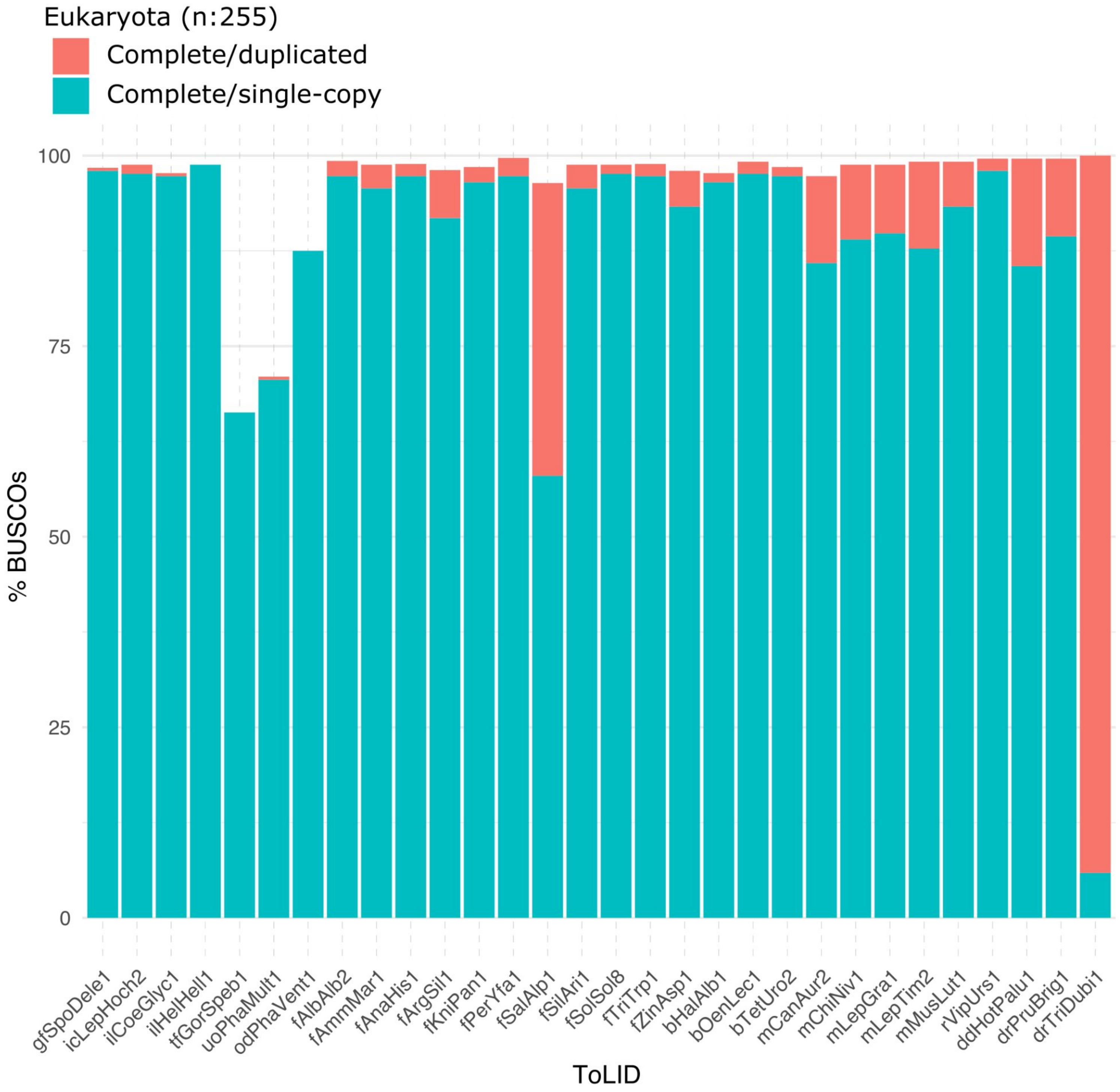


**Extended Data Figure X:** BUSCO completeness scores (single and duplicated) for completed (pre-curation and curated) assemblies using "Eukaryota" orthologs database.

**Extended Data Figure X: Sequencing QC metaanalysis of data produced in Florence.**
<Await contribution from Henrique>

**OTHER - MAY NOT BE USED**

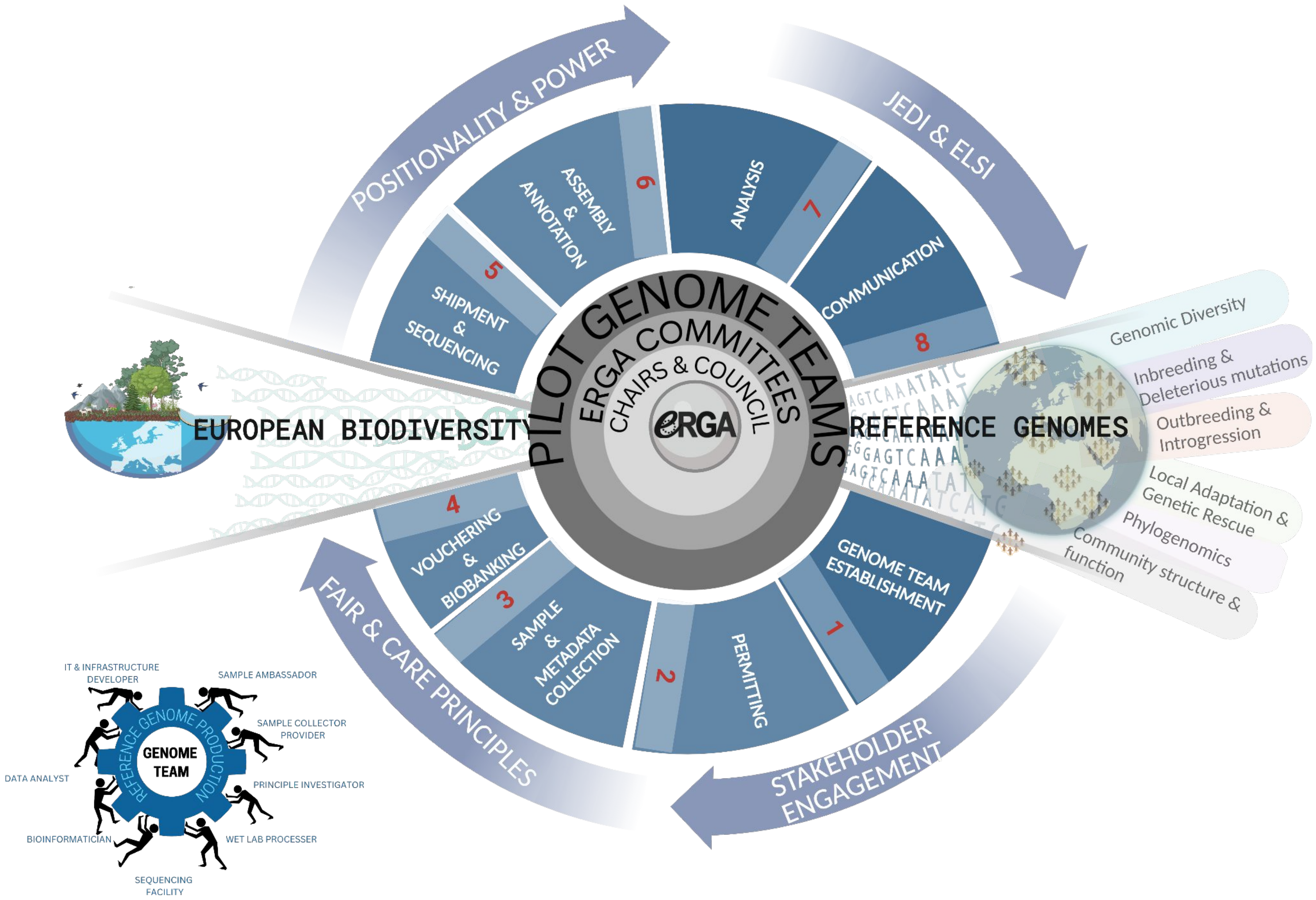


**Extended Data Figure X:** Establishing an inclusive, accessible, distributed and pan-European genomic infrastructure that could support the streamlined and scalable production of genomics resources for all European species.
