## Supplementary material for "The European Reference Genome Atlas: piloting a decentralised approach to equitable biodiversity genomics": Abstract_Translations

English:

A global genome database of all of Earth's species diversity could be a treasure trove of scientific discoveries. However, regardless of the major advances in genome sequencing technologies, only a tiny fraction of species have genomic information available. To contribute to a more complete planetary genomic database, scientists and institutions across the world have united under the Earth BioGenome Project (EBP), which plans to sequence and assemble high-quality reference genomes for all ~1.5 million recognized eukaryotic species through a stepwise phased approach. As the initiative transitions into Phase II, where 150,000 species are to be sequenced in just four years, worldwide participation in the project will be fundamental to success. As the European node of the EBP, the European Reference Genome Atlas (ERGA) seeks to implement a new decentralised, accessible, equitable and inclusive model for producing high-quality reference genomes, which will inform EBP as it scales. To embark on this mission, ERGA launched a Pilot Project to establish a network across Europe to develop and test the first infrastructure of its kind for the coordinated and distributed reference genome production on 98 European eukaryotic species from sample providers across 34 European countries. Here we outline the process and challenges faced during the development of a pilot infrastructure for the production of reference genome resources, and explore the effectiveness of this approach in terms of high-quality reference genome production, considering also equity and inclusion. The outcomes and lessons learned during this pilot provide a solid foundation for ERGA while offering key learnings to other transnational and national genomic resource projects.

French:

Une base de données génomiques mondiale regroupant toute la diversité des espèces de la Terre pourrait constituer un trésor de découvertes scientifiques. Cependant, malgré les avancées majeures des technologies de séquençage du génome, seule une infime partie des espèces dispose d'informations génomiques. Afin de contribuer à la constitution d'une base de données génomiques planétaires plus complète, des scientifiques et des institutions du monde entier se sont unis dans le cadre du Earth BioGenome Project (Projet BioGénome de la Terre, EBP), qui prévoit de séquencer et d'assembler des génomes de référence de haute qualité pour l'ensemble des quelque 1,5 million d'espèces eucaryotes connues. Alors que l'initiative passe à la phase II, au cours de laquelle 150 000 espèces doivent être séquencées en seulement quatre ans, la participation mondiale au projet sera essentielle à sa réussite. Branche européenne de l'EBP, l'European Reference Genome Atlas (Atlas Européen de Génomes de Référence, ERGA) cherche à mettre en œuvre un nouveau modèle décentralisé, accessible, équitable et inclusif de production de génomes de référence de haute qualité, et transmettra les informations à l'EBP au fur et à mesure de sa progression. Pour se lancer dans cette mission, l'ERGA a lancé un projet pilote visant à établir un réseau à travers l'Europe afin de développer et de tester la première infrastructure de ce type pour la production coordonnée et distribuée de génomes de référence sur 98 espèces eucaryotes européennes à partir d'échantillons provenant de 34 pays européens. Nous décrivons ici le processus et les défis rencontrés lors du développement d'une infrastructure pilote pour la production de ressources génomiques de référence, et explorons l'efficacité de cette approche en termes de production de génomes de référence de haute qualité, en tenant compte également de l'équité et de l'inclusion. Les résultats et les enseignements tirés de ce projet pilote constituent une base solide pour l'ERGA, tout en offrant des enseignements clés à d'autres projets transnationaux et nationaux visant à établir de nouvelles ressources génomiques.

German:

Eine globale Genomdatenbank für die gesamte Artenvielfalt der Erde könnte eine Schatzkiste für wissenschaftliche Entdeckungen darstellen. Trotz großer Fortschritte bei den Technologien zur Genomsequenzierung liegen bislang allerdings nur für einen winzigen Bruchteil der Arten Informationen ihres gesamten Erbgutes vor. Um zu einer umfassenderen weltweiten Genomdatenbank beizutragen, haben sich Wissenschaftler*innen und Institutionen aus aller Welt im Earth BioGenome Project (EBP) zusammengeschlossen, das schrittweise die Sequenzierung und Assemblierung hochwertiger Referenzgenome für alle ca. 1,5 Millionen bekannten eukaryontischen Arten plant. Während die Initiative in Phase II übergeht, in der innerhalb von nur vier Jahren 150.000 Arten sequenziert werden sollen, wird eine weltweite Beteiligung am Projekt von grundlegender Bedeutung für den Erfolg sein. Der Europäische Referenzgenom-Atlas (ERGA) stellt den europäischen Knotenpunkt des EBP dar und soll ein neues dezentrales, leicht zugängliches, faires und integratives Modell für die Erstellung hochwertiger Referenzgenome zur Verfügung stellen, welches das EBP bei seiner Ausweitung inhaltlich unterstützen wird. Zu diesem Zweck hat ERGA ein Pilotprojekt für ein europaweites

Netzwerk gestartet und die erste Infrastruktur ihrer Art für eine koordinierte und dezentrale Produktion von Referenzgenomen für 98  eukaryontische europäische Arten entwickelt und getestet, wobei Proben durch Projektbeteiligte aus 34 europäischen Ländern geliefert wurden. In diesem Artikel werden Prozesse und Herausforderungen beschrieben, die sich bei der Entwicklung einer Pilotinfrastruktur zur Erstellung von Referenzgenomressourcen ergeben haben, sowie die Wirksamkeit dieses Ansatzes für eine qualitativ hochwertige Referenzgenom-Erstellung - unter Berücksichtigung von Fairness und Einbindung - untersucht. Die Ergebnisse und Erfahrungen aus diesem Pilotprojekt bilden eine solide Grundlage für ERGA und stellen gleichzeitig wichtige Erkenntnisse für andere transnationale und nationale Projekte zur Erarbeitung genomischer Ressourcen dar.

Greek:

Η δημιουργία μιας βάσης γονιδιωματικών δεδομένων για το σύνολο των ειδών του πλανήτη μας θα αποτελέσει μοναδικό θησαυρό από τον οποίο θα προκύψει πλήθος επιστημονικών ανακαλύψεων. Ωστόσο, παρά τη σημαντική πρόοδο στις τεχνολογίες προσδιορισμού της αλληλουχίας των γονιδιωμάτων, τα διαθέσιμα γονιδιώματα προέρχονται από πολύ μικρό ποσοστό των ειδών του πλανήτη μας. Για τη δημιουργία μιας πληρέστερης βάσης γονδιωματικών δεδομένων σε παγκόσμιο επίπεδο, επιστήμονες και ιδρύματα από όλο τον κόσμο έχουν ενώσει τις δυνάμεις τους στο πλαίσιο του Earth BioGenome Project (EBP), το οποίο σχεδιάζει σταδιακά να αλληλουχήσει και να συγκεντρώσει υψηλής ποιότητας γονιδιώματα αναφοράς για το σύνολο των περίπου 1,5 εκατομμύριων αναγνωρισμένων ευκαρυωτικών ειδών της Γης. Καθώς το έργο αυτό εισέρχεται στη δεύτερη φάση του (ΦΑΣΗ ΙΙ), κατά την οποία πρόκειται να αλληλουχηθούν τα γονιδιώματα από 150.000 είδη σε χρονικό διάστημα μόλις τέσσερα χρόνια, η συμμετοχή ερευνητών από όλο τον κόσμο κρίνεται καταλυτική για την επιτυχία του εγχειρήματος. Ο Ευρωπαϊκός Άτλας Γονιδιωμάτων Αναφοράς (European Reference Genome Atlas, ERGA) που αποτελεί τον ευρωπαϊκό κόμβο του EBP, επιδιώκει να εφαρμόσει ένα νέο αποκεντρωμένο, προσβάσιμο, δίκαιο και περιεκτικό μοντέλο για την παραγωγή υψηλής ποιότητας γονιδιωμάτων αναφοράς, το οποίο, όσο προχωράει, θα επικαιροποιεί το EBP. Για τον σκοπό αυτό, το ERGA ξεκίνησε ένα πιλοτικό έργο με στόχο τη δημιουργία ευρωπαϊκού δικτύου για την ανάπτυξη και την εφαρμογή της πρώτης, στο είδος της, υποδομής με στόχο τη συντονισμένη παραγωγή γονιδιωμάτων αναφοράς από 98 ευρωπαϊκά ευκαρυωτικά είδη. Τα δείγματα των ειδών αυτών προέρχονται από φορείς συλλογών που εδρεύουν σε 34 διαφορετικές ευρωπαϊκές χώρες. Στο άρθρο αυτό περιγράφουμε τη διαδικασία και τις προκλήσεις που αντιμετωπίσαμε κατά την ανάπτυξη αυτής της πιλοτικής υποδομής για την παραγωγή υψηλής ποιότητας γονιδιωμάτων αναφοράς, και διερευνούμε την αποτελεσματικότητα αυτής της προσέγγισης, λαμβάνοντας υπόψη επίσης τη δίκαιη συμμετοχή και την ενσωμάτωση. Τα αποτελέσματα και τα διδάγματα που αντλήθηκαν κατά τη διάρκεια αυτού του πιλοτικού προγράμματος παρέχουν μια σταθερή βάση για το ERGA ενώ παράλληλα προσφέρουν τις βασικές γνώσεις για άλλα διακρατικά και εθνικά έργα παραγωγής γονιδιωματικών υποδομών.

Irish:

D'fhéadfadh bunachar sonraí d'éagsúlacht speiceas domhanda a bheith ina thaisce d'fhionnachtana eolaíochta. Cé go bhfuil dul chun cinn ollmhór déanta i dteicneolaíocht sheiceamhú géanóm níl eolas géanómaíoch ar fáil ach do líon fíorbheag de speiceas. Tá eolaithe agus institiúid ar fud na cruinne ag comhoibriú faoi bhratach an tionscnaimh EarthBioGenome Project (EBP) ar mhaithe le bheith ag cur le bunachar sonraí géanómaíoch domhanda atá níos iomláine. Tá sé mar aidhm ag an EBP géanóim thagartha d'ardchaighdeán a sheicheamhú agus a chur le chéile do gach ceann de na ~1.5 milliún speiceas eocarótach aitheanta trí phróiseas céim ar chéim. De réir mar a bhogann an tionscnamh ar aghaidh go Céim II beidh sé riachtanach do rath an tionscnaimh go mbeidh rannpháirtíocht domhanda toisc go ndéanfar seicheamhú ar 150,000 speiceas taobh istigh de cheithre bliana. Mar lárphointe Eorpach an EBP, tá sé mar aidhm ag Atlas Géanóm Tagartha na hEorpa (ERGA) samhail atá díláraithe, inrochtana, cothromasacha agus ionchuimsitheach chur i bhfeidhm maidir le géanóim thagartha a chur ar fáil. Déanfaidh sé seo eolas a thabhairt don EBP de réir mar a mhéadaíonn sé. Le tús a chur leis an aistear seo, chuir an ERGA tús le treoirthionscadal ar mhaithe le líonra a bhunú fud fad na hEorpa. Sprioc an treoirthionscadail ná chun an chéad bhonneagar dá leithéid a fhorbairt agus a thástáil le haghaidh táirgeadh géanóim tagartha comhordaithe ar 98 speiceas eocarótach Eorpach ó sholáthraí samplacha thar 34 tír Eorpach. Déanann muid cur síos anseo ar an bpróiseas agus na dúshláin a bhaineann le bonneagar píolótach a fhorbairt ar mhaithe le hacmhainní géanóim thagartha a tháirgeadh. Anuas air sin déanfar cíoradh ar éifeachtacht an cur chuige seo maidir le táirgeadh géanóim thagartha d'ardchaighdeán le trácht déanta do chothromas agus ionchuimsitheacht. Tugann na torthaí agus na ceachtanna a d'fhoghlaimíodh i rith an phíolóta seo bunchloch láidir d’ERGA, ag an am céanna tugann sé eochairphointí foghlama go tionscadail acmhainní géanómaíochta náisiúnta agus trasnáisiúnta.

Italian:

Un database di genomi che rappresenti tutta la biodiversità globale potrebbe essere un tesoro di scoperte scientifiche. Tuttavia, nonostante gli enormi progressi nelle tecnologie di sequenziamento del genoma, solo una piccola frazione delle specie note dispone di informazioni genomiche. Per contribuire a un database genomico planetario più completo, scienziati e istituzioni di tutto il mondo si sono uniti sotto l’egida dell'Earth BioGenome Project (EBP), che prevede di sequenziare e assemblare genomi di riferimento di alta qualità per tutte le circa 1,5 milioni di specie eucariotiche note. La partecipazione mondiale al progetto sarà fondamentale per il successo della Fase II, in cui si prevede di sequenziare 150.000 specie nei prossimi quattro anni. In quanto nodo europeo dell’EBP, l’European Reference Genome Atlas (ERGA) cerca di implementare un nuovo modello decentralizzato, accessibile, equo e inclusivo per la produzione di genomi di riferimento di alta qualità per le specie europee, che aiuterà a informare gli sforzi dell’EBP. A questo scopo, ERGA ha lanciato un progetto pilota per sviluppare e testare la prima infrastruttura per la produzione coordinata e distribuita di genomi di riferimento su 98 specie eucariotiche europee da 34 paesi. Descriviamo il processo e le sfide affrontate durante lo sviluppo dell’infrastruttura ed esploriamo l’efficacia di questo approccio in termini di produzione di genoma di riferimento di alta qualità. I risultati e le lezioni apprese durante questo progetto pilota forniscono una solida base per ERGA, offrendo allo stesso tempo insegnamenti chiave ad altri progetti dedicati alla produzione di risorse genomiche nazionali e transnazionali.

Latvian:

Globāla genoma datu bāze ar informāciju par visu sugu daudzveidību, vecinātu zinātniskos atklājumus un pētniecību. Tomēr, neskatoties uz genoma sekvenēšanas tehnoloģiju ievērojamo attīstību, tikai nelielai daļai sugu ir pieejama genomiskās sekvences informācija. Lai veicinātu pilnīgāku globālo genomisko datu bāzi, zinātnieki un iestādes visā pasaulē ir apvienojušies Pasaules BioGenoma projekta (Earth BioGenome Project - EBP) ietvaros, kas, izmantojot pakāpenisku pieeju, plāno izveidot un apvienot augstas kvalitātes references genomus visām ~ 1,5 miljoniem eikariotiskām sugām. Pašreiz sākās EBP iniciatīvas otrā posma, kurā paredzēts sekvencējot 150 000 sugu genomus tikai četru gadu laikā, un visas pasaules dalība projektā būs būtiska, lai gūtu panākumus. Eiropas references genoma atlants (European Reference Genome Atlas - ERGA), kā EBP Eiropas pārstāvis plāno ieviest jaunu decentralizētu, pieejamu, taisnīgu un atvērtu modeli augstas kvalitātes references genomu ražošanai. Lai izpildītu šo misiju, ERGA izveidoja Eiropas pētniecības tīklu, izstrādāja un pārbaudīja pirmo šāda veida koordinētu un sadalītu genoma sekvenēšanas infrastruktūru un uzsāka izmēģinājuma projektu, sekvencējot 98 Eiropas eikariotisko sugu paraugus no 34 Eiropas valstīm. Šeit mēs aprakstam references genoma sekvenēšanas procesu un problēmas, kas radušās infrastruktūras izstrādes laikā, un pētām šīs pieejas efektivitāti augstas kvalitātes genoma ražošanā, ņemot vērā arī taisnīgumu atvērtību un iesaisti. Šā izmēģinājuma gaitā gūtie rezultāti un gūtā pieredze ir stabils pamats ERGA, vienlaikus piedāvājot pamatzināšanas citiem starptautiskiem un valstu genoma sekvencēšanas pētījumiem.

Lithuanian:

Pasaulinė visų Žemės rūšių įvairovės genomų duomenų bazė galėtų tapti mokslinių atradimų lobiu. Tačiau, nepaisant didžiulės genomų sekų nustatymo technologijų pažangos, šiuo metu nuskaityta tik nedidelės dalies rūšių genominė informacija. Siekdami prisidėti prie visų planetoje esančių organizmų genomų duomenų bazės kūrimo, mokslininkai ir institucijos visame pasaulyje susivienijo į Žemės biogenomo projektą (angl. Earth BioGenome Project, EBP), kuriuo planuojama susekvenuoti ir palaipsniui sukaupti aukštos kokybės etaloninius visų ~1,5 mln. pripažintų eukariotų rūšių genomus. Iniciatyvai pereinant į II etapą, kuriame per ketverius metus turi būti susekvenuota 150 000 rūšių genomų, įvairių pasaulio šalių atstovų dalyvavimas projekte bus labai svarbus sėkmei užtikrinti. Europos etaloninių genomų atlaso (ERGA) iniciatyva, kuri yra vienas pagrindinių EBP Europos centrų, siekia įgyvendinti naują decentralizuotą, prieinamą, teisingą ir įtraukų aukštos kokybės etaloninių genomų kūrimo modelį, kuriuo bus remiamasi plečiant EBP. Šiai misijai pradėti ERGA inicijavo bandomąjį projektą, kurio tikslas - sukurti tinklą visoje Europoje ir išbandyti pirmąją tokio pobūdžio infrastruktūrą, skirtą koordinuoti ir paskirstyti 98 Europos eukariotų rūšių etaloninių genomų nuskaitymą iš mėginių surinktų 34 Europos šalyse. Čia aprašome procesą ir iššūkius su kuriais susidurta kuriant bandomąją etaloninių genomų išteklių kūrimo infrastruktūrą, bei nagrinėjame šio metodo veiksmingumą siekiant aukštos kokybės etaloninių genomų kūrimo, atsižvelgdami ir į lygiateisiškumą bei įtrauktį. Šio bandomojo projekto rezultatai ir išmoktos pamokos suteikia tvirtą pagrindą ERGA ir kartu suteikia svarbios patirties kitiems tarptautiniams ir nacionaliniams genomų išteklių projektams.

Dutch:

Een wereldwijde genoom database gevuld met de complete diversiteit aan soorten op aarde kan een schatkamer voor wetenschappelijke ontdekkingen vormen. Ondanks de grote vooruitgang in genoom sequencing-technieken is er momenteel slechts genoom data beschikbaar voor een minuscule fractie van alle soorten op aarde. Om bij te dragen aan een meer complete genoom database van deze planeet zijn wetenschappers en instituten uit de hele wereld samengekomen in het Earth BioGenome Project (EBP). Dit project heeft als doel het sequensen en samenvoegen van hoge kwaliteit referentie genomen voor alle ~1,5 miljoen bekende eukaryote soorten in een stapsgewijze aanpak. Het initiatief gaat momenteel over naar fase II, waarbij in slechts 4 jaar tijd de genoomsequentie van 150.000 soorten moet worden bepaald. Hierbij is wereldwijde deelname cruciaal voor succes. De Europese tak van het EBP, de Europese Referentie Genoom Atlas (ERGA), heeft tot doel het implementeren van een nieuw, gedecentraliseerd, toegankelijk, rechtvaardig en inclusief model voor het produceren van hoge kwaliteit referentie genomen, wat zal bijdragen aan de EBP wanneer het opschaalt. Om dit te realiseren heeft ERGA een proefproject gelanceerd. Hierin is een Europees netwerk ingericht voor het ontwikkelen en testen van de eerste infrastructuur van zijn soort voor de gecoördineerde en gedecentraliseerde productie van referentie genomen van 98 Europese eukaryote soorten verzameld in 34 Europese landen. Hier schetsen we het proces en de uitdagingen die we tegenkwamen tijdens het ontwikkelen van deze proef-infrastructuur voor het produceren van referentie genomen en -databases, en onderzoeken we de effectiviteit van deze aanpak aangaande de productie van hoge kwaliteit referentie genomen, waarbij de rechtvaardigheid en inclusie ook zijn meegenomen. De uitkomsten en geleerde lessen tijdens het proefproject vormen een solide onderbouwing voor ERGA en bieden tegelijkertijd een aantal belangrijke lessen die ook van toepassing zijn op andere transnationale en nationale projecten rond het beschikbaar maken van genoom data.

Polish:

Ogólnoświatowa baza danych zawierająca w sobie dane genomowe wszystkich gatunków żyjących na Ziemi, byłaby skarbnicą wiedzy dla przyszłych badaczy. Pomimo dużych postępów w rozwoju technologii sekwencjonowania, dane genomowe są ogólnodostępne tylko dla niewielkiej część gatunków. W celu stworzenia bardziej kompletnej międzynarodowej bazy danych genomicznych, naukowcy i instytucje z całego świata zjednoczyli się w ramach projektu Earth BioGenome Project (EBP), który planuje stopniowe sekwencjonowanie i składanie wysokiej jakości genomów referencyjnych dla wszystkich ok. 1,5 miliona znanych gatunków eukariotycznych. W najbliższym czasie inicjatywa przechodzi do fazy II, w której w ciągu zaledwie czterech lat mają zostać zsekwencjonowane genomy 150 000 gatunków. W związku z tym międzynarodowe zaangażowanie będzie miało fundamentalne znaczenie. Europejski węzeł EBP, o nazwie Europejski Atlas Genomów Referencyjnych (ang. European Reference Genome Atlas; ERGA) ma na celu wdrożenie nowego, zdecentralizowanego, sprawiedliwego i dostępnego dla wszystkich modelu sekwencjonowania i składania wysokiej jakości genomów referencyjnych, a także stopniowe przekazywanie tych informacji do EBP. W celu realizacji przyjętej misji, ERGA uruchomiła projekt pilotażowy (ang. pilot project) zmierzający do utworzenia w całej Europie sieci współpracy. Projekt ten będzie polegał na opracowaniu i przetestowaniu zastosowania rozproszonej infrastruktury do skoordynowanego zsekwencjonowania i składania genomów referencyjnych dla 98 europejskich gatunków eukariotycznych, zebranych przez badaczy z 34 europejskich krajów. W niniejszej pracy przedstawiamy proces i wyzwania przed którymi stoimy podczas rozwijania tego projektu. Analizujemy także jego skuteczność do generowania wysokiej jakości genomów referencyjnych, mając na uwadze także równość szans i inkluzywność. ERGA ma nadzieję, że wyniki i wnioski wyciągniete z realizacji projektu pilotażowego będą cenne nie tylko dla tej inicjatywy, ale także że będą cennymi wskazówkami w realizacji podobnych projektów o zasięgu krajpowym i międzynarodowym w przyszłości.

Portuguese:

Uma base de dados genómica de toda a diversidade de espécies da Terra poderá ser um tesouro de descobertas científicas. No entanto, independentemente dos grandes avanços nas tecnologias de sequenciação de genomas, apenas uma pequena fração das espécies tem informação genómica disponível. Para contribuir para uma base de dados genómica planetária mais completa, cientistas e instituições de todo o mundo uniram-se no Earth BioGenome Project (EBP), que planeia sequenciar e gerar genomas de referência de alta qualidade para todas as cerca de 1,5 milhões de espécies eucarióticas conhecidas, através de uma abordagem gradual e faseada. À medida que a iniciativa transita para a Fase II, na qual 150.000 espécies serão sequenciadas em apenas quatro anos, a participação de cientistas e instituições de todo o mundo será fundamental para o seu sucesso. Como nó Europeu do EBP, o European Reference Genome Atlas (ERGA) procura implementar um novo modelo descentralizado, acessível, equitativo e inclusivo para a produção de genomas de referência de alta qualidade, que informará o EBP enquanto este cresce. Para embarcar nesta missão, o ERGA lançou um Projeto Piloto para estabelecer uma rede através da Europa para desenvolver e testar a primeira infraestrutura deste tipo, para a produção coordenada e distribuída de genomas de referência de 98 espécies eucarióticas europeias, a partir de doadores de amostras de 34 países europeus. Aqui descrevemos o processo e os desafios enfrentados durante o desenvolvimento de uma infraestrutura piloto para a produção de recursos genómicos de referência e exploramos a eficácia desta abordagem em termos de produção de genomas de referência de alta qualidade, considerando também a equidade e a inclusão. Os resultados e lições aprendidas durante este piloto fornecem uma base sólida para o ERGA, e conhecimento importante para a implementação de outros projetos de recursos genómicos transnacionais e nacionais.

Romanian:

O bază de date genomică globală, cu toată diversitatea speciilor de pe Pământ ar putea fi o comoară de descoperiri științifice. Cu toate acestea, în ciuda progreselor majore în tehnologiile de secvențiere genomică, doar o mică parte din specii au informații genomice disponibile. Pentru a contribui la completarea bazei de date genomice planetare, oamenii de știință și instituțiile din întreaga lume s-au unit în cadrul Proiectului Earth BioGenome (EBP), care intenționează să secvențieze și să asambleze genomuri de referință de înaltă calitate pentru toate ~1,5 milioane de specii de eucariote recunoscute printr-o abordare treptată. Pe măsură ce inițiativa trece la Faza II, unde 150.000 de specii urmează să fie secvențiate în doar patru ani, participarea la nivel mondial la acest proiect va fi fundamentală pentru succes. În calitate de nod european al EBP, Atlasul European al Genomurilor de Referință (ERGA) încearcă să implementeze un nou model descentralizat, accesibil, echitabil și incluziv pentru producerea de genomuri de referință de înaltă calitate. Pentru a se angaja în această misiune, ERGA a lansat un proiect pilot pentru a stabili o rețea în întreaga Europă pentru a dezvolta și testa prima infrastructură de acest gen pentru producția coordonată și distribuită de genomuri de referință pe 98 de specii eucariote europene colectate de furnizorii de mostre din 34 de țări europene. Aici descriem procesul și dificultățile cu care s-a confruntat dezvoltarea infrastructurii pilot necesară pentru obținerea de genomuri de referință, și explorăm eficacitatea acestei abordări în ceea ce privește producția de genomuri de referință de înaltă calitate, luând în considerare atât echitatea, cât și incluziunea. Rezultatele și lecțiile învățate în timpul acestui proiect pilot oferă o bază solidă pentru ERGA, oferind în același timp lecții cheie altor proiecte transnaționale și naționale de resurse genomice.

Slovakian:

Globálna databáza genómov všetkých druhov na Zemi by mohla byť mimoriadne významným zdrojom mnohých vedeckých objavov. Napriek veľkému pokroku v technológiách sekvenovania, informácie o celých genómoch sú k dispozícii stále len u veľmi malej časti druhov. S cieľom prispieť ku kompletnejšej databáze genómov sa vedci a inštitúcie z celého sveta spojili v rámci projektu Earth BioGenome Project (EBP), ktorý plánuje sekvenovať a zostaviť vysokokvalitné referenčné genómy pre všetkých ~1,5 milióna známych eukaryotických druhov prostredníctvom postupného, fázového prístupu. Keďže iniciatíva prechádza do Fázy II, v ktorej sa majú sekvenovať genómy 150 000 druhov v priebehu 4 rokov, zásadná bude pre úspech projektu celosvetová účasť. Konzorcium European Reference Genome Atlas (ERGA) ako európsky uzol EBP sa snaží zaviesť nový, decentralizovaný, dostupný, spravodlivý a inkluzívny model produkcie vysokokvalitných referenčných genómov, ktorý poskytne informácie EBP, ako cieľ Fázy II efektívne dosiahnuť. Aby sa konzorcium ERGA mohlo pustiť do tejto misie, spustili sme pilotný projekt na vytvorenie európskej siete s cieľom vyvinúť a otestovať prvú infraštruktúru svojho druhu na koordinovanú a distribuovanú produkciu referenčných genómov 98 európskych druhov od poskytovateľov vzoriek z 34 európskych krajín. V tejto štúdii uvádzame proces a výzvy, ktorým sme čelili počas vývoja pilotnej infraštruktúry a hodnotíme účinnosť tohto prístupu z hľadiska produkcie vysokokvalitných referenčných genómov, pričom zohľadňujeme aj spravodlivosť a inklúziu. Výsledky a skúsenosti získané počas tohto pilotného projektu poskytujú dôležitý základ pre fungovanie konzorcia a zároveň ponúkajú kľúčové poznatky pre iné nadnárodné a národné projekty využívajúce genomické dáta.

Slovenian:

Globalna zbirka podatkov o genomih vseh vrst na Zemlji bi lahko bila zakladnica znanstvenih odkritij. Kljub velikemu napredku na področju tehnologij sekvenciranja genomov pa imamo trenutno na razpolago genomske podatke le za majhno število vrst. Da bi prispevali k popolnejši planetarni podatkovni bazi genomov, so se znanstveniki in institucije po vsem svetu združili v projektu Earth BioGenome Project (EBP), katerega cilj je postopno sekvenciranje in sestavljanje visokokakovostnih referenčnih genomov za približno 1,5 milijona priznanih evkariontskih vrst. Ker pobuda prehaja v drugo fazo, v kateri naj bi v štirih zaporednih letih sekvencirali 150.000 vrst, bo za njen uspeh ključno globalno sodelovanje. Evropski referenčni genomski atlas (ERGA) kot evropsko središče projekta EBP želi vzpostaviti nov decentraliziran, dostopen, pravičen in vključujoč model za zagotavljanje visokokakovostnih referenčnih genomov, ki bo podpiral nadaljnje razširjene pobude projekta EBP. Konzorcij ERGA je začel izvajati pilotni projekt za vzpostavitev vseevropskega omrežja za razvoj in prvo testiranje dostopne infrastrukture za usklajeno in porazdeljeno sekvenciranje referenčnih genomov za 98 evropskih evkariontskih vrst, katerih vzorce so predložili raziskovalci iz 34 evropskih držav. V nadaljevanju opisujemo postopek in izzive, s katerimi smo se soočili med pilotnim preizkusom infrastrukture za referenčne genome, hkrati pa ocenjujemo učinkovitost tega pristopa pri določanju visokokakovostnih referenčnih genomov ob upoštevanju pravičnosti in vključenosti. Rezultati in izkušnje, pridobljene pri tem pilotnem projektu, so trdna podlaga za nadaljnje delo konzorcija ERGA, hkrati pa ponujajo ključne izkušnje za druge mednarodne in nacionalne projekte na področju analize genomov.

Spanish:

Una base de datos global de genomas de toda la diversidad de especies de la Tierra podría ser un tesoro de descubrimientos científicos. Sin embargo, independientemente de los grandes avances en las tecnologías de secuenciación, tan solo una pequeña fracción de las especies tiene información genómica disponible. Para contribuir a una base de datos genómica planetaria más completa, científicos e instituciones de todo el mundo se han unido bajo el Proyecto Earth

BioGenome (EBP), el cual planea secuenciar y ensamblar genomas de referencia de alta calidad para las ~1,5 millones de especies eucariotas reconocidas a través de una aproximación por fases. A medida que esta iniciativa entre en la Fase II, en la que se secuenciarán 150.000 especies en tan solo cuatro años, la participación mundial en el proyecto será fundamental para su éxito. Como nodo europeo de la EBP, el Atlas Europeo de Genomas de Referencia (ERGA) busca implementar un nuevo modelo descentralizado, accesible, equitativo e inclusivo para producir genomas de referencia de alta calidad, el cual informará a la EBP a medida que vaya escalando su producción. Para embarcarse en esta misión, ERGA lanzó un proyecto piloto con la intención de establecer una red en toda Europa con el fin de desarrollar y probar la primera infraestructura de este tipo destinada a la producción coordinada y distribuida de genomas de referencia en 98 especies eucariotas europeas, procedentes de 34 países europeos proveedores. Aquí describimos el proceso y los desafíos a los que nos hemos enfrentado durante el desarrollo de una infraestructura piloto para la producción de recursos genómicos de referencia, y exploramos la efectividad de este enfoque en términos de producción de genomas de referencia de alta calidad, considerando también la equidad y la inclusión. Los resultados y las lecciones aprendidas durante este piloto constituyen una base sólida para ERGA, al tiempo que ofrecen aprendizajes clave para otros proyectos de recursos genómicos transnacionales y nacionales.

Swedish:

En global databas över jordens alla arters hela genom (arvsmassa) skulle utgöra en veritabel skattkista för vetenskapliga upptäckter. Men trots stora teknologiska framsteg inom DNA-sekvensering så har hittills bara en bråkdel av alla arters hela genom sekvenserats. För att bidra till en mer komplett planetär genomdatabas har forskare och institutioner världen över gått samman i Earth Biogenome Project (EBP), ett projekt som med en stegvis strategi planerar att sekvensera och sätta samman högkvalitativa referensgenom för alla ~1.5 miljoner kända arter av Eukaryoter. I nästa fas är målet att sekvensera 150 000 arter på bara fyra år, och för att nå dit krävs storskaligt globalt engagemang. European Reference Genome Atlas (ERGA), Europas nod av EBP, utvecklar nu en ny decentraliserad, öppen, rättvis och inkluderande modell för produktion av högkvalitativa referensgenom, en modell med stor relevans för EBPs målsättning. Som start lanserade ERGA ett pilotprojekt med syfte att etablera ett europeiskt nätverk för att utveckla och testa denna första infrastruktur av sitt slag. Piloten innefattade koordinering av en distribuerad produktion av referensgenom för 98 europeiska arter från 34 olika europeiska länder. Här beskriver vi processen och utmaningarna för utvecklingen av pilot-infrastrukturen och utvärderar effektiviteten vad gäller produktion av högkvalitativa referensgenom, beaktande såväl inklusivitet som rättviseaspekter. Resultaten och lärdomarna från pilotprojektet utgör en stabil grund för ERGA med stor nytta även för andra nationella och internationella genomresursprojekt.

Icelandic:

Almennur gagnagrunnur erfðamengjasem spannar líffræðilegan fjölbreytileika jarðar væri sannkölluð fjársjóðskista vísindalegra upplýsinga. Þrátt fyrir miklar framfarir í raðgreiningum erfðamengja eingöngu erfðamengaraðir lítils brots af öllum tegundum aðgengilegar. Til að fá heilsteyptari gagnagrunn yfir erfðamengi lífvera á jörðinni hafa vísindamenn og stofnanir víðaum heim sameinast í Earth BioGenome Project (EBP), sem stefnir að því að raðgreina og kortleggja hágæða-viðmiðunarerfðamengi fyrir allar þær ~1,5 miljónir tegundir heilkjörnunga sem lýst hefur verið, meið skipulöðum hætti. Þar sem annar áfangi verkefnisins (e.Phase II), þar sem 150.000 tegundir verða raðgreindar á aðeins fjórum árum, er nú að hefjast er almenn og alþjóðleg þáttaka mikilvæg til að árangur náist . Evrópuhluta EBP verkefnisins, evrópska viðmiðunarerfðamengja-atlasinum (e. the European Reference Genome Atlas (ERGA)), er ætlað að útfæra nýja aðferð til að setja saman hágæða-viðmiðunarerfðamengi með dreifðri þátttöku og almennu aðgengi þar sem jafnræði er tryggt, sem mun miðla upplýsingum til EBP jafnóðum. Til að framfylgja þessu hefur ERGA sett af stað forverkefni (e. a Pilot Project) sem byggir á samstarfsneti sem spannar alla Evrópu, til að þróa og prófa slíka innviði fyrir greiningu og miðlun viðmiðunarerfðamengja fyrir 98 tegundir evrópskra heilkjörnunga sem safnað var af þátttakendum frá 34 Evrópulöndum. Hér greinum við frá aðferðafræðinni og áskorunum sem við stóðum frammi fyrir við að þróa þessa innviði og hvernig við fundum leiðir til að safna þessum viðmiðunarerfðamengjum. Auk þess metum við hversu vel það gekk m.t.t. jafnræðis og virkrar þátttöku allra þátttakenda. Þær niðurstöður og sá lærdómur sem við höfum aflað í þessu forverkefni leggur góðan grunn að ERGA auk þess að miðla grunnþekkingu til annarra alþjóðlegra og landsbundinna erfðamengjaverkefna.

Norwegian:

En global genomdatabase over hele jordens artsmangfold kunne vært en skattkiste for vitenskapelige oppdagelser. Men til tross for store fremskritt innen genomsekvenseringsteknologi er det kun en svært liten del av artene som har genomisk informasjon tilgjengelig. For å bidra til en mer komplett verdensomspennende genomdatabase har forskere og institusjoner over hele verden gått sammen i Earth BioGenome prosjektet (EBP), som planlegger å sekvensere og assemblere referansegenomer av høyeste kvalitet for alle ~1,5 millioner anerkjente eukaryote artene, gjennom en trinnvis tilnærming. Initiativet går nå over i den andre fasen, hvor 150 000 arter skal sekvenseres i løpet av kun fire år, og verdensomfattende deltakelse i prosjektet er derfor avgjørende for å lykkes. European Reference Genome Atlas (ERGA), som er den europeiske grenen i EBP, har som mål å implementere en ny desentralisert, tilgjengelig, rettferdig og inkluderende modell for produksjonen av referansegenomer av høyeste kvalitet, som vil informere EBP samtidig som den utarbeides. For å ta fatt på dette oppdraget lanserte ERGA et pilotprosjekt for å etablere et nettverk over hele Europa som skal utvikle og teste den første infrastrukturen i sitt slag for koordinert og distribuert referansegenomproduksjon for 98 europeiske eukaryote arter. Disse prøvene kommer fra samarbeid med partnere i 34 europeiske land. Her skisserer vi prosessen og utfordringene under utviklingen av en pilotinfrastruktur for produksjonen av referansegenomressurser, og undersøker hvor effektiv denne tilnærmingen er når det gjelder produksjon av referansegenomer av høyeste kvalitet, samtidig som vi tar hensyn til rettferdighet og inkludering. Resultatene og erfaringene fra dette pilotprosjektet danner et solid grunnlag for ERGA, samtidig som det gir viktige erfaringer til andre transnasjonale og nasjonale genomiske ressursprosjekter.

Faroese:

Ein heimsfevnandi genomdátugrunnur við øllum lívveru sløgum á jørðini, kundi verið ein dýrgripur av vísindaligum gjøgnumbrotum. Men tíverri er hetta ikki veruleiki enn. Hóast stóra framgongd innan genom tøkni, so er tað bert ein lítil brotpartur av øllum lívveru sløgum, ið eru genom kanna. Fyri at fáa gongd á ein slíkan genomdátugrunn hava vísindafólk og stovnar runt allan heimin skipa seg undir heitinum Earth BioGenome Project (EBP). Hetta er ein verkætlan ið hevur til endamáls at framleiða hágóðsku tilvísingargenom, fyri tey áleið 1.5 milliónir kendu lívveru sløgini. Verkætlanin nærkast nú øðrum stigi, har tilvísingargenom fyri 150,000 lívveru sløg skulu framleiðast uppá fýra ár. Fyri at klára hesa stóru uppgávu, er neyðugt at allur heimurin tekur lut í hesi verkætlan. European Reference Genome Atlas (ERGA), ið er tann europeiski parturin av verkætlanin, er í holt við at gera eina forskrift fyri hvussu vit framleiða hágóðsku tilvísingargenom. Hesin frymil tekur hædd fyri miðspjaðan, atkomuligheit, rættvísi og inklusjón. Til hesa uppgávu, hevur ERGA skapa eitt europeiskt netverk, til at menna og royna eitt undirstøðukervi, ið samskipar framleiðsluna av tilvísingargenomum fyri 98 europeisk lívveru sløg frá 34 europeiskum londum. Í hesi grein útgreina vit mannagongdir og avbjóðingar, ið tóku seg upp tá tilvísingargenom undirstøðukervið var ment, og kanna hvussu effektivt hetta hevur verið í mun til framleiðslu av hágóðsku tilvísingargenomum, við rættvísi og inklusjón í huganum. Úrtøkurnar og vitanin ið er funnin í hesum fyrsta partinum av verkætlanini, er góður grundsteinur til víðari menning av ERGA, og gevur týdningarmiklar leiðreglur til aðrar líknandi verkætlanir.

Hebrew:

למאגר מידע גנומי גלובלי של מגוון המינים על פני כדור הארץ יש פוטנציאל להוות אוצר של תגליות מדעיות. עם זאת, למרות ההתקדמות המשמעותית בטכנולוגיות הריצוף הגנומי, קיים מידע גנומי זמין רק לחלק זעיר ממגוון המינים בעולם. על מנת לתרום ליצירת מאגר מידע גנומי מלא של כל כדור הארץ, מדענים ומרכזי מחקר מכל קצוות תבל חברו תחת פרויקט הביוגנום העולמי - EBP - אשר שם לו כמטרה ריצוף והרכבה של גנום יחוס באיכות גבוהה של כל אחד ממיליון וחצי המינים האאוקריוטים המוכרים. מטרה זו תושג באמצעות גישה רב-שלבית מדורגת ומתואמת. עם המעבר לשלב השני של היוזמה, שבמהלכו ירוצף הגנום של 150,000 מינים על פני ארבע שנים בלבד, החשיבות של מעורבות עולמית בפרויקט הופכת להיות מרכזית. אטלס הייחוס הגנומי האירופי (ERGA) מתוכנן להיות הצומת האירופאי של EBP. הוא שואף ליישם גישה מבוזרת, נגישה, שוויונית ומכילה לייצור ריצופי ייחוס גנומיים באיכות גבוהה, שניתן יהיה להגדילה עם הזמן ותוך תיאום עם EBP. כנקודת זינוק למשימה זו, EGRA השיק פרויקט פיילוט שמטרתו לבסס רשת כלל ארופאית שתפתח ותבדוק את התשתית הראשונה מסוגה להפקת רצפי גנום יחוס באופן מתואם ומבוזר. במסגרת הפיילוט הופקו גנומים מלאים עבור 98 מינים אאוקריוטים מאירופה, על בסיס דגימות שהגיעו מ-34 מדינות אירופאיות. אנו מציגים כאן בראשי פרקים גם את התהליך אותו עבר הפרויקט וגם את האתגרים עימם התמודד במהלך פיתוח התשתיות לייצור משאבים גנומיים. אנו בוחנים את יעילות הגישה בה נקט ERGA לייצור רצפי ייחוס גנומיים ברמה הגבוהה תוך שאנו גם לוקחים בחשבון הוגנות והכללה. התוצרים שהופקו והלקחים שנלמדו במהלך פרויקט הפיילוט מספקים תשתית יציבה להמשך עבור ERGA, ויכולים גם לשמש כבסיס ידע לפרויקטים גנומיים לאומיים ובינלאומיים אחרים.

Serbian:

Globalna baza podataka genoma svih vrsta na Zemlji mogla bi predstavljati veoma značajno naučno otkriće. Međutim, bez obzira na veliki napredak u tehnologijama sekvenciranja genoma, samo mali deo vrsta ima dostupne genomske informacije. Da bi doprineli potpunijoj planetarnoj genomskoj bazi podataka, naučnici i institucije širom sveta su se ujedinili u okviru projekta Earth BioGenome Project (EBP), u okviru kog se planira sekvenciranje i sakupljanje visokokvalitetnih referentnih genoma za svih ~1,5 miliona poznatih eukariotskih vrsta kroz višefazni proces. Kako inicijativa prelazi u fazu II, gde će 150.000 vrsta biti sekvencionirano za samo četiri godine, široko učešće u projektu biće od suštinskog značaja za uspeh. Kao evropski čvor EBP-a, Evropski referentni atlas genoma (ERGA) nastoji da implementira novi decentralizovan, pristupačan, pravičan i inkluzivan model za generisanje visokokvalitetnih referentnih genoma, koji će doprineti EBP-u. Da bi se upustila u ovu misiju, ERGA je pokrenula Pilot projekat za uspostavljanje mreže širom Evrope za razvoj i testiranje prve infrastrukture te vrste za koordinisano generisanje i distribuciju referentnih genoma 98 evropskih eukariotskih vrsta obezbeđenih iz 34 evropske zemlje. Ovde prikazujemo proces i izazove sa kojima se suočavamo tokom razvoja pilot infrastrukture za generisanje referentnih genoma, i istražujemo efikasnost ovog pristupa u smislu određivanje referentnih genoma visokog kvaliteta, uzimajući u obzir i jednakost i inkluziju. Ishodi i lekcije naučene tokom ovog pilot-projekta pružaju solidnu osnovu za ERGA dok nude ključna znanja drugim transnacionalnim i nacionalnim projektima genoma.

Ukrainian:

Глобальна база даних геномів усього різноманіття видів Землі може стати скарбницею наукових відкриттів. Однак, незважаючи на значні досягнення в технологіях секвенування геному, лише незначна частка видів має наявну геномну інформацію. Щоб зробити свій внесок у створення більш повної планетарної геномної бази даних, вчені та інституції з усього світу об'єдналися в рамках проекту *Earth BioGenome Project (EBP)*, який планує секвенувати та зібрати високоякісні референсні геноми для всіх ~1,5 мільйонів визнаних еукаріотичних видів шляхом підходу поетапного дослідження. Оскільки ця ініціатива переходить у другу фазу, де 150 000 видів мають бути секвеновані всього за чотири роки, залучення учасників з усього світу до проекту буде фундаментальним для його успіху. Як європейський вузол *EBP, European Reference Genome Atlas (ERGA)* прагне запровадити нову децентралізовану, доступну, справедливу та інклюзивну модель для створення високоякісних референсних геномів, яка інформуватиме *EBP* у міру його масштабування. Щоб розпочати цю місію, *ERGA* запустила пілотний проект для створення мережі по всій Європі для розробки та тестування першої інфраструктури такого роду для скоординованого та розподіленого створення референсних геномів 98 європейських еукаріотичних видів від постачальників зразків з 34 європейських країн. Тут ми окреслюємо процес та виклики, з якими ми зіткнулися під час розробки пілотної інфраструктури для створення референсних геномних ресурсів, і досліджуємо ефективність цього підходу з точки зору створення високоякісних референсних геномів, враховуючи також справедливість та інклюзивність. Результати та уроки, отримані під час цього пілотного проекту, створюють фундаментальні підвалини для *ERGA*, водночас пропонуючи ключові знання для інших транснаціональних та національних проектів з геномних ресурсів.

Catalan:

Una base de dades global que contingui el genoma de la diversitat de totes les espècies del planeta Terra podria ser un tresor de descobriments científics. Tanmateix, malgrat els grans avenços en la tecnologia de seqüenciació del genoma, només es disposa informació genòmica d’una petita fracció del conjunt de totes les espècies. Per contribuir a una base de dades genòmica planetària més completa, científics i institucions de tot el món s'han unit sota el *Earth BioGenome Project* (EBP). Aquest projecte, mitjançant un enfoc gradual dividit en diferents fases, té com objectiu seqüenciar i fer l’assemblatge de genomes de referència d'alta qualitat per aproximadament l’1,5 milions de totes les espècies eucariotes conegudes. A mesura que la iniciativa passa a la Fase II, on es té previst seqüenciar 150.000 espècies en només quatre anys, la participació en el projecte de tots els actors a nivell mundial serà fonamental per assolir els objectius. Com a node europeu de l'EBP, l'Atles de Genomes de Referència Europeu (*European Reference Genome Atlas*, ERGA) busca implementar un nou model descentralitzat, accessible, equitatiu i inclusiu per produir genomes de referència d'alta qualitat, que a mesura que s’avanci, anirà informant l'EBP. En iniciar-se aquesta missió, l’ERGA va llançar un Projecte Pilot que va establir una xarxa europea per desenvolupar i provar una primera infraestructura d'aquest tipus.

En un primer moment es va realitzar la producció coordinada i la distribució dels genomes de referència de 98 espècies eucariotes europees, amb proveïdors de mostres de 34 països europeus. A continuació es descriuen els processos i els reptes que s'han hagut d'afrontar durant el desenvolupament de la infraestructura del Projecte Pilot per a la producció de recursos de genomes de referència, i s'explora l'eficàcia d'aquest enfocament en la producció de genomes de referència d'alta qualitat, considerant també els principis d'equitat i inclusió. Els resultats i les lliçons apreses durant aquest Projecte Pilot proporcionen una base sòlida per a l'ERGA, alhora que ofereixen aprenentatges clau per altres projectes de recursos genòmics transnacionals i nacionals.

Croatian:

Globalna baza podataka s genomima svih vrsta na Zemlji biti će riznica znanstvenih otkrića. Međutim, bez obzira na veliki napredak u tehnologiji sekvenciranja genoma, samo mali dio vrsta ima dostupne genomske podatke. Kako bi pridonijeli u stvaranju što potpunije globale baze genoma, znanstvenici i institucije diljem svijeta ujedinili su se u okviru inicijative Earth BioGenome Project (EBP), kojoj je cilj postepeno sekvencirati i sastaviti visokokvalitetne referentne genome za ~1,5 milijun poznatih eukariotskih vrsta. Kako inicijativa ulazi u drugu fazu, kroz koju će se u samo četiri godine sekvencirati genomi 150 000 vrsta, širenje sudjelovanja u projektu na cijeli svijet biti će ključno za njegov uspjeh. Kao europski predstavnik EBP-a, inicijativa Europski atlas referentnih genoma (European Reference Genome Atlas, ERGA) nastoji implementirati novi decentralizirani, pristupačan, pravičan i uključiv model za proizvodnju visokokvalitetnih referentnih genoma. Kao prvi korak ove misije, ERGA je pokrenula pilot projekt za uspostavljanje europske mreže s ciljem razvoja i testiranja infrastrukture, prve te vrste, za koordinirano i distribuirano sekvenciranje referentnih genoma na 98 europskih eukariotskih vrsta odabranih od predstavnika iz 34 europske zemlje. Ovdje opisujemo proces i izazove s kojima smo se suočili u pilot projektu tijekom razvoja infrastrukture potrebne za produkciju referentnih genoma, i istražujemo učinkovitost ovog pristupa za proizvodnju visokokvalitetnih referentnih genoma, uzimajući u obzir jednakost i uključenost. Ishodi i lekcije naučene tijekom ovog pilot-projekta daju čvrstu osnovu za ERGA-u, a istovremeno nude ključna znanja drugim internacionalnim i nacionalnim genomskim projektima.

Czech:

Globální databáze genomů všech druhů organismů, žijících na zemi by byla neocenitelnou pomůckou na cestě ke vědeckým objevům. Bohužel, i přes významné pokroky v technologiích sekvenace genomů jsou genomické informace dostupné pouze pro zanedbatelné množství druhů. Proto, aby bylo možné vytvořit úplnější globální databázi genomů, se spojili vědci a výzkumné instituce v rámci projektu Earth BioGenome Project (EBP), jehož cílem je postupně sekvenovat a zkompletovat vysoce kvalitní referenční genomy pro všech ~ 1.5 milionu známých druhů eukaryotních organismů. Protože tato iniciativa nyní vstupuje do druhé fáze, během níž by mělo být v průběhu čtyř let sekvenováno 150,000 druhů, je pro její úspěch nezbytná participace vědců z celého světa. European Reference Genome Atlas (ERGA), jako evropský uzel EBP, usiluje o vytvoření decentralizovaného, dostupného, spravedlivého a inkluzivního postupu tvorby kvalitních referenčních genomů, který by k tomuto úsilí přispěl. Jako počátek této mise spustila ERGA pilotní projekt, který měl za cíl vytvořit celoevropskou síť pro vývoj a testování infrastruktury pro koordinovanou a distribuovanou produkci genomů 98 evropských druhů eukaryot, pocházejících ze vzorků, dodaných spolupracovníky ze 34 evropských států. Zde popisujeme postupy a výzvy, se kterými se tato iniciativa setkala při tvorbě pilotní infrastruktury pro produkci referenčních genomů a posuzujeme efektivitu tohoto přístupu, přičemž bereme v úvahu rovněž spravedlnost a inkluzi. Praktické výstupy i zkušenosti, získané v průběhu této pilotní studie tvoří solidní základ pro další činnost ERGA a nabízí rovněž získané zkušenosti dalším genomickým projektům národních i nadnárodních úrovní.

Estonian:

Ülemaailmne, kogu maakera liigilist mitmekesisust hõlmav genoomne andmebaas võib kujuneda teaduslike avastuste aardelaekaks. Hoolimata genoomi sekveneerimis- ehk järjestamistehnoloogia suurtest edusammudest on genoomne teave olemas vaid väga väikesel osal liikidest. Selleks, et kaasa aidata üha terviklikuma, kogu maakera genoomiandmebaasi loomisele, on teadlased ja institutsioonid üle kogu maailma koondunud Maa biogenenoomimiprojekti (Earth BioGenome Project, EBP) alla, mille raames on kavas järk-järgult järjestada kõigi ~1,5 miljoni teadaoleva eukarüoodi ehk päristuumse liigi kvaliteetsed referentsgenoomid. Kuna algatus on liikumas üle II etappi, milles nelja aasta jooksul järjestatakse 150 000 liigi genoomid, on ülemaailmne osalus projekti õnnestumiseks väga oluline. EBP Euroopa sõlmena püüab Euroopa referentsgenoomide atlas (ERGA) rakendada uut, detsentraliseeritud, juurdepääsetavat, õiglast ja kaasavat mudelit kvaliteetsete referentsgenoomide loomiseks. Selle missiooni alustamiseks käivitas ERGA pilootprojekti, mille raames loodi üle-Euroopaline võrgustik, et arendada ja katsetada esimest omataolist taristut ning nii järjestati koordineeritud viisil 34st Euroopa riigist pärit proovimaterjali põhjal 98 eukarüootse liigi referentsgenoomid. Siinkohal kirjeldame referentsgenoomi ressursside loomise piloottaristu väljatöötamist ning sellega kaasnenud probleeme ning uurime selle lähenemisviisi tõhusust kvaliteetsete referentsgenoomide loomise seisukohast, võttes sealjuures arvesse ka võrdsust ja kaasatust. Selle pilootprojekti tulemused ja õppetunnid on ERGA jaoks heaks vundamendiks, pakkudes samal ajal olulisi õppetunde teistele riikidevahelistele ja -sisestele genoomiprojektidele.

Finnish:

Maailmanlaajuinen genomitietokanta koko planeetan lajien monimuotoisuudesta voisi olla tieteellisten löytöjen aarreaitta. Siitä huolimatta, että genomin sekvensointitekniikoissa on otettu suuria edistysaskelia, genomitietoa on saatavilla vain pienestä osasta lajeja. Täydentääkseen genomitietokantaa tutkijat ja instituutiot ympäri maailman ovat liittyneet Earth BioGenome Project (EBP) -projektiin, jonka tavoitteena on vaiheittain sekvensoida ja koota korkealaatuisia referenssigenomeja jokaiselle noin 1,5 miljoonasta tunnetusta eukaryootti lajista. Aloitteen siirtyessä vaiheeseen II, jossa 150 000 lajia on määrä sekvensoida neljässä vuodessa, maailmanlaajuinen osallistuminen hankkeeseen on edellytys sen menestykselle. EBP:n eurooppalaisena osana, European Reference Genome Atlas (ERGA) pyrkii luomaan uuden hajautetun, esteettömän, tasapuolisen ja osallistavan mallin korkealaatuisten referenssigenomien tuottamiseksi, joka pitää EBP:n ajan tasalla prosessin etenemisestä. Ryhtyessään tähän tehtävään, ERGA käynnisti pilottihankkeen luodakseen Euroopan poikki kulkevan verkoston, jonka tarkoituksena on kehittää sekä testata uudenlaista infrastruktuuria, jota käytetään koordinoituun ja hajautettuun referenssi genomien tuotantoon 98:n eurooppalaisen lajin kohdalla, joiden näytteet tulevat 34:stä eri Euroopan maasta. Tässä hahmottelemme prosessia ja haasteita, joita kohtasimme kehittäessämme pilotti-infrastruktuuria referenssigenomien tuotantoa varten, ja tutkimme tämän lähestymistavan tehokkuutta korkealaatuisen referenssigenomi tuotannon kannalta, ottaen huomioon myös tasa-arvon ja inklusiivisuuden. Tämän pilotin aikana saadut tulokset ja opetukset antavat vankan perustan ERGA:lle ja tarjoavat samalla keskeisiä oppitunteja muille monikansallisille ja kansallisille genomi resursseja koskeville projekteille.

Bulgarian:

Световната геномна база данни за видовото разнообразие на Земята може да бъде съкровищница за научни открития. Въпреки това, независимо от големия напредък в технологиите за секвениране на генома, само за малка част от видовете има налична геномна информация. За да допринесат за създаване на по-пълна планетарна геномна база данни, учени и институции от целия свят се обединиха в рамките на Earth BioGenome Project (EBP), който планира да секвенира и сглоби висококачествени референтни геноми за всички ~ 1,5 милиона установени еукариотни вида, чрез стъпаловиден поетапен подход.

Тъй като инициативата преминава във фаза II, където 150 000 вида трябва да бъдат секвенирани само за четири години, световното участие в проекта ще бъде определящо за успеха. Като европейска възлова точка на EBP, Европейският референтен геномен атлас (ERGA) се стреми да приложи нов децентрализиран, достъпен, справедлив и приобщаващ модел за получаване на висококачествени референтни геноми, информирайки  EBP. За да започне тази мисия, ERGA стартира пилотен проект за създаване на мрежа в цяла Европа за разработване и тестване на първата по рода си инфраструктура за координирано и разпределено получаване на референтни геноми на 98 европейски еукариотни вида от 34 европейски държави. Тук очертаваме процеса и предизвикателствата, с които се сблъскваме по време на разработването на пилотната инфраструктура за получаване на референтни геномни ресурси, и изследваме ефективността на този подход по отношение на висококачественото референтно геномно производство, като се има предвид също справедливостта и обхвата. Резултатите и уроците, научени по време на този пилотен проект, осигуряват солидна основа за ERGA, като същевременно предлагат ключови знания за други транснационални и национални проекти за геномни ресурси.

### Hungarian

A Föld teljes faji sokféleségének globális genom-adatbázisa a tudományos felfedezések kincsesbányája lehetne. A genomszekvenálási technológiák jelentős fejlődésétől függetlenül azonban a fajoknak csak egy kis töredéke rendelkezik genomi információkkal. A teljesebb bolygószintű genomikai adatbázis létrehozásához való hozzájárulás érdekében a tudósok és intézmények világszerte összefogtak az Earth BioGenome Project (EBP) keretében, amely fokozatos megközelítéssel tervezi a ~1,5 millió elismert eukarióta faj jó minőségű referencia genomjának szekvenálását és összerakását. Mivel a kezdeményezés a II. fázisba lép, ahol mindössze négy év alatt 150 000 faj szekvenálását tervezik elvégezni, a projektben való világméretű részvétel alapvető fontosságú a sikerhez. Az EBP európai csomópontjaként az Európai Referencia Genom Atlasz (ERGA) egy új, decentralizált, hozzáférhető, méltányos és inkluzív modellt kíván megvalósítani a kiváló minőségű referencia genomok előállítására, amely az EBP növekedéséhez hozzájárulandó, az EBP-t is tájékoztatni fogja. E küldetés megvalósítása érdekében az ERGA kísérleti projektet indított egy európai hálózat létrehozására, amelynek célja a maga nemében első olyan infrastruktúra kifejlesztése és tesztelése, amely 98 európai eukarióta faj koordinált és elosztott referencia-genomjának előállítását teszi lehetővé 34 európai ország mintaadóitól. A következőkben felvázoljuk a referencia-genomforrások előállítására szolgáló kísérleti infrastruktúra fejlesztése során felmerült folyamatokat és kihívásokat, és megvizsgáljuk e megközelítés hatékonyságát a magas színvonalú referencia-genom összerakás szempontjából, figyelembe véve az egyenlőséget és az integrációt is. A kísérleti projekt során elért eredmények és tanulságok szilárd alapot biztosítanak az ERGA számára, miközben kulcsfontosságú ismereteket kínálnak más transznacionális és nemzeti genomikai erőforrás-projektek számára.
