## Supplementary_Table_6 for "The European Reference Genome Atlas: piloting a decentralised approach to equitable biodiversity genomics"

| Scientific name | ToLID | Taxa_1 | Taxa_2 | Status | Type | LongReads_tec | HiC_tec | Asm_Protocol | Mapped | PCR_Dup | NoDup | NoDup_Trans |
|---|---|---|---|---|---|---|---|---|---|---|---|---|
| *Andrena humilis* | iyAndHumi1 | Insecta | Hymenoptera | Non-final | pseudo_hap | HiFi | Proximo | hifiasm_yahs | 59.44% | 57.51% | 1.93% | 17.18% |
| *Osmia cornuta* | iyOsmCorn1 | Insecta | Hymenoptera | Non-final | pseudo_hap | HiFi | Proximo | hifiasm_salsa | 16.63% | 16.43% | 0.20% | 28.85% |
| *Carabus granulatus* | icCarGran1 | Insecta | Coleoptera | Non-final | pseudo_hap | HiFi | Proximo | hifiasm_yahs | 47.14% | 43.10% | 4.04% | 40.94% |
| *Carabus intricatus* | icCarIntr1 | Insecta | Coleoptera | Non-final | pseudo_hap | HiFi | Proximo | hifiasm_yahs | 43.00% | 38.74% | 4.26% | 16.97% |
| *Epeorus assimilis* | ieEpeAssi1 | Insecta | Ephemeroptera | Non-final | pseudo_hap | HiFi | Proximo | hifiasm_yahs | 69.03% | 67.46% | 1.57% | 22.47% |
| *Leptodirus hochenwarti* | icLepHoch2 | Insecta | Coleoptera | Curated | pseudo_hap | HiFi | Arima | hifiasm_yahs | 20.73% | 9.90% | 10.83% | 18.94% |
| *Stylops ater* | ivStyAter1 | Insecta | Strepsiptera | Non-final | pseudo_hap | HiFi | None | hifiasm | NA | NA | NA | NA |
| *Coenonympha glycerion* | ilCoeGlyc1 | Insecta | Lepidoptera | Pre-curation | pseudo_hap | ONT | OmniC | nextdenovo_yahs | 39.81% | 24.65% | 15.15% | 22.28% |
| *Helleia helle* | ilHelHell1 | Insecta | Lepidoptera | Pre-curation | pseudo_hap | ONT | OmniC | nextdenovo_yahs | 47.31% | 45.25% | 2.06% | 15.17% |
| *Parnassius mnemosyne* | ilParMnem1 | Insecta | Lepidoptera | Non-final | pseudo_hap | HiFi | OmniC | hifiasm | 36.88% | 2.96% | 33.92% | 45.64% |
| *Spongipellis delectans* | gfSpoDele1 | Fungi | Agaricomycetes | Curated | pseudo_hap | HiFi | None | hifiasm | NA | NA | NA | NA |
| *Botryllus schlosseri* | kaBotSchl2 | Ascidiacea | Ascidiacea | Non-final | pseudo_hap | ONT | Arima | nextdenovo_yahs | 25.66% | 12.49% | 13.17% | 53.44% |
| *Gordionus montsenyensis* | tfGorSpeb1 | Nematomorpha | Nematomorpha | Pre-curation | pseudo_hap | ONT | Arima | flye_instagraal | 52.04% | 7.92% | 44.13% | 11.03% |
| *Phakellia ventilabrum* | odPhaVent1 | Porifera | Demospongiae | Pre-curation | pseudo_hap | ONT | Arima | nextdenovo_yahs | NA | NA | NA | NA |
| *Phaeosaccion multiseriatum* | uoPhaMult1 | Phaeophyceae | Phaeophyceae | Curated | pseudo_hap | ONT | OmniC | necat_3Ddna | 76.74% | 25.45% | 51.29% | 43.69% |
| *Alburnus alburnus* | fAlbAlb2 | Vertebrata | Actinopterygii | Curated | HiC_phased | HiFi | Arima | hifiasm_salsa | 57.60% | 22.73% | 34.86% | 12.57% |
| *Ammodytes marinus* | fAmmMar1 | Vertebrata | Actinopterygii | Curated | pseudo_hap | HiFi | Arima | hifiasm_yahs | 59.75% | 11.65% | 48.09% | 39.01% |
| *Anaecypris hispanica* | fAnaHis1 | Vertebrata | Actinopterygii | Curated | HiC_phased | HiFi | Arima | hifiasm_salsa | 59.89% | 12.08% | 47.81% | 27.49% |
| *Argentina silus* | fArgSil1 | Vertebrata | Actinopterygii | Pre-curation | pseudo_hap | HiFi | Arima | hifiasm_yahs | 61.93% | 12.10% | 49.83% | 27.70% |
| *Knipowitschia panizzae* | fKniPan1 | Vertebrata | Actinopterygii | Pre-curation | pseudo_hap | ONT | OmniC | flye_yahs | 48.12% | 2.84% | 45.27% | 45.34% |
| *Perca* sp. 'yellow fin Alpine' | fPerYfa1 | Vertebrata | Actinopterygii | Pre-curation | pseudo_hap | HiFi | Arima | hifiasm_yahs | 59.89% | 2.80% | 57.09% | 38.95% |
| *Salvelinus alpinus* | fSalAlp1 | Vertebrata | Actinopterygii | Pre-curation | HiC_phased | HiFi | OmniC | hifiasm_yahs | 34.64% | 6.85% | 27.79% | 48.73% |
| *Silurus aristotelis* | fSilAri1 | Vertebrata | Actinopterygii | Curated | pseudo_hap | HiFi | Arima | hifiasm_yahs | 70.86% | 25.30% | 45.56% | 29.33% |
| *Solea solea* | fSolSol8 | Vertebrata | Actinopterygii | Pre-curation | pseudo_hap | HiFi | Arima | hifiasm_yahs | 76.69% | 26.67% | 50.02% | 11.26% |
| *Tripterygion tripteronotum* | fTriTrp1 | Vertebrata | Actinopterygii | Curated | pseudo_hap | ONT | OmniC | flye_yahs | 70.89% | 5.07% | 65.82% | 24.03% |
| *Zingel asper* | fZinAsp1 | Vertebrata | Actinopterygii | Pre-curation | pseudo_hap | HiFi | Arima | hifiasm_yahs | 62.12% | 4.89% | 57.23% | 38.36% |
| *Haliaeetus albicilla* | bHalAlb1 | Vertebrata | Aves | Curated | pseudo_hap | HiFi | Arima | hifiasm_yahs | 81.32% | 14.11% | 67.21% | 35.89% |
| *Oenanthe leucura* | bOenLec1 | Vertebrata | Aves | Pre-curation | pseudo_hap | HiFi | OmniC | hifiasm_yahs | 73.61% | 13.81% | 59.80% | 22.08% |
| *Tetrao urogallus* | bTetUro2 | Vertebrata | Aves | Pre-curation | pseudo_hap | HiFi | Arima | hifiasm_yahs | 86.62% | 24.61% | 62.01% | 18.10% |
| *Canis aureus* | mCanAur2 | Vertebrata | Mammalia | Pre-curation | HiC_phased | HiFi | Arima | hifiasm_yahs | 74.77% | 5.50% | 69.27% | 36.09% |
| *Chionomys nivalis* | mChiNiv1 | Vertebrata | Mammalia | Curated | pseudo_hap | HiFi | Arima | hifiasm_yahs | 78.50% | 22.69% | 55.81% | 20.77% |
| *Lepus granatensis* | mLepGra1 | Vertebrata | Mammalia | Pre-curation | HiC_phased | HiFi | OmniC | hifiasm_yahs | 73.80% | 20.52% | 53.28% | 20.19% |
| *Lepus timidus* | mLepTim2 | Vertebrata | Mammalia | Pre-curation | pseudo_hap | HiFi | Arima | hifiasm_yahs | 81.37% | 34.18% | 47.18% | 17.23% |
| *Mustela lutreola* | mMusLut1 | Vertebrata | Mammalia | Pre-curation | pseudo_hap | ONT | OmniC | flye_3Ddna | 78.67% | 37.33% | 41.33% | 24.03% |
| *Vipera ursinii* | rVipUrs1 | Vertebrata | Reptilia | Curated | pseudo_hap | HiFi | Arima | hifiasm_yahs | 62.47% | 11.31% | 51.17% | 33.34% |
| *Hottonia palustris* | ddHotPalu1 | Viridiplantae | Ericales | Pre-curation | HiC_phased | HiFi | OmniC | hifiasm_yahs | 48.41% | 4.55% | 43.86% | 40.11% |
| *Prunus brigantina* | drPruBrig1 | Viridiplantae | Rosales | Curated | pseudo_hap | ONT | OmniC | necat_3Ddna | 48.97% | 17.65% | 31.32% | 20.28% |
| *Trifolium dubium* | drTriDubi1 | Viridiplantae | Fabales | Curated | pseudo_hap | HiFi | Arima | hifiasm_yahs | 44.29% | 14.64% | 29.65% | 27.96% |

| Scientific name | NoDup_Cis1kb | NoDup_Cis10kb | gs2_kcov | Asm_Size | GC | Scaff | Cont | Gaps | Gaps_perGbp | Gaps_sum | Gaps_perc | Gaps_auN | Longest_Scaff | Scaff_auN | Scaff_N50 | Scaff_L50 | Scaff_L95 |
|---|---|---|---|---|---|---|---|---|---|---|---|---|---|---|---|---|---|
| *Andrena humilis* | 72.81% | 62.03% | 63.8 | 285,328,606 | 42.62 | 1,067 | 2,468 | 1,401 | 4,910 | 280,200 | 0.098 | 200.0 | 67,981,348 | 32,435,998.6 | 29,133,403 | 4 | 389 |
| *Osmia cornuta* | 51.45% | 40.02% | 7.2 | 436,776,399 | 35.11 | 1,171 | 1,197 | 26 | 60 | 5,200 | 0.001 | 200.0 | 15,011,453 | 4,382,519.9 | 1,362,549 | 49 | 597 |
| *Carabus granulatus* | 48.05% | 39.38% | 163.7 | 166,604,269 | 35.07 | 1,645 | 3,048 | 1,403 | 8,421 | 280,600 | 0.168 | 200.0 | 10,403,647 | 3,929,391.5 | 4,121,613 | 14 | 984 |
| *Carabus intricatus* | 62.03% | 47.10% | 133.9 | 222,370,733 | 34.80 | 356 | 1,127 | 771 | 3,467 | 154,200 | 0.069 | 200.0 | 14,414,676 | 6,740,113.3 | 7,122,293 | 12 | 87 |
| *Epeorus assimilis* | 59.01% | 49.64% | 16.8 | 414,854,051 | 27.01 | 816 | 1,957 | 1,141 | 2,750 | 228,200 | 0.055 | 200.0 | 47,203,024 | 28,202,247.6 | 37,434,202 | 5 | 179 |
| *Leptodirus hochenwarti* | 67.41% | 62.02% | 23.7 | 492,378,077 | 31.80 | 75 | 409 | 334 | 678 | 66,800 | 0.014 | 200.0 | 51,371,384 | 37,025,669.3 | 37,274,531 | 6 | 13 |
| *Stylops ater* | NA | NA | 58.0 | 51,310,711 | 32.64 | 72 | 72 | 0 | 0 | 0 | 0.000 | 0.0 | 5,437,292 | 2,346,880.5 | 1,867,595 | 8 | 37 |
| *Coenonympha glycerion* | 54.47% | 49.01% | 40.2 | 458,639,573 | 37.58 | 65 | 116 | 51 | 111 | 5,100 | 0.001 | 100.0 | 20,120,474 | 15,814,368.8 | 16,537,477 | 13 | 28 |
| *Helleia helle* | 51.64% | 47.19% | 35.7 | 547,306,268 | 37.76 | 25 | 25 | 0 | 0 | 0 | 0.000 | 0.0 | 34,208,996 | 23,709,742.1 | 23,624,374 | 10 | 23 |
| *Parnassius mnemosyne* | 40.42% | 35.41% | 186.6 | 1,494,369,942 | 38.04 | 261 | 261 | 0 | 0 | 0 | 0.000 | 0.0 | 57,533,117 | 23,729,140.7 | 24,417,079 | 22 | 103 |
| *Spongipellis delectans* | NA | NA | 383.9 | 30,870,404 | 48.06 | 11 | 14 | 3 | 97 | 246 | 0.001 | 166.9 | 4,107,596 | 2,982,307.8 | 3,054,357 | 5 | 11 |
| *Botryllus schlosseri* | 21.52% | 17.81% | 51.5 | 704,149,733 | 40.42 | 1,632 | 1,632 | 0 | 0 | 0 | 0.000 | 0.0 | 6,582,172 | 1,092,712.8 | 772,554 | 250 | 1,185 |
| *Gordionus montsenyensis* | 70.44% | 64.80% | 59.8 | 287,795,482 | 26.93 | 398 | 558 | 160 | 556 | 3,228 | 0.001 | 105.2 | 64,441,940 | 47,960,667.3 | 52,653,292 | 3 | 6 |
| *Phakellia ventilabrum* | NA | NA | 40.0 | 188,607,397 | 43.70 | 83 | 226 | 143 | 758 | 14,300 | 0.008 | 100.0 | 17,765,667 | 7,899,171.6 | 7,068,862 | 11 | 25 |
| *Phaeosaccion multiseriatum* | 30.75% | 25.13% | 598.8 | 67,144,605 | 58.38 | 22 | 25 | 3 | 45 | 300 | 0.000 | 100.0 | 7,404,487 | 4,547,222.2 | 3,942,975 | 7 | 15 |
| *Alburnus alburnus* | 15.42% | 10.74% | 52.9 | 1,005,737,902 | 38.64 | 112 | 332 | 220 | 219 | 87,380 | 0.009 | 437.9 | 59,056,062 | 41,092,524.7 | 38,752,591 | 11 | 24 |
| *Ammodytes marinus* | 30.21% | 24.88% | 29.3 | 777,835,610 | 41.22 | 617 | 1,634 | 1,017 | 1,307 | 203,400 | 0.026 | 200.0 | 39,742,375 | 30,198,617.4 | 31,846,471 | 12 | 24 |
| *Anaecypris hispanica* | 28.88% | 21.32% | 50.1 | 983,738,696 | 38.78 | 52 | 516 | 464 | 472 | 191,962 | 0.020 | 449.9 | 57,885,559 | 40,477,428.4 | 38,642,748 | 11 | 24 |
| *Argentina silus* | 18.33% | 16.88% | 42.4 | 672,210,335 | 44.75 | 1,055 | 2,776 | 1,721 | 2,560 | 344,200 | 0.051 | 200.0 | 66,768,702 | 25,083,908.6 | 23,520,047 | 11 | 77 |
| *Knipowitschia panizzae* | 27.44% | 21.71% | 10.5 | 871,098,461 | 42.57 | 1,388 | 3,148 | 1,760 | 2,020 | 352,000 | 0.040 | 200.0 | 50,443,959 | 31,636,443.5 | 34,494,022 | 12 | 193 |
| *Perca* sp. 'yellow fin Alpine' | 48.06% | 43.04% | 16.6 | 1,012,652,072 | 41.18 | 257 | 606 | 349 | 345 | 69,800 | 0.007 | 200.0 | 50,815,995 | 40,988,722.5 | 41,708,117 | 12 | 23 |
| *Salvelinus alpinus* | 32.29% | 19.10% | 33.3 | 2,399,975,417 | 43.20 | 1,791 | 4,793 | 3,002 | 1,251 | 600,400 | 0.025 | 200.0 | 125,110,484 | 51,575,364.1 | 46,829,268 | 17 | 404 |
| *Silurus aristotelis* | 19.21% | 14.24% | 27.0 | 800,118,513 | 38.90 | 241 | 1,036 | 795 | 994 | 159,000 | 0.020 | 200.0 | 44,060,567 | 29,035,302.9 | 30,578,072 | 12 | 27 |
| *Solea solea* | 11.67% | 9.07% | 16.2 | 645,225,451 | 40.88 | 250 | 613 | 363 | 563 | 72,600 | 0.011 | 200.0 | 47,017,204 | 28,621,223.7 | 29,008,568 | 10 | 22 |
| *Tripterygion tripteronotum* | 61.56% | 49.91% | 24.2 | 772,862,888 | 41.76 | 468 | 1,228 | 760 | 983 | 152,000 | 0.020 | 200.0 | 64,425,221 | 36,667,700.5 | 34,928,193 | 10 | 21 |
| *Zingel asper* | 47.60% | 43.32% | 46.1 | 1,014,494,583 | 41.09 | 1,202 | 1,912 | 710 | 700 | 142,000 | 0.014 | 200.0 | 47,957,273 | 34,799,441.0 | 38,808,765 | 12 | 70 |
| *Haliaeetus albicilla* | 40.26% | 34.78% | 30.3 | 1,320,298,205 | 43.15 | 188 | 612 | 424 | 321 | 84,800 | 0.006 | 200.0 | 85,729,264 | 46,840,544.8 | 44,398,590 | 12 | 30 |
| *Oenanthe leucura* | 66.05% | 58.04% | 21.6 | 1,254,332,796 | 43.33 | 840 | 1,219 | 379 | 302 | 72,791 | 0.006 | 199.1 | 150,389,447 | 60,381,493.1 | 60,889,355 | 7 | 310 |
| *Tetrao urogallus* | 47.09% | 39.66% | 11.6 | 1,013,338,082 | 41.49 | 343 | 1,291 | 948 | 936 | 189,600 | 0.019 | 200.0 | 193,500,925 | 83,780,028.7 | 71,401,156 | 5 | 21 |
| *Canis aureus* | 56.76% | 49.20% | 14.4 | 2,456,326,569 | 41.49 | 109 | 371 | 262 | 107 | 52,400 | 0.002 | 200.0 | 124,578,940 | 68,562,753.7 | 65,319,744 | 15 | 37 |
| *Chionomys nivalis* | 52.13% | 44.02% | 13.2 | 2,393,406,913 | 41.98 | 161 | 358 | 197 | 82 | 39,400 | 0.002 | 200.0 | 205,897,943 | 98,950,611.9 | 88,844,950 | 10 | 26 |
| *Lepus granatensis* | 69.89% | 59.54% | 17.7 | 2,601,373,884 | 44.14 | 191 | 412 | 221 | 85 | 42,076 | 0.002 | 198.8 | 186,673,286 | 122,776,620. | 125,842,53 | 9 | 22 |
| *Lepus timidus* | 47.39% | 44.71% | 11.1 | 2,964,172,812 | 44.03 | 771 | 1,156 | 385 | 130 | 77,000 | 0.003 | 200.0 | 184,151,585 | 115,052,459. | 123,691,39 | 10 | 55 |
| *Mustela lutreola* | 45.63% | 36.34% | 45.2 | 2,467,121,698 | 41.54 | 34 | 204 | 170 | 69 | 13,513,522 | 0.548 | 429,898.1 | 282,538,888 | 158,415,938. | 148,796,03 | 7 | 17 |
| *Vipera ursinii* | 55.56% | 50.42% | 14.2 | 1,625,023,540 | 40.93 | 384 | 2,203 | 1,819 | 1,119 | 363,800 | 0.022 | 200.0 | 359,753,992 | 198,305,116. | 212,821,32 | 3 | 14 |
| *Hottonia palustris* | 49.41% | 44.23% | 59.6 | 790,186,607 | 35.95 | 650 | 657 | 7 | 9 | 3,500 | 0.000 | 500.0 | 108,788,391 | 62,068,300.3 | 63,854,627 | 6 | 15 |
| *Prunus brigantina* | 54.88% | 47.05% | 127.9 | 259,721,188 | 38.25 | 69 | 107 | 38 | 146 | 3,800 | 0.001 | 100.0 | 50,927,966 | 31,413,653.8 | 29,256,389 | 4 | 8 |
| *Trifolium dubium* | 49.09% | 41.53% | 36.9 | 679,499,717 | 32.66 | 156 | 742 | 586 | 862 | 117,200 | 0.017 | 200.0 | 64,644,275 | 46,988,571.2 | 46,006,535 | 7 | 15 |

| Scientific name | Longest_Cor | Cont_auN | Cont_N50 | Cont_L50 | Cont_ | QV | Completeness | BUSCOdb_recen | BUSCOs_recent | BUSCOd_recent | BUSCOf_recent | BUSCOm_recent | BUSCOdb_ancien | BUSCOs_ancient | BUSCOd_ancient | BUSCOf_ancient | BUSCOm_ancien | BUSCOdb_euka | BUSCOs_euka | BUSCOd_euka | BUSCOf_euka | BUSCOm_euka |
|---|---|---|---|---|---|---|---|---|---|---|---|---|---|---|---|---|---|---|---|---|---|---|
| *Andrena humilis* | 1,175,059 | 314,521.1 | 245,000 | 326 | 1,744 | 58.1 | 98.46 | hymenoptera | 95.70% | 0.50% | 0.70% | 3.10% | arthropoda | 98.20% | 0.60% | 0.80% | 0.40% | eukaryota | 96.90% | 0.80% | 1.60% | 0.70% |
| *Osmia cornuta* | 13,604,000 | 4,067,934.6 | 1,242,809 | 53 | 623 | 32.7 | 93.38 | hymenoptera | 96.40% | 1.10% | 0.40% | 2.10% | arthropoda | 98.90% | 0.70% | 0.20% | 0.20% | eukaryota | 97.30% | 2.00% | 0.80% | 0.10% |
| *Carabus granulatus* | 1,465,338 | 171,552.5 | 94,774 | 429 | 2,388 | 51.4 | 84.98 | insecta | 92.70% | 0.90% | 0.50% | 5.90% | arthropoda | 91.70% | 1.20% | 0.90% | 6.20% | eukaryota | 91.80% | 0.40% | 1.20% | 6.60% |
| *Carabus intricatus* | 1,993,519 | 653,602.0 | 459,766 | 121 | 738 | 53.9 | 90.42 | insecta | 98.00% | 0.70% | 0.10% | 1.20% | arthropoda | 98.00% | 1.10% | 0.20% | 0.70% | eukaryota | 99.60% | 0.00% | 0.00% | 0.40% |
| *Epeorus assimilis* | 2,791,144 | 703,580.1 | 544,464 | 216 | 1,216 | 51.8 | 88.77 | insecta | 94.80% | 1.80% | 1.70% | 1.70% | arthropoda | 96.90% | 1.20% | 1.00% | 0.90% | eukaryota | 94.10% | 1.60% | 2.70% | 1.60% |
| *Leptodirus hochenwarti* | 16,373,000 | 5,045,367.8 | 3,374,168 | 37 | 225 | 66.1 | 91.14 | insecta | 98.60% | 0.30% | 0.70% | 0.40% | arthropoda | 98.80% | 0.20% | 0.60% | 0.40% | eukaryota | 97.60% | 1.20% | 0.80% | 0.40% |
| *Stylops ater* | 5,437,292 | 2,346,880.5 | 1,867,595 | 8 | 37 | 26.1 | 26.86 | insecta | 83.00% | 1.50% | 4.10% | 11.40% | arthropoda | 84.30% | 1.50% | 4.40% | 9.80% | eukaryota | 91.40% | 0.00% | 4.70% | 3.90% |
| *Coenonympha glycerion* | 20,120,474 | 11,014,122. | 11,561,668 | 15 | 56 | 44.1 | 71.90 | lepidoptera | 97.90% | 0.60% | 0.40% | 1.10% | arthropoda | 97.60% | 0.90% | 0.70% | 0.80% | eukaryota | 97.30% | 0.40% | 1.60% | 0.70% |
| *Helleia helle* | 34,208,996 | 23,709,742. | 23,624,374 | 10 | 23 | 46.1 | 93.35 | lepidoptera | 98.00% | 0.30% | 0.30% | 1.40% | arthropoda | 98.60% | 0.50% | 0.50% | 0.40% | eukaryota | 98.80% | 0.00% | 1.20% | 0.00% |
| *Parnassius mnemosyne* | 57,533,117 | 23,729,140. | 24,417,079 | 22 | 103 | 15.4 | 89.54 | lepidoptera | 97.70% | 1.30% | 0.20% | 0.80% | arthropoda | 97.80% | 1.70% | 0.40% | 0.10% | eukaryota | 98.80% | 0.00% | 0.80% | 0.40% |
| *Spongipellis delectans* | 4,107,596 | 2,828,109.8 | 2,680,827 | 5 | 11 | 66.9 | 74.49 | polyporales | 93.60% | 0.60% | 1.00% | 4.80% | fungi | 97.90% | 0.40% | 0.30% | 1.40% | eukaryota | 98.00% | 0.40% | 0.80% | 0.80% |
| *Botryllus schlosseri* | 6,582,172 | 1,092,712.8 | 772,554 | 250 | 1,185 | 21.0 | 50.12 | metazoa | 75.90% | 15.30% | 3.80% | 5.00% | metazoa | 75.90% | 15.30% | 3.80% | 5.00% | eukaryota | 80.80% | 15.70% | 2.70% | 0.80% |
| *Gordionus montsenyensis* | 34,589,484 | 10,364,727. | 6,141,318 | 11 | 96 | 36.9 | 84.98 | metazoa | 59.20% | 0.30% | 12.10% | 28.40% | metazoa | 59.20% | 0.30% | 12.10% | 28.40% | eukaryota | 66.30% | 0.00% | 20.80% | 12.90% |
| *Phakellia ventilabrum* | 5,977,719 | 2,325,625.0 | 2,199,000 | 30 | 133 | 32.0 | 65.29 | metazoa | 78.20% | 0.70% | 9.00% | 12.10% | metazoa | 78.20% | 0.70% | 9.00% | 12.10% | eukaryota | 87.50% | 0.00% | 7.10% | 5.40% |
| *Phaeosaccion multiseriatum* | 7,404,487 | 4,320,244.5 | 3,942,975 | 7 | 16 | 46.1 | 70.45 | stramenopiles | 89.00% | 1.00% | 6.00% | 4.00% | stramenopiles | 89.00% | 1.00% | 6.00% | 4.00% | eukaryota | 70.60% | 0.40% | 16.50% | 12.50% |
| *Alburnus alburnus* | 35,110,498 | 9,924,716.2 | 7,667,869 | 39 | 172 | 61.9 | 76.95 | actinopterygii | 95.20% | 1.30% | 0.90% | 2.60% | vertebrata | 96.60% | 0.70% | 0.40% | 2.30% | eukaryota | 97.30% | 2.00% | 0.00% | 0.70% |
| *Ammodytes marinus* | 11,859,408 | 2,499,417.3 | 2,041,615 | 115 | 758 | 55.2 | 85.67 | actinopterygii | 97.10% | 0.90% | 0.30% | 1.70% | vertebrata | 96.80% | 0.70% | 0.90% | 1.60% | eukaryota | 95.70% | 3.10% | 0.00% | 1.20% |
| *Anaecypris hispanica* | 15,099,894 | 4,839,130.3 | 4,199,513 | 75 | 327 | 60.8 | 91.67 | actinopterygii | 95.70% | 1.40% | 0.70% | 2.20% | vertebrata | 96.90% | 0.70% | 0.50% | 1.90% | eukaryota | 97.30% | 1.60% | 0.40% | 0.70% |
| *Argentina silus* | 7,041,626 | 1,430,074.9 | 858,168 | 187 | 1,409 | 54.1 | 92.32 | actinopterygii | 93.50% | 2.60% | 1.60% | 2.30% | vertebrata | 94.80% | 2.50% | 1.00% | 1.70% | eukaryota | 91.80% | 6.30% | 0.00% | 1.90% |
| *Knipowitschia panizzae* | 10,900,401 | 1,437,514.3 | 743,701 | 281 | 1,782 | 25.1 | 83.09 | actinopterygii | 92.80% | 1.00% | 2.00% | 4.20% | vertebrata | 93.50% | 0.80% | 2.30% | 3.40% | eukaryota | 96.50% | 2.00% | 1.20% | 0.30% |
| *Perca* sp. 'yellow fin Alpine' | 32,403,000 | 14,379,344. | 13,498,500 | 24 | 201 | 54.9 | 96.67 | actinopterygii | 97.80% | 0.70% | 0.40% | 1.10% | vertebrata | 97.90% | 0.50% | 0.50% | 1.10% | eukaryota | 97.30% | 2.40% | 0.00% | 0.30% |
| *Salvelinus alpinus* | 9,985,737 | 1,855,422.8 | 1,225,000 | 484 | 2,805 | 56.3 | 91.98 | actinopterygii | 63.30% | 33.30% | 1.00% | 2.40% | vertebrata | 61.40% | 35.80% | 1.00% | 1.80% | eukaryota | 58.00% | 38.40% | 0.80% | 2.80% |
| *Silurus aristotelis* | 19,180,065 | 5,809,939.4 | 4,577,990 | 49 | 406 | 53.6 | 98.01 | actinopterygii | 95.80% | 1.10% | 0.80% | 2.30% | vertebrata | 96.20% | 0.90% | 0.80% | 2.10% | eukaryota | 95.70% | 3.10% | 0.40% | 0.80% |
| *Solea solea* | 11,520,506 | 3,178,823.7 | 2,593,403 | 76 | 307 | 61.8 | 85.51 | actinopterygii | 97.10% | 1.20% | 0.50% | 1.20% | vertebrata | 97.70% | 0.70% | 0.50% | 1.10% | eukaryota | 97.60% | 1.20% | 0.40% | 0.80% |
| *Tripterygion tripteronotum* | 15,170,232 | 4,082,332.9 | 2,804,349 | 71 | 491 | 29.9 | 81.50 | actinopterygii | 96.00% | 0.90% | 1.00% | 2.10% | vertebrata | 96.60% | 0.60% | 1.00% | 1.80% | eukaryota | 97.30% | 1.60% | 1.20% | 0.10% |
| *Zingel asper* | 17,717,955 | 4,079,789.6 | 3,102,298 | 96 | 595 | 55.7 | 95.71 | actinopterygii | 97.50% | 0.80% | 0.50% | 1.20% | vertebrata | 97.50% | 0.90% | 0.70% | 0.90% | eukaryota | 93.30% | 4.70% | 1.20% | 0.80% |
| *Haliaeetus albicilla* | 18,990,351 | 5,706,164.9 | 4,576,000 | 85 | 357 | 70.0 | 90.32 | aves | 96.60% | 0.60% | 0.60% | 2.20% | vertebrata | 95.70% | 0.80% | 1.50% | 2.00% | eukaryota | 96.50% | 1.20% | 1.60% | 0.70% |
| *Oenanthe leucura* | 50,893,993 | 14,558,428. | 10,025,093 | 28 | 566 | 61.2 | 96.53 | aves | 96.80% | 0.30% | 0.60% | 2.30% | vertebrata | 95.70% | 0.60% | 1.20% | 2.50% | eukaryota | 97.60% | 1.60% | 0.80% | 0.00% |
| *Tetrao urogallus* | 10,141,983 | 2,456,608.6 | 1,909,808 | 152 | 687 | 60.3 | 97.73 | aves | 96.30% | 0.30% | 0.60% | 2.80% | vertebrata | 95.10% | 0.50% | 1.00% | 3.40% | eukaryota | 97.30% | 1.20% | 0.80% | 0.70% |
| *Canis aureus* | 62,284,301 | 23,263,959. | 21,906,000 | 37 | 171 | 64.2 | 96.92 | mammalia | 94.00% | 1.90% | 1.00% | 3.10% | vertebrata | 94.20% | 3.60% | 0.90% | 1.30% | eukaryota | 85.90% | 11.40% | 2.00% | 0.70% |
| *Chionomys nivalis* | 78,579,696 | 33,429,891.6 | 30,960,136 | 25 | 139 | 62.2 | 93.67 | mammalia | 94.10% | 2.50% | 0.70% | 2.70% | vertebrata | 94.30% | 4.10% | 0.60% | 1.00% | eukaryota | 89.00% | 9.80% | 1.20% | 0.00% |
| *Lepus granatensis* | 110,458,692 | 43,855,375. | 39,429,213 | 21 | 130 | 65.3 | 89.11 | mammalia | 91.90% | 2.30% | 0.70% | 5.10% | vertebrata | 92.90% | 3.50% | 0.90% | 2.70% | eukaryota | 89.80% | 9.00% | 0.40% | 0.80% |
| *Lepus timidus* | 141,329,000 | 48,340,981. | 43,409,466 | 21 | 297 | 62.7 | 90.06 | mammalia | 93.10% | 3.30% | 0.70% | 2.90% | vertebrata | 93.20% | 4.80% | 1.00% | 1.00% | eukaryota | 87.80% | 11.40% | 0.80% | 0.00% |
| *Mustela lutreola* | 167,065,337 | 56,150,510. | 41,193,119 | 18 | 70 | 47.8 | 98.97 | mammalia | 95.20% | 1.60% | 0.70% | 2.50% | vertebrata | 95.70% | 2.30% | 0.90% | 1.10% | eukaryota | 93.30% | 5.90% | 0.80% | 0.00% |
| *Vipera ursinii* | 12,662,354 | 2,709,639.1 | 2,137,891 | 215 | 1,177 | 58.6 | 95.26 | sauropsida | 91.10% | 1.50% | 1.00% | 6.40% | vertebrata | 95.20% | 1.10% | 1.20% | 2.50% | eukaryota | 98.00% | 1.60% | 0.40% | 0.00% |
| *Hottonia palustris* | 97,747,157 | 46,390,374. | 43,471,964 | 7 | 22 | 61.1 | 97.50 | embryophyta | 94.90% | 2.70% | 1.10% | 1.30% | viridiplantae | 95.80% | 3.50% | 0.50% | 0.20% | eukaryota | 85.50% | 14.10% | 0.00% | 0.40% |
| *Prunus brigantina* | 26,391,212 | 12,923,413.1 | 12,745,445 | 8 | 35 | 34.0 | 97.28 | embryophyta | 96.60% | 2.20% | 0.80% | 0.40% | viridiplantae | 95.80% | 2.80% | 0.90% | 0.50% | eukaryota | 89.40% | 10.20% | 0.40% | 0.00% |
| *Trifolium dubium* | 10,329,705 | 3,277,282.8 | 3,006,263 | 75 | 283 | 69.1 | 98.66 | embryophyta | 5.80% | 93.70% | 0.10% | 0.40% | viridiplantae | 6.10% | 93.90% | 0.00% | 0.00% | eukaryota | 5.90% | 94.10% | 0.00% | 0.00% |
