## Supplementary_Information for "The European Reference Genome Atlas: piloting a decentralised approach to equitable biodiversity genomics"

### Supplementary Tables

**Supplementary Table 1: ERGA Sequencing Centre Partners, Biobanks and Museums Collection and Locations**

| Sequencing Centres | Location |
|---|---|
| GIGA-Genomics University Liege | Belgium |
| VIB-University of Antwerp | Belgium |
| Université libre de Bruxelles (ULB) | Belgium |
| University of Copenhagen | Denmark |
| Genoscope - Centre National de Séquençage | France |
| West German Genome Center | Germany |
| Dresden Max Planck Institute of Molecular Cell Biology and Genetics (MPI | Germany |
| NCCT (NGS Competence Center Tübingen) | Germany |
| Hungarian Centre for Genomics and Bioinformatics, University of Pécs | Hungary |
| University of Bari and Consiglio Nazionale delle Ricerche | Italy |
| University of Florence | Italy |
| Marine Animal Ecology / Animal Breeding and Genomics (Wageningen University) | Netherlands |
| Norwegian Sequencing center | Norway |
| Centro Nacional de Análisis Genómico (CNAG) | Spain |
| SciLifeLab | Sweden |
| University of Bern, Next Generation Sequencing (NGS) Platform | Switzerland |
| Functional Genomic Centre Zurich | Switzerland |

| | |
|---|---|
| Genomic Technologies Facility Lausanne | Switzerland |
| Wellcome Sanger Institute | UK |
| Earlham Institute | UK |

**Supplementary Table 2: Roles and Responsibilities of Genome Team Members**

| **Team Member** | **Role** |
|---|---|
| Principal Investigator | Each genome has a designated Principal Investigator (PI) who is responsible to ensure the coordination of the project. If a sample ambassador specifically requests to lead the species genome analysis, this person will become the PI for this species' genome. Participants covering the costs for the genomes can also request to be PI or co-PIs, in which case the sample ambassador must be in agreement. In case the participants that are covering sequencing costs are not based in the same country as the sample origin, council members from the country of origin must agree to the request. Other participants can also request to become PI or co-PIs. The sample ambassador, in agreement with other potential co-PIs, decides on the request. Co-PIs appoint a coordinator among them responsible for the project, along with organizing the handling of the samples, to ensure that proper documentation meeting the Nagoya Protocol and local regulations is provided. |
| Sample collector and/or sample provider | Responsible for the ethical and legal collection of samples for ERGA and compliance with ERGA's 'Sample Collection Code of Best Practices' and 'Data Sharing and Management Policy'. |
| Taxonomist and/or ex-situ sample manager | Individual(s) responsible for ensuring the taxonomic validity of the sample obtained and its deposition into an appropriate biobank or collection understanding that the PUID associated with both will be disclosed in the ERGA metadata manifest. |
| Sample ambassador | The sample ambassador coordinates and organizes all of the samples, permits, barcoding, and storage components of the project, up to and including the shipment to laboratory and storage of vouchers. The individual (s) should be based in the country of origin or have an established research project in the area of sampling that justifies the involvement in genome establishment for species from another country. If not based in the country, s/he must provide proof of compliance with CBD Nagoya protocol as well as permits for in-country sampling, |

| | |
|---|---|
| | sample import/export and handling. The sample ambassador can also act as the sample provider. |
| Wet-lab processor | Facilitates all wet lab components of the project including HMW DNA extraction, library preps, sequencing. This is a hands-<br>on researcher(s), or group, or could even be a facility PI. |
| Genome assembly manager | The individual(s) responsible overseeing the development, optimization, and correction of the genome assembly. Role also ensures appropriate computational resources are available. |
| Genome assembly and/or curation generator | The individual(s) responsible for the generation of the genome assembly (hands-on role). |
| Genome annotation and/or genome analysis generator | The individual(s) responsible for the generation of the gene annotation and data analysis (hands-on role). |

**Supplementary Table 3: OmniC sequencing statistics from ERGA Sequencing Hub. Estimated genome sizes for *C. barii* and *T. fluviatilis* were obtained from Genomes on a Tree (GoaT).**

| | Species | Tissue | % GC | Average read length (bp) | Read pairs (millions) | Unique read pairs (millions) | Genome size (Gbp) | Coverage |
|---|---|---|---|---|---|---|---|---|
| *1* | ***Botryllus schlosseri*** (Golden Star Tunicate) | whole specimen | **42%** | **150** | **282.6** | **155.1** | **0.70** | **66** |
| *2* | ***Cryptocephalus barii*** | whole specimen | **43%** | **150** | **305.3** | **180.6** | **0.50** | **108** |
| *3* | ***Hottonia palustris*** (water violet) | whole specimen | **37%** | **151** | **129.6** | **108.5** | **0.86** | **38** |
| *4* | ***Knipowitschia panizzae*** (Adriatic dwarf goby) | gills | **45%** | **150** | **104.4** | **88.9** | **0.87** | **31** |
| *5* | ***Lepus granatensis*** (Granada hare) | kidney | **45%** | **150** | **578.1** | **443.4** | **2.65** | **50** |

| | | | | | | | | |
|---|---|---|---|---|---|---|---|---|
| *6* | ***Mullus barbatus*** (red mullet) | liver | **47%** | **150** | **32.8** | **30.4** | **0.54** | **17** |
| *7* | ***Oenanthe leucura*** (Black wheatear) | blood | **44%** | **150** | **339.3** | **252.4** | **1.36** | **56** |
| *8* | ***Parnassius mnemosyne*** (Clouded Apollo) | whole specimen | **40%** | **151** | **29.1** | **25.3** | **1.46** | **5** |
| *9* | ***Salvelinus alpinus*** (Arctic char) | liver | **45%** | **150** | **386.4** | **275.5** | **2.73** | **30** |
| *10* | ***Theodoxus fluviatilis*** (river nerite) | whole specimen | **48%** | **150** | **321.2** | **177.1** | **0.86** | **62** |
| *11* | ***Tripterygion tripteronotum*** (red-black triplefin) | brain | **44%** | **149** | **236.2** | **204.3** | **0.77** | **79** |

**Supplementary Table 4: RNA-seq statistics from ERGA Sequencing Hub**

| Species | No. of libraries | Tissues | % Duplications | % GC | Average read length (bp) | Read pairs (millions) |
|---|---|---|---|---|---|---|
| ***Asellus aquaticus*** (water hoglouse) | 1 | whole specimen | **81%** | **39%** | **141** | **38** |
| ***Acanthodactylus schreiberi*** (Schreiber's fringe-fingered lizard) | 4 | brain, kidney, liver, muscle | **51%** | **47%** | **136** | **124** |
| ***Andrena vaga*** (grey-backed mining bee) | 1 | whole specimen | **63%** | **46%** | **132** | **119** |
| ***Bufotes viridis*** (European green toad) | 7 | heart, kidney, liver, lung, muscle, spleen, skin | **55%** | **46%** | **136** | **217** |

| | | | | | | |
|---|---|---|---|---|---|---|
| ***Corema album*** (Portuguese crowberry) | 1 | leaf | **77%** | **52%** | **88** | **4** |
| ***Cryptocephalus barii*** | 2 | whole specimens | **78%** | **43%** | **132** | **277** |
| ***Hottonia palustris*** (water violet) | 4 | old buds, young buds, leaves, flowers/petals | **65%** | **48%** | **138** | **94** |
| ***Laurus azorica*** (the Azores laurel) | 1 | leaf | **60%** | **52%** | **121** | **6** |
| ***Lepus granatensis*** (Granada hare) | 5 | kidney, liver, lung, spleen, testes | **50%** | **54%** | **136** | **179** |
| ***Mullus barbatus*** (red mullet) | 4 | fin, gonad, kidney, muscle | **60%** | **52%** | **126** | **262** |
| ***Nepa anophthalma*** (Stygobiotic Waterscorpion) | 1 | whole specimen | **81%** | **36%** | **140** | **53** |
| ***Oenanthe leucura*** (Black wheatear) | 1 | blood | **77%** | **54%** | **128** | **93** |
| ***Palingenia longicauda*** (Tisza mayfly) | 2 | larvae | **90%** | **39%** | **140** | **75** |
| ***Parnassius mnemosyne*** (Clouded Apollo) | 1 | whole specimen | **76%** | **42%** | **138** | **94** |
| ***Spinachia spinachia*** (Fifteen-spined stickleback) | 4 | pelvic fin and muscle, internal organs, brain and eyes, gills | **51%** | **51%** | **133** | **171** |
| ***Stylops ater*** | 2 | whole specimens | **70%** | **39%** | **139** | **115** |

| | | | | | | |
|---|---|---|---|---|---|---|
| ***Theodoxus fluviatilis*** (river nerite) | 1 | foot | **74%** | **45%** | **138** | **44** |
| ***Trechus terceiranus*** | 1 | whole specimen | **73%** | **41%** | **142** | **52** |
| ***Zostera noltei*** (dwarf eelgrass) | 1 | leaf | **86%** | **49%** | **135** | **74** |
| ***Zygaena transalpina*** (Transalpine Burnet Moth) | 1 | whole specimen | **62%** | **43%** | **144** | **66** |

#### Supplementary Case Studies

**Case-study 1: Navigating Nagoya Compliance**

Ten genome teams across eight countries and regions (Malta, Azores, Croatia, France, Greece,, Hungary, Portugal, and Slovakia) had to obtain a Nagoya permit prior to collecting samples for the project. For all countries the process for obtaining a permit in a relatively short period of time, averaging two months, was centred on an initial engagement with the Access and Benefit Sharing (ABS) National Focal Point (NFP) or Competent National Authority (CNA). Both NFPs and CNAs offered initial guidelines for the specific national procedures. Some countries have online portals to streamline the Nagoya permitting process e.g, France (https://www.ecologie.gouv.fr/acces-et-partage-des-avantages-decoulant-lutilisation-des-ressources-genetiques-et-des-connaissances), Azores (https://servicos-sraa.azores.gov.pt/doit/servicos.asp?id_dep=3&id_form=18) and Malta (https://www.servizz.gov.mt/en/Pages/Environment_-Energy_-Agriculture-and-Fisheries/Agriculture/Agriculture/WEB05310/default.aspx). Other countries had easily accessible downloadable application templates, whilst others coordinated permitting through email engagements with the National Focal Point (Azores) being asked to provide information regarding: sampling, uses, intent to transfer and confirming they would cooperate with any knowledge transfer or benefit sharing obligations.

Most sample providers declared non-commercial use (research purpose) for the sample collection purposes. The terms and conditions of the IRCCs obtained were diverse and ranged in complexity. For Azores, the IRCC laid out provisions for sampling, reporting, benefit-sharing and third party transfers. In terms of benefit-sharing, the Azorean team ensures the disclosure of the IRCC permit ID in all scientific publications, a key form of non-monetary benefit sharing[1]. For France, just sampling details were required and the initial permit has to be updated if a change in utilisation occurs (further biochemical analyses, commercial uses etc…). For Croatia, a short report on the samples collected after the duration of the permit was requested along with the collection methodology and number of specimens collected.

For Malta, terms and conditions were more comprehensive including: notification if a change in contract terms was required; compliance with local authorities; respectful use of Traditional Knowledge if relevant; a summary report on project conclusion; copy of all associated research publications (must be made freely available and attribute the sample provider country); annual progress reports; documentation storage for 20 months after project completion; inform local authorities when data becomes publicly available; and compliance with Europe's Due Diligence requirements through 'DECLARE'. For Hungary, the initial permit had been previously acquired, outlining the terms for collection of *Vipera ursinii rakosiensis* samples to facilitate a genetic screening for a reintroduction program of this endangered species. For the purposes of ERGA and sample sequencing in the UK, the sample ambassador had to obtain a new permit in order to send specimens outside of the EU. The permitting process was free of cost for most teams apart from Hungary (20,000 HUF). Four genome teams also required CITES permits.

**Case-study 2: Sample Collection and DNA extraction barriers**

Criteria for inclusion were developed for ERGA to prioritise species where genomic data would have immediate potential impacts e.g., endangered species. However, to ensure feasibility, more straightforward species were also prioritised, e.g., haploid, <~1Gb genome size etc.. In practice however, acquiring suitable samples presented challenges for some species, for instance in the case of the threatened Siberian flying squirrel (*Pteromys volans*) and the golden jackal (*Canis aureus*). Due to the threatened status of the flying squirrel, only pre-existing samples could be obtained. These initial samples belonged to found-dead animal, however during DNA extraction the samples yielded insufficient levels of high molecular weight DNA (HMW-DNA). After this, a sample from an ear biopsy from a live animal was obtained, but again yielded inadequate HMW-DNA. Similarly for the jackal, two samples taken from dead animals and stored at -80 had inadequate HMW-DNA volumes. The Rhône Streber, one of the rarest and most strongly threatened fish species of Europe, also highlighted the complexity of sampling endangered species. Over the past decade, this species has declined sharply both in Switzerland and France and lately has fallen below the limit of detection in Switzerland. Obtaining samples from the wild population was not permitted however, fortunately a single individual was provided for the purposes of the ERGA from an international ex-situ breeding program at Aquatis Aquarium Vivarium Lausanne.

Interestingly, in a few cases (*Lepus timidus* & *Lepus Europaeus)*, reference genomes were successfully produced from HMW-DNA extracted from fibroblast cell lines. This technique did not require invasive sampling or large tissue volumes. Utilizing ex-vivo cell lines could provide a solution for producing reference genomes for a wide range of species where obtaining fresh, flash frozen samples may be a challenge, particularly endangered or protected species, as it results in minimal harm to the individual or population.

Intentionally prioritising species with smaller genome sizes resulted in additional challenges. For instance, for the reference genome for *Stylops ater* (body weight =2mg) [2], DNA from a single individual was ideal and so an amplification step was undertaken. This resulted in the successful production of long reads but failure of HiC data, and so the assembly produced failed to meet the EBP metric as it could not be adequately scaffolded and curated. *Gyrodactylus teuchis*, a monogenean species, was prioritised for inclusion as it would be the first species within the entire Monogenea class of parasitic flatworms to have a reference genome produced. Due to its small size, a single individual yielded only 1ng of DNA and the pooling of worms would introduce unwanted sequence variation into the reference genome. The challenge of DNA extraction and library preparation from single worms was tackled using specific low-input protocols by the team of the Darwin Tree of Life initiative. *G.teuchis* is an obligate parasitic species living and feeding on the skin of its host species, making the extraction of DNA, without contamination from the host species, essentially impossible. Further, the genome of the host (in this case *Salmo trutta*) is about two orders of magnitude larger than the parasites, so even minute contamination with host cells will lead to considerable contamination at the read level. Such reads will be removed rigorously using the reference genome of the host[3] (), as well as in the assembly process through iteratively assessing assemblies and filtering based on aggregate properties, such as coverage and GC content[4]. For *Stylops ater* this was solved by sampling the free-living stage of adult males and not females that never leave the host body[5]. *Cladonia norvegica*, a lichen symbiosis, also yielded low volumes of HMW-DNA, with contamination (at least one fungus, one alga and many bacteria) being problematic for successful DNA extraction. Although sequencing has not yet been completed for this organism, it is highly likely to also contain mite DNA, a regular inhabitant of this symbiosis. Even with tissue free of extracellular contamination, intracellular symbiotic or parasitic bacteria can pose a challenge. An estimated 20% of all insects are infected with Wolbachia[6,7]. Genomes of three different strains of Wolbachia, two complete and one incomplete, could be filtered out and assembled from the long-read data of *Stylops ater*.

Such cases highlight the challenges faced when producing high quality reference genomes from endangered and threatened species as the production of genomes that meet the EBP metrics require large quantities of HMW-DNA that can more likely be obtained by freshly collected and flash-frozen samples. It also showcases the need for adequately training biodiversity researchers new to the field of reference genome production, prior to sample collection, to understand the importance of sampling methodology, specifically in terms of tissue type, tissue quality, preservation method and storage. For 48% of the teams participating, it was the first experience in producing high quality reference genomes and therefore, many were inexperienced in the community accepted best practices for sampling. Without sufficient training prior to sample collection, the likelihood of samples of suboptimal quality being sent for sequencing is much greater, resulting in the wasting of time-, financial- and personnel resources.

**Case-study 3: Swiss Hi-C Protocol Challenges**

Attempts at generating sufficient Hi-C data for the two bees (*Andrena humilis* and *Osmia cornuta*), two beetles (*Carabus intricatus* and *C. granulatus*), and the mayfly (*Epeorus assimilis*) were ultimately unsuccessful. The first trial was performed using *A. humilis* thorax tissue following the ProximoTM Hi-C Kit (Animal) Protocol v4.0 from Phase Genomics. This yielded 27 Gbp from 90M reads, however, contamination checks revealed 75% of reads mapping to *Pseudomonas* bacteria leaving insufficient read coverage for scaffolding. For the second set using the same protocol, head tissues were used for the bees and the mayfly while leg tissue was used for the beetles. Sequencing yielded variable read counts of 290M *A. humilis*, 23M *O. cornuta*, 212M *C. intricatus*, 275M *C. granulatus*, 100M *E. assimilis*, however after deduplication there remained only 14%, 5%, 29%, 24%, and 6% of reads, respectively. The high levels of duplicates observed resulted from performing more than the maximum recommended number of PCR cycles during amplification steps in an attempt to increase overall yield. The low numbers of unique and mappable reads meant that the read coverage obtained for each of the five species was not sufficient to use for scaffolding the primary assemblies. Resampling is now underway (*E. assimilis* and *C. granulatus* collected, others ongoing) to collect new individuals from which to obtain samples.

**Case-study 4: Situating ERGA inside the global biodiversity community**

As part of its commitment to biodiversity research, ERGA is keenly aware of the importance of preserving biodiversity hotspots and unique ecosystems, and strives to be involved in conservation campaigns to protect them. An example of such a campaign is the one to protect Ayyalon Cave in Israel, a unique isolated ecosystem based solely on chemoautotrophic food production by sulfur-oxidising microorganisms[8]. The cave was discovered inside an active quarry in central Israel in 2006. It has probably been isolated from the surface for as long as six million years [9]. Nearly all of the species found in the cave were new to science and are endemic to this ecosystem. The cave and its specialised fauna first came to ERGA's attention when a suggestion was advanced to sequence the genomes of some of its unique species as part of the pilot project. Shortly afterwards it emerged that the very existence of the Ayyalon Cave ecosystem was under threat due to a planned project to use the quarry in which the cave resides as an overflow reservoir for flood management. Allowing flood waters into the quarry would almost definitely inundate the cave, disrupting its unique food web, and would lead to the extinction of its endemic fauna.

A group of Israeli scientists and conservationists rapidly mobilised to oppose this plan. They organised a public campaign with a series of online petitions on various platforms, wrote professional letters to relevant government agencies and decision makers, and appealed to the international scientific community to provide letters of support for the protection of Ayyalon Cave [10]. ERGA was one of the international organisations and societies providing letters of support. Ultimately, the public campaign and international support were successful[10], the flood management plan was modified so as to not include Ayyalon quarry, and the cave was saved. This story highlights the role that biodiversity genomics initiatives can play not only in the effort to document biodiversity but also in the never-ending struggle to preserve it.

**Case-study 5: Accessibility of Cold Chain Shipment**

Sample quality and DNA integrity are essential for the extraction of HMW DNA, which in turn is essential for the production of complete reference genomes that rely on long-read data. To this end, sample collection, preservation and storage are key to the successful production of high-quality reference genomes. To increase the likelihood of success, and in accordance with community-accepted best practices, ERGA endorsed all samples to be ethically and legally sourced, immediately flash frozen, and stored at -80ºC.

For sustaining sample integrity during shipment to ERGA-Pilot associated sequencing facilities, shipment on a continuous cold chain using dry ice was preferred. In 2022, there was a global shortage of dry ice due to the rising cost of gas and other factors that greatly impacted ERGA-Pilot causing weeks of delays in shipping for some teams. Interestingly, 41% of teams (n=93) experienced a challenge during shipment, and 43% required additional samples to be sent. Most reported insufficient sample quality or DNA quantity as the main reason for reshipment. Several reported delays in courier shipment as the cause for sample quality degradation.

Almost 50% of teams paid between €100- €500 per shipment, and 34% < €100. However, for many teams that had less genomics experience or from a country/region that was under-resourced regarding genomics, the costs associated and the certifications required for cold chain shipping were prohibitive and made this an inaccessible option. Highlighting this issue was the sulfidic groundwater aquifer samples collected from the deep recesses of Movile Cave. Here, old 20 m deep hand-dug drinking wells were the only windows of access making sample collection challenging and requiring expertise in single rope techniques to reach the sulfidic sites. Even more challenging was shipping these samples from the cave to the sequencing centre on a cold chain, with the shipment cost estimated at €300 per sample - a prohibitive expense for the genome team. The most viable and cost-effective option for the team was to purchase a roundtrip plane ticket at €80 and travel with the live samples to the sequencing centre.

In other cases, teams experienced a reluctance from couriers to send biological materials on dry ice. Despite declaring on public-facing websites their ability to do so, when contacted their response was either negative or expressed a requirement for a shipment certificate. These required certificates take time to obtain but also have associated costs and would potentially be under-utilised by those sending only a few samples. As a result, it was arranged that it would be more time- and cost-effective to hand deliver some samples. One team travelled 900 km and met halfway with colleagues from the sequencing centre.

Moving forward, ERGA will test alternative and less costly methods for shipment e.g., DNA/RNA Shield for RNAseq samples, to increase the accessibility of the production of reference genomes to all across Europe. ERGA will also consider how to coordinate the shipment of species from a country, weighing up whether it is more time- and cost-effective to first centralise the samples within a country so that a single shipment can be conducted, or alternatively ship samples from multiple locations across the same country. Additionally, as ERGA grows and gains insights into the permitting and certification procedures necessary for shipment across European countries, it could become possible to develop shipment guidelines to support participating researchers or indeed develop partnerships with courier services with centralised ERGA accounts to streamline the process and potentially obtain discounted prices.

**Case-study 6: Experiences from ERGA Library Preparation Hubs**

Having dedicated resources (financial, infrastructural, personnel) to facilitate members from institutions, regions, or countries that are equity deserving in terms of genomics research would greatly expedite the successful utilisation of the infrastructure. For the pilot test, the ERGA Library Preparation and Sequencing Hubs stood at the front-line of tackling equity barriers, and so faced several challenges. One such challenge was obtaining samples that were of a quality and quantity that could support long-read and Hi-C data production [**Supplementary Case-study 2].** Many samples needed to be recollected and reshipped, e.g., *Trechus terceiranus* (icTreTerc1), a cave adapted endemic beetle from the island of Terceira (Azores, Portugal), was resampled due to sample spoilage caused by dry-ice evaporation during cold-chain shipment [see **Supplementary Case-study 5**]. DNA extraction and library construction from recalcitrant species also presented challenges [**Supplementary Case-study 2,3**]. For instance, *Palingenia longicauda* (iePalLong1) has a large cuticle-to-tissue ratio and the presence of large wings interfered with our ability to obtain suitable samples, and *Stylops ate*r (ivStyAter1), an endoparasite of the grey-backed mining bee *Andrena vaga*, failed the library preparation step multiple times due to a limited volume of starting material. Finally, for plant species RNA containing ribosomes from different organelles results in multiple RNA bands making it challenging to accurately analyse the integrity of the RNA. Additionally for both arthropods and molluscs the 28S subunit rRNA is susceptible to a gap deletion that causes band fragmentation. This collapse appears as a single band that resembles the 18S rRNA subunit that can easily be misinterpreted as rRNA degradation with the Rin Integrity Numbers obtained being extremely low[43].
